# Murine CMV infection unmasks macrophage-driven inflammatory cardiomyopathy in *Pkp2*, but not in *Ttn* mutant mice

**DOI:** 10.64898/2025.12.11.693827

**Authors:** Alexandra Cirnu, Tatjana Williams, Mike Nörpel, Julian Kammerer, Svenja Kannt, Mairin Heil, Laura Kimmel, Anahi-Paula Arias-Loza, Manivel Lhoda, Thomas Henning, DiyaaElDin Ashour, Sarah-Lena Puhl, Giuseppe Rizzo, Clément Cochain, Tobias Krammer, Alexander Leipold, Antoine-Emmanuel Saliba, Matthias Mack, Nicole Ziegler, Nancy Ernst, Ralf Joachim Ludwig, Gustavo Campos Ramos, Stefan Frantz, Lars Dölken, Brenda Gerull

## Abstract

2.

**Background:** Genetic cardiomyopathies display variable penetrance and phenotypic expression, highlighting the influence of environmental modulators. Myocarditis, commonly triggered by cardiotropic viruses, overlaps clinically with genetic cardiomyopathies. Consequently, these infections are implicated as secondary factors that accelerate disease onset and progression, yet their precise impact in specific genetic settings remains unexplored.

**Methods:** To interrogate this, genetic mouse models heterozygous for a mutant allele of desmosomal plakophilin-2 (*Pkp2^+/−^*) or sarcomeric titin (*Ttn^+/−^*), genes frequently linked to acute myocarditis, were challenged with murine cytomegalovirus (MCMV) to determine how latent infection influences myocardial inflammation, tissue remodeling, and cardiac performance. Integrated experimental approaches, including echocardiography, histology, flow cytometry, single-cell RNA sequencing, as well as cytokine and kinome analyses, defined immune and signaling responses in infected versus non-infected hearts.

**Results:** Acute, MCMV-induced viral myocarditis and subsequent latent MCMV infection unmasked early disease onset in *Pkp2^+/−^* animals, leading to progressive systolic impairment, whereas in *Ttn^+/−^* mice cardiac structure and function remained preserved throughout infection. Cardiac immune profiling uncovered infection- and genotype-specific divergence: both genetic models showed a stable myocardial effector-memory CD8^+^ T-cell response to MCMV, but only *Pkp2^+/−^* hearts recruited additional Ly6C^+^ CCR2^+^ monocytes and macrophages with distinct inflammatory signatures. In the absence of infection, *Pkp2* insufficiency initiated subclinical CCL2 secretion and subsequent recruitment of CCR2^+^ cells, reflecting early immune activation preceding age-associated functional and structural decline. At this stage, cytokine and kinase evaluations indicated a balance between proinflammatory and compensatory signals. However, with aging or following MCMV challenge, this balance shifted towards persistent inflammation, evidenced by chronic upregulation of cytokines and activation of signaling pathways, which ultimately led to adverse effects and myocardial dysfunction.

**Conclusions:** Manifestation of genetic cardiomyopathies depends on interactions between inherited susceptibility and environmental stressors. Here, we show that cytomegalovirus infection intensifies inflammation in *PKP2*-related cardiomyopathy. In contrast, *TTN*-linked cardiomyopathy does not exhibit increased inflammation under the same conditions. For individuals carrying desmosomal variants, infection control and tailored anti-inflammatory strategies may attenuate or delay disease manifestation and progression.

## 3. Introduction

Clinical presentation and outcomes of patients with cardiomyopathies are determined by a combination of genetic and environmental factors. Among these, viral infections are thought to represent a contributing factor by initiating early cardiac damage and innate immune responses, leading to an acute stage of an inflammatory myopericardial syndrome (IMPS)^1^. Acute myocarditis-like episodes have been reported, especially in patients with arrhythmogenic right ventricular cardiomyopathy/arrhythmogenic cardiomyopathy (ARVC/ACM)^2^. Cardiotropic viruses, such as enterovirus, adenovirus and cytomegalovirus, have been detected in sporadic cases^3^, suggesting a pathogenic link between ACM and viral infection. This has led to the hypothesis that pathogenic variants in desmosomal genes could increase the heart muscle’s susceptibility to viral infections or inflammation. In this context, genetic alterations in desmoplakin (DSP) are frequently associated with myocarditis-like symptoms^2^; however, genetic variants in the most common disease-associated gene, plakophilin-2 (PKP2)^4^, exhibit similar clinical features^5,6^. Heterozygous PKP2 variants show incomplete penetrance, resulting in a broad spectrum of phenotypes, ranging from patients with severe arrhythmias and heart failure to asymptomatic carriers^3^. As a result, the impact of non-genetic factors, especially viral infections, is gaining increasing importance. A recent meta-analysis revealed a high prevalence of pathogenic or likely pathogenic gene variants, notably in the sarcomeric gene titin (TTN) in addition to desmosomal genes (DSP, PKP2, and DSG2) among adults with myocarditis^7^. The presence of these gene variants coupled with acute myocarditis has been linked to a higher risk of adverse cardiovascular events, highlighting an essential connection between inflammation and accelerated disease progression in cardiomyopathies^8^.

Titin truncating variants (TTNtv) are a common cause of dilated cardiomyopathy (DCM), accounting for up to 25% of cases^9^. Several modifying factors, including pregnancy, alcohol abuse, chemotherapy, and other comorbidities, contribute to overt disease^10^. Besides numerous human studies, we previously demonstrated that second hits like angiotensin II or isoproterenol led to the development of cardiomyopathy with cardiac dysfunction and fibrotic remodeling in a mouse model carrying an A-band Ttn truncating variant^11^, however, the role of viral infections remains unknown, even though cardiotropic viruses have been detected in endomyocardial biopsies of DCM patients^12^. Animal models are therefore essential to dissect how secondary triggers interact with genetic susceptibility, shape immune activation and determine the timing of therapeutic interventions. Therefore, we aimed to study the mechanistic interplay among genetic predispositions, viral pathogenesis, and immune responses during infection with the common viral pathogen cytomegalovirus (CMV) in murine models of cardiomyopathy.

Human CMV infections generally remain subclinical in healthy individuals, but have been associated with cardiovascular diseases such as myocarditis. Additionally, seropositivity and lifelong latency are linked to a higher risk of developing inflammatory diseases later in life^13,14^. CMV can infect various cells in peripheral organs, including the heart, triggering typical immune responses involving innate and adaptive immune cells. Murine CMV (MCMV) closely resembles human CMV, mimicking its primary functional pathways, and is therefore used to study the acute and latent effects of viral infections on cardiovascular diseases^15^.

Consequently, we examined the effects of MCMV infection as a second hit in two heterozygous genetic mouse models that mimic *PKP2*- and *TTN*-associated genetic predispositions. Interestingly, we observed distinct outcomes regarding the development of cardiomyopathy and the role of the innate immune system in the disease process. Heterozygous cardiac-restricted *Pkp2^+/−^* mice developed progressive cardiac dysfunction linked to a myeloid cell and cytokine-driven inflammatory response. In contrast, hearts of *Ttn^+/−^* mice remained unaffected despite an appropriate MCMV-associated T-cell response. Furthermore, detailed age- and time-dependent analyses of cardiac immune cell composition, cytokines, and their receptors in *Pkp2^+/−^*mice indicate an early, intrinsic role for the CCL2-CCR2 axis and suggest a potential therapeutic window for targeted inflammatory therapies.

## 4. Methods

All mouse studies and animal numbers used were in compliance with the Directive 2010/63/EU of the European Parliament, approved by the local ethics committee (RUF-55.2.2-2532-2-663, RUF-55.2.2-2532-2-962, and RUF-55.2.2-2532-2-1244) and conducted in accordance with the institutional guidelines.

### Data Availability

A detailed methods section and all data supporting the findings of this study are provided within the article and its Supplemental Material. The RNA sequencing dataset is available from the NCBI Gene Expression Omnibus (GEO) under the GSE311777 accession number.

## 5. Results

### Cardiomyocyte-specific heterozygous loss of *Pkp2* reveals normal myocardial structure, function and immune cell composition

To investigate how loss of one or both alleles of PKP2 affects cardiac structure, function, myocardial inflammation and immune responses in a cardiac-specific manner, we intercrossed *Pkp2* floxed mice with Cre-deleter mice in which the Cre recombinase gene is expressed under the control of the α-myosin heavy chain (αMHC) promoter which yielded homozygous (*Pkp2^−/−^*, Figure 1A; S1A) and heterozygous (*Pkp2^+/−^*) knock-out mice. Homozygous loss of PKP2 protein (Figure S1B+C) did not induce premature mortality, but caused marked biventricular dysfunction at 6-8 weeks of age compared to age-matched control (Ctr) animals (Figure 1B; Table S1). Consistent with this, picrosirius red (PSR) staining revealed significantly increased collagen deposition in both ventricles (Figure 1C). However, flow cytometry showed no differences in total CD45^+^ leukocytes, monocytes/macrophages or infiltrating Ly6C^+^ monocytes compared to Ctr, suggesting no active cell-based immune response at this already advanced stage of disease (Figure 1D).

**Figure 1.**
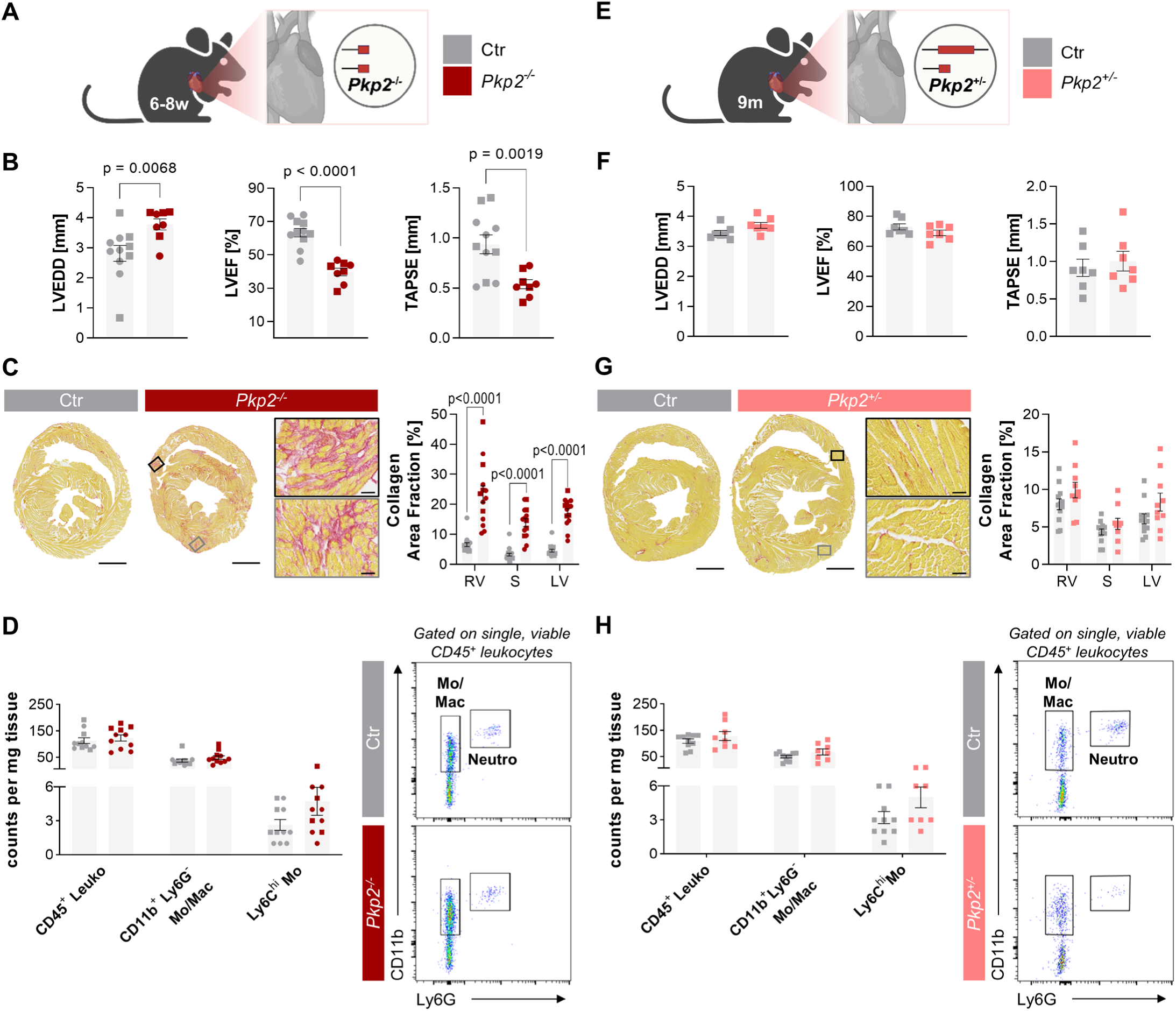
Heterozygous *Pkp2* deficiency does not impact cardiac structure and function, while homozygous loss of *Pkp2* leads to early cardiomyopathy. **A**, Scheme of homozygous cardiac-specific *Pkp2* knockout (*Pkp2^−/−^*) mouse model; signature colors: Ctr – grey, *Pkp2^−/−^* dark red. **B**, Quantification of echocardiographic parameters at 6-8 weeks; Ctr n = 11, *Pkp2^−/−^* n = 8. **C**, Picrosirius red (PSR) stained cardiac cross-sections and quantification of collagen area fraction; scale bars: overview 1000µm, detail 50µm; Ctr n = 17, *Pkp2^−/−^* n = 14. **D**, Quantification of different cardiac CD45^+^ leukocytes by flow cytometry in counts per milligram tissue; representative dot plots showing monocytes/macrophages and neutrophils; Ctr n = 11, *Pkp2^−/−^* n = 11. **E**, Scheme of heterozygous cardiac-specific *Pkp2* knockout (*Pkp2^+/−^*) mouse model; signature colors: Ctr – grey, *Pkp2^+/−^* pink. **F**, Quantification of echocardiographic parameters at 9 months; Ctr n = 7, *Pkp2^+/−^*n = 7. **G**, PSR stained cardiac cross-sections and quantification of collagen area fraction; scale bars: overview 1000µm, detail 50µm; Ctr n = 12, *Pkp2^+/−^* n = 10. **H**, Quantification of different cardiac CD45^+^ leukocytes by flow cytometry in counts per milligram tissue; representative gating showing monocytes/macrophages and neutrophils; Ctr n = 10, *Pkp2^+/−^* n = 8. Data as mean ± SEM. Male: squares; female: circles. Unpaired t test (± Welch’s correction) or Mann–Whitney test, as appropriate. Ctr – control, LVEDD – left ventricular end-diastolic diameter, LVEF – left ventricular ejection fraction, TAPSE – tricuspid annular plane systolic excursion, RV – right ventricle, S – septum, LV – left ventricle, Leuko – leukocytes, Mo/Mac – monocytes/macrophages, Mo – monocytes, Neutro – neutrophils.

Since ACM is inherited in an autosomal dominant manner^16^, we analyzed *Pkp2^+/−^* mice, which more closely reflect the human condition (Figure 1E). Consistent with findings from other monoallelic desmosomal models^17–19^, *Pkp2^+/−^* mice showed structurally and functionally normal hearts at 9 months of age: left ventricular end-diastolic diameter (LVEDD), left ventricular ejection fraction (LVEF) and tricuspid annular plane systolic excursion (TAPSE) were preserved, while myocardial collagen content or immune cell infiltration were not increased (Figure 1F-H; Table S2).

### MCMV infection causes progressive cardiac dysfunction in *Pkp2^+/−^* hearts

The absence of an overt phenotype in *Pkp2^+/−^* mice provides an opportunity to examine how viral triggers interact with genetic predisposition and cardiac immune responses to potentially trigger disease. To assess this, 3-month-old *Pkp2^+/−^* and Ctr mice were infected with 5x10^5^ plaque forming units (PFU) Δm157-MCMV-eGFP (MCMV), a mutant strain that bypasses natural killer (NK) cell-mediated clearance in C57BL/6J mice and ensures productive infection^20^. Mice of both genotypes received phosphate-buffered saline (PBS) as a control (Figure 2A). Analyses covered time points of acute (5-14 days post-infection, dpi), sub-acute (1 month post-infection, mpi) and latent MCMV infection (3; 6mpi). Both infected genotypes developed splenomegaly during acute infection, indicating immune system activation (Figure 2B). Hematoxylin-eosin (HE) and immunostaining revealed myocardial immune infiltrates composed primarily of CD8^+^ T-cell plaques and scattered CD4^+^ T-cells in either genotype during the acute phase (Figure 2C; Figure S2A). Quantitative PCR for the viral *ie1* gene confirmed viral DNA persistence in heart, salivary gland and lung with a gradual decline from acute to latent stages (Figure 2D; Figure S2B+C). Serial echocardiography from 1 to 6mpi demonstrated a progressive decline in LVEF exclusively in MCMV-infected *Pkp2^+/−^*mice as of 3mpi and worsening by 6mpi, while LVEDD remained stable across all groups (Figure 2E; Table S3-S6). Collagen content increased with age in all groups in a genotype- and infection-independent manner (Figure 2F+G, Figure S2D-F). CD8^+^ T-cell numbers decreased from 1 to 6mpi in both infected genotypes, but remained higher than in non-infected hearts (Figure 2H; Figure S2G; Table S7). Flow cytometry at 6mpi identified CD8^+^ KLRG1^+^ terminally differentiated effector memory T-cells as the dominant expanded population, consistent with MCMV-induced memory inflation in the CD8^+^ T-cell compartment^21^. Strikingly, monocytes/macrophages appeared more abundant in *Pkp2^+/−^* MCMV than in Ctr MCMV hearts (Figure 2I, Figure S3). Together, these findings show that while MCMV infection induces T-cell infiltration and persistent viral DNA in both genotypes, *Pkp2^+/−^* MCMV mice selectively develop progressive systolic dysfunction accompanied by monocyte/macrophage accumulation, establishing MCMV as a potent trigger of cardiac disease onset in *Pkp2^+/−^* mice.

**Figure 2.**
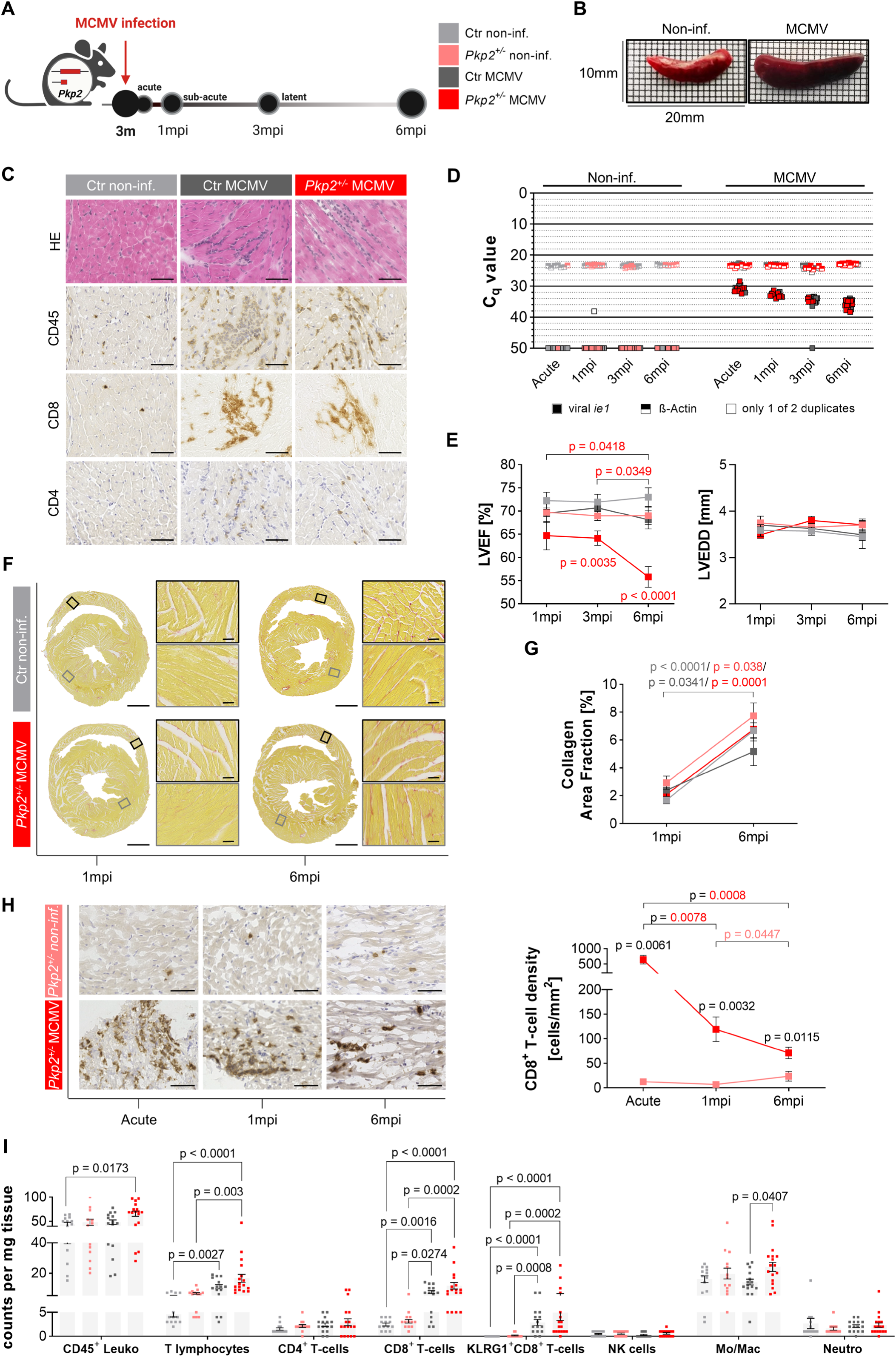
MCMV infection accelerates functional cardiac abnormalities in genetically predisposed *Pkp2^+/−^*mice. **A**, Schematic experimental design. 3-month-old *Pkp2^+/−^*and control (Ctr) mice were infected with 5x10^5^ plaque-forming units Δm157-MCMV-eGFP (MCMV) or received phosphate-buffered saline (PBS) and were analyzed at different time points post-infection. Signature colors: Ctr non-infected (light grey), *Pkp2^+/−^* non-infected (light pink), Ctr MCMV (dark grey) and *Pkp2^+/−^* MCMV (red). **B**, Splenomegaly was observed in response to acute MCMV infection. **C**, Representative images of cardiac sections stained with hematoxylin-eosin (HE) and antibodies against the surface markers CD45, CD4 and CD8 during acute infection; scale bar 50µm. **D**, Viral *ie1* (*immediate early 1*, filled squares) and murine ß-actin (half-top-filled squares) DNA copies detected by quantitative polymerase-chain reaction in heart samples from acute to latent infection represented by quantification cycle values (C_q_). Measurements were performed in technical duplicates. It is indicated by an unfilled symbol when only one value was usable; n = Acute/1mpi/3mpi/6mpi: Ctr non-infected n = 6/6/6/4, *Pkp2^+/−^* non-infected n = 1/7/8/4, Ctr MCMV n = 6/6/9/8, *Pkp2^+/−^* MCMV n = 8/9/6/13 (no statistics calculated). **E**, Quantification of echocardiographic parameters from 1 to 6 months post-infection (mpi); n = 1mpi/3mpi/6mpi: Ctr non-infected n = 6/9/7, *Pkp2^+/−^* non-infected n = 7/10/7, Ctr MCMV n = 6/14/5, *Pkp2^+/−^*MCMV n = 8/13/7. **F+G**, Representative picrosirius red staining of cardiac sections and quantification of collagen area fraction at 1 and 6mpi; scale bars: overview 1000µm, detail 50µm; n = 1mpi/6mpi: Ctr non-infected n = 4/12, *Pkp2^+/−^* non-infected n = 5/9, Ctr MCMV n = 2/10, *Pkp2^+/−^*MCMV n = 9/12. **H**, Representative images of CD8^+^ immunostaining and quantification of CD8^+^ T-cell density from acute to latent infection; scale bar 50µm; n = Acute/1mpi/3mpi/6mpi: *Pkp2^+/−^*non-infected n = 4/6/11/9, *Pkp2^+/−^* MCMV n = 7/11/6/7. **I**, Flow cytometry quantification of cardiac CD45^+^ leukocytes in absolute counts per milligram tissue at 6mpi; Ctr non-infected n = 13, *Pkp2^+/−^*non-infected n = 13, Ctr MCMV n = 15, *Pkp2^+/−^*MCMV n = 17. Data are shown as mean ± SEM. Only male mice were used (square symbol). Ordinary one-way ANOVA with or without Brown-Forsythe and Welch’s correction and Tukey’s or Dunnett’s T3 multiple comparisons, respectively or Kruskal-Wallis test with Dunn’s multiple comparisons, as appropriate for >2 groups. Unpaired t test with or without Welch’s correction or Mann-Whitney test, as appropriate for 2 groups. P-values in **E** are shown for one group over time (bracket) and across groups at one time point compared to Ctr non-infected. All other p-values can be found in Table S3-S6. m – months, LVEF – left ventricular ejection fraction, LVEDD left ventricular end-diastolic diameter, NK cells – natural killer cells, Mo/Mac – monocytes/macrophages.

### CITE-seq and transcriptomic profiling reveal genotype-dependent and virus-specific alterations of the cardiac immune compartment

To define immune cell composition in chronically infected hearts, we performed cellular indexing of transcriptomes and epitopes by sequencing (CITE-seq) of ventricular CD45^+^ leukocytes at 6mpi, excluding circulating cells by intravenous anti-CD45.2-APC injection. Two independent experiments were performed, each including 10 pooled samples (non-infected groups, n=2; MCMV groups, n=3), labeled with TotalSeq-A hashtags that were used for demultiplexing (Figure 3A; Figure S4A; Supplemental Methods). After quality control, doublet removal, and dataset integration (Figure S4B), 26,656 cells remained for further analysis. Initial clustering defined 16 immune cell groups that were annotated using canonical lineage markers for T-cells (CD3, TCRβ), B cells (CD19), NK cells (NK1.1), neutrophils (Ly6G), dendritic cells (CD11c), monocytes (Ly6C), and macrophages (CD64, CD88). Sub-clustering of the lymphoid and myeloid compartments refined the annotation to 23 distinct populations, defined by combined transcriptomic and protein signatures (Figure S5).

**Figure 3.**
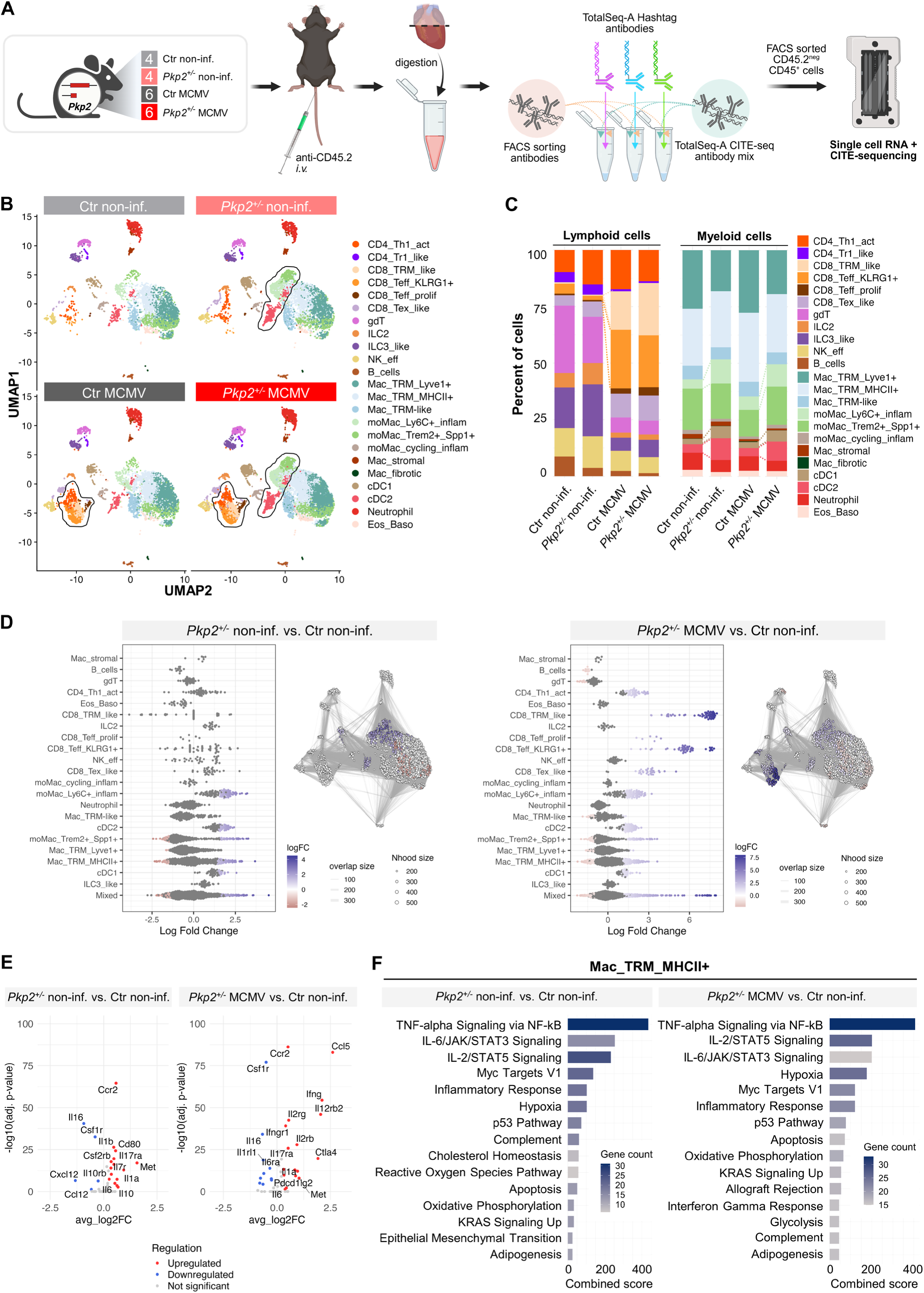
*Pkp2^+/−^*genotype is sufficient to induce subclinical accumulation of myeloid cell types before cardiac dysfunction emerges. **A**, Schematic workflow of the CITE-seq experiment. At 6 months post-infection (mpi), mice received an intravenous (*i.v.*) injection of anti-CD45.2-APC before heart dissociation (ventricles only). Samples were multiplexed with TotalSeq-A hashtags and labeled with CITE-seq antibodies plus CD45 FACS antibody. Circulating cells were excluded by sorting CD45⁺ CD45.2-APC⁻ leukocytes. Two libraries from 10 pooled samples each were sequenced; Ctr non-infected n = 4, *Pkp2^+/−^* non-infected n = 4, Ctr MCMV n = 6, *Pkp2^+/−^* MCMV n = 6. **B**, UMAP representation of Seurat clustered leukocytes (in total 26,656) split by experimental group. **C**, Relative abundance of lymphoid and myeloid immune populations across groups. **D**, Assessment of statistically relevant changes in abundance using the MiloR package in *Pkp2^+/−^* non-infected and *Pkp2^+/−^* MCMV versus Ctr non-infected, respectively. **E**, Comparative cytokine expression analysis in cardiac leukocytes in *Pkp2^+/−^* non-infected and *Pkp2^+/−^*MCMV versus Ctr non-infected, respectively. **F**, Pathway enrichment analysis using MSig_DB_Hallmark_2020 database via the enrichR pipeline on pseudobulk-dataset. Shown are the top upregulated pathways by combined score in the subcluster Mac_TRM_MHCII+ of *Pkp2^+/−^* non-infected and *Pkp2^+/−^* MCMV versus Ctr non-infected, respectively. Ctr - control, CITE-seq - cellular indexing of transcriptomes and epitopes by sequencing, FACS – fluorescence-activated cell sorting, UMAP - uniform manifold approximation and projection.

Dimensional reduction and compositional analyses revealed infection- and genotype-dependent reshaping of the cardiac immune landscape. In both genotypes, latent MCMV infection induced robust expansion of effector T-cell subsets, including activated CD4⁺ Th1 cells (CD4_Th1_act; *Cd4, Cd28, Ctla4*), tissue-resident memory- like CD8⁺ effectors (CD8_TRM_like; *Cd8a, Itga1, Gzmk, Klrc1*), terminal KLRG1⁺ effector memory (CD8_Teff_KLRG1+; *CD8a, Klrg1, Gzmb, Ccl5*), and proliferating CD8⁺ T cells (CD8_Teff_prolif; *Mki67, Top2a, Birc5*). Beyond this shared antiviral response, *Pkp2^+/−^*MCMV hearts displayed a distinct myeloid signature with a marked increase in cDC1 (*Xcr1, Clec9a, Itgae*) and antigen-presenting cDC2 cells (*Cd209a, Bhlhe40, Dpp4*), as well as in two subsets of monocyte-derived macrophages. These comprised a pro-inflammatory subset (moMac_Ly6C⁺_inflam; *Plac8, Chil3, Ly6c2*) and a reparative *Trem2^+^Spp1^+^* subset (moMac_Trem2⁺_Spp1⁺; *Trem2, Spp1, Fabp5*), the latter previously associated with profibrotic and tissue-remodeling functions in the infarcted heart^22^. Interestingly, these subsets were already elevated in non-infected *Pkp2^+/−^* myocardium compared to Ctr non-infected hearts, indicating genotype-dependent priming of the myeloid compartment in the *Pkp2* heterozygous setting (Figure 3B+C). No additional populations were selectively enriched in the *Pkp2^+/−^*MCMV group, indicating convergence of viral and genetic factors on a common immune response. To validate these compositional shifts, we performed differential abundance testing using the MiloR framework, comparing *Pkp2^+/−^* non-infected and *Pkp2^+/−^* MCMV hearts to Ctr non-infected mice, thereby allowing assessment of genotype-dependent baseline changes and infection-driven responses on the *Pkp2^+/−^*background. moMac_Ly6C+_inflam, cDC1, and cDC2 populations were significantly enriched when comparing non-infected *Pkp2^+/−^*and Ctr hearts, confirming genotype-dependent expansion of myeloid subsets. Following MCMV infection (*Pkp2^+/−^* MCMV vs. Ctr non-infected), CD8_TRM_like and CD8_Teff_KLRG1+ populations overall displayed the strongest expansion, while the underlying *Pkp2*-dependent myeloid alterations persisted (Figure 3D; Figure S6A).

To identify inflammatory mediators underlying these compositional changes, we performed differential expression analysis of cytokines and their receptors across all immune cell types (Figure 3E). Even in the absence of infection, non-infected *Pkp2^+/−^*hearts showed a distinct pro-inflammatory profile: *Ccr2* was strongly upregulated, indicating enhanced recruitment and activation of monocytes and macrophages via ligands like *Ccl2, Ccl7 or Ccl12*^23^. Elevated expression of *Il1b*, *Il1a,* and *Il6* further supported activation of innate inflammatory pathways, while increased *Il17r* and *Il17ra* indicated responsiveness to IL-17 signaling, typically associated with pro-inflammatory Th17 cells^24^. In both genotypes, MCMV induced upregulation of *Ccl5*, *Ifng* and its receptor *Ifngr1*, as well as *Il2rg* and *Il2rb*, components of the interleukin-2 receptor, implicated in regulating infection-associated T-cell responses^25,26^. In addition, inhibitory molecules such as *Ctla4* were upregulated (Figure 3E). When comparing *Pkp2^+/−^*MCMV to Ctr MCMV to isolate the genotype effect, a persistent myeloid-dominant inflammatory signature emerged – again driven by *Ccr2* and related mediators already identified in the *Pkp2^+/−^* non-infected hearts (Figure S6B).

Finally, pseudobulk differential expression was performed per cell type and subjected to pathway enrichment analysis using the database MSigDB Hallmarks. In Mac_TRM_MHCII+ macrophages, both comparisons (*Pkp2^+/−^* non-infected vs. Ctr non-infected; *Pkp2^+/−^* MCMV vs. Ctr non-infected) revealed consistent enrichment of immune activation pathways, with TNF-α signaling via NF-κB as the top hit. Additional pathways included IL-6/JAK/STAT3 signaling, IL-2/STAT5 signaling and a broad inflammatory response signature (Figure 3F, Figure S6C). Consistently, STAT3 protein levels were increased in infected and non-infected *Pkp2^+/−^* mice (Figure S6D). Together, single-cell RNA and CITE-sequencing captured a subclinical accumulation and activation of specific monocyte/macrophage populations already in phenotypically unaffected non-infected *Pkp2^+/−^* mice, similar to those observed in *Pkp2^+/−^* MCMV mice with systolic dysfunction. Therefore, the mutation itself already reshapes the cardiac immune landscape. Nevertheless, an additional trigger is required to actually cross the threshold for developing a clinical phenotype.

### The important role of the CCL2-CCR2 axis

Next, we sought to identify soluble mediators that might attract or maintain inflammatory cells in whole-heart protein samples and serum. This was done using proximity extension assay-based proteomics at 6mpi (corresponding to 9 months of age). Given the strong *Ccr2* upregulation observed in cardiac leukocytes, we expected to identify one of its attractants *Ccl2, Ccl7* or *Ccl12*^23^. Interestingly, aside from CCL2, several additional mediators were significantly upregulated in phenotypically unaffected *Pkp2^+/−^* hearts (Figure 1E-H), including chemokines involved in monocyte and macrophage recruitment (CCL2, CCL4, CXCL1), pro-inflammatory cytokines (IL-1β) and immunoregulatory mediators (PD-L2, CCL22). Additionally, the metabolic regulator FGF21 was increased and has been reported to exert cardioprotective effects^27,28^. In contrast, these factors were not elevated in serum. This cytokine response was amplified upon MCMV infection in *Pkp2^+/−^* mice: in addition to FGF21, CCL2, CXCL1, CCL22, and CCL4, we observed upregulation of CCL12 and CCL5. In serum, the most prominent change was a slight induction of IL-6, consistent with systemic MCMV-driven inflammation, whereas heart-specific mediators remained largely unchanged, indicating a predominantly local cardiac response (Figure 4A; Figure S7A-C; Table S8+S9). Thus, these findings demonstrate that *Pkp2*-deficiency alone is sufficient to elicit an early subclinical pro-inflammatory milieu in the heart, consistent with recruitment of pro-inflammatory Ly6C^+^ CCR2^+^ monocytes. Even though MCMV induced a CD8⁺ T-cell response, this was associated with only a modest change in the overall cytokine spectrum compared with the genotype-driven signals.

**Figure 4.**
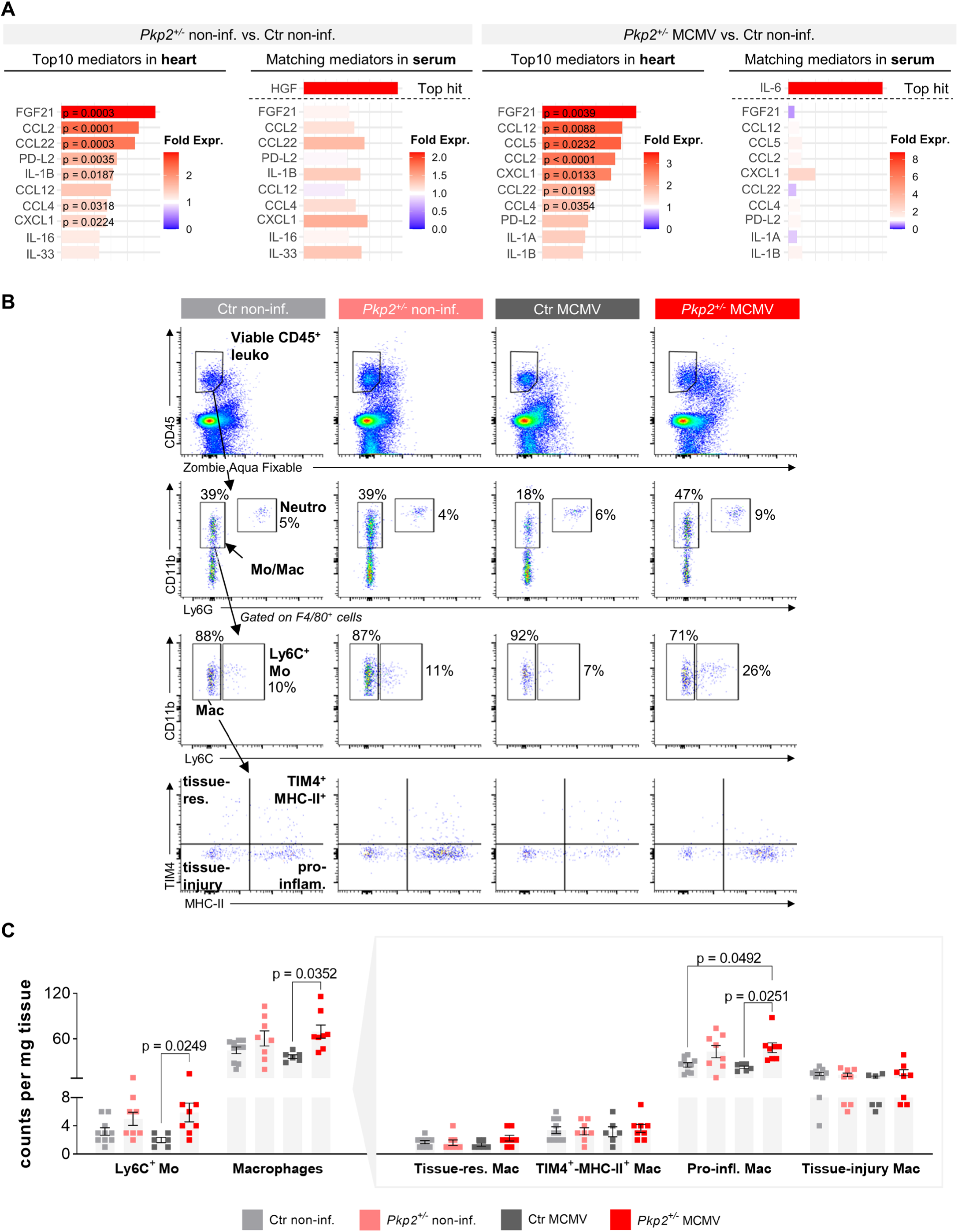
Heart-specific subclinical upregulation of cytokines in non-infected *Pkp2^+/−^* mice. **A**, Cytokine profiling based on proximity extension assay at 6 months post-infection (mpi). Shown are the top 10 differentially expressed heart mediators with corresponding serum levels, plus the most altered serum-specific mediator. Expression ratios compare *Pkp2^+/−^* non-infected and *Pkp2^+/−^*MCMV to Ctr non-infected, respectively; n = heart/serum: Ctr non-infected n = 10/10, *Pkp2^+/−^* non-infected n = 10/10, Ctr MCMV n = 10/7, *Pkp2^+/−^*MCMV n = 10/8. **B+C**, Representative gating and quantification of a myeloid-tailored flow cytometry panel of cardiac CD45^+^ leukocytes from 6mpi hearts shown as absolute counts per milligram tissue; Ctr non-infected n = 10, *Pkp2^+/−^* non-infected n = 8, Ctr MCMV n = 6, *Pkp2^+/−^* MCMV n = 8. Data are shown as mean ± SEM. Only male mice were used (square symbol). Unpaired t test with or without Welch’s correction or Mann-Whitney test, as appropriate for 2 groups. Ordinary one-way ANOVA with or without Brown-Forsythe and Welch’s correction and Tukey’s or Dunnett’s T3 multiple comparisons, respectively or Kruskal-Wallis test with Dunn’s multiple comparisons, as appropriate for >2 groups. Ctr – control, Neutro – neutrophils, Mo/Mac – monocytes/macrophages, Mo – monocytes, Mac – macrophages.

We next examined whether the expansion of myeloid populations in *Pkp2^+/−^*hearts could be recapitulated using a macrophage-tailored flow cytometry panel. While in *Pkp2^+/−^* MCMV hearts, Ly6C^+^ monocytes (typically also CCR2^+^) and pro-inflammatory macrophages (TIM4^−^, MHC-II^+^) were significantly increased, non-infected *Pkp2^+/−^*hearts, only showed a trend towards an increased abundance of these subsets (Figure 4B+C), indicating that single-cell analysis is more sensitive at detecting an early and cell-specific immune response. Together, these data show that *Pkp2^+/−^* hearts mount a local cytokine response amplified by MCMV, with recruitment of Ly6C^+^ CCR2^+^ monocytes highlighting the CCL2/CCR2 axis as a central driver of myocardial inflammation.

### MCMV infection in *Ttn^+/−^* mice does not induce cardiac dysfunction

To test whether the combined effects of genetic predisposition and viral infection reflect a broader principle in genetic cardiomyopathies, as observed in human myocarditis patients^7^, we applied the same experimental setup in mice with a heterozygous global titin truncating mutation (*Ttn^+/−^*), a model developing cardiomyopathy under experimental stress conditions^11,29^. 9-month-old *Ttn^+/−^* mice were indistinguishable from controls (Ctr), regarding cardiac function, collagen deposition and immune cell abundance (Figure S8A–D). In accordance with the *Pkp2* work, we infected 3-month-old *Ttn^+/−^* and Ctr mice with 5×10^5^ PFU MCMV and performed analyses at 1, 3 and 6mpi (Figure 5A). Non-infected *Ttn^+/−^*and Ctr mice received PBS and were combined into one control group. Echocardiography and histology showed no genotype- or infection-dependent functional or structural alterations at any time point following infection (Figure 5B+C; Figure S8E+F; Table S10-S13). At 6mpi, immunostaining revealed scattered CD8^+^ T-cells in infected hearts irrespective of genotype (Figure 5D). Flow cytometry confirmed infection-induced expansion of cytotoxic CD8^+^ T-cells and KLRG1^+^ CD8^+^ T-cells in both genotypes, with no change in monocyte/macrophage numbers (Figure 5E, Figure S8G).

**Figure 5.**
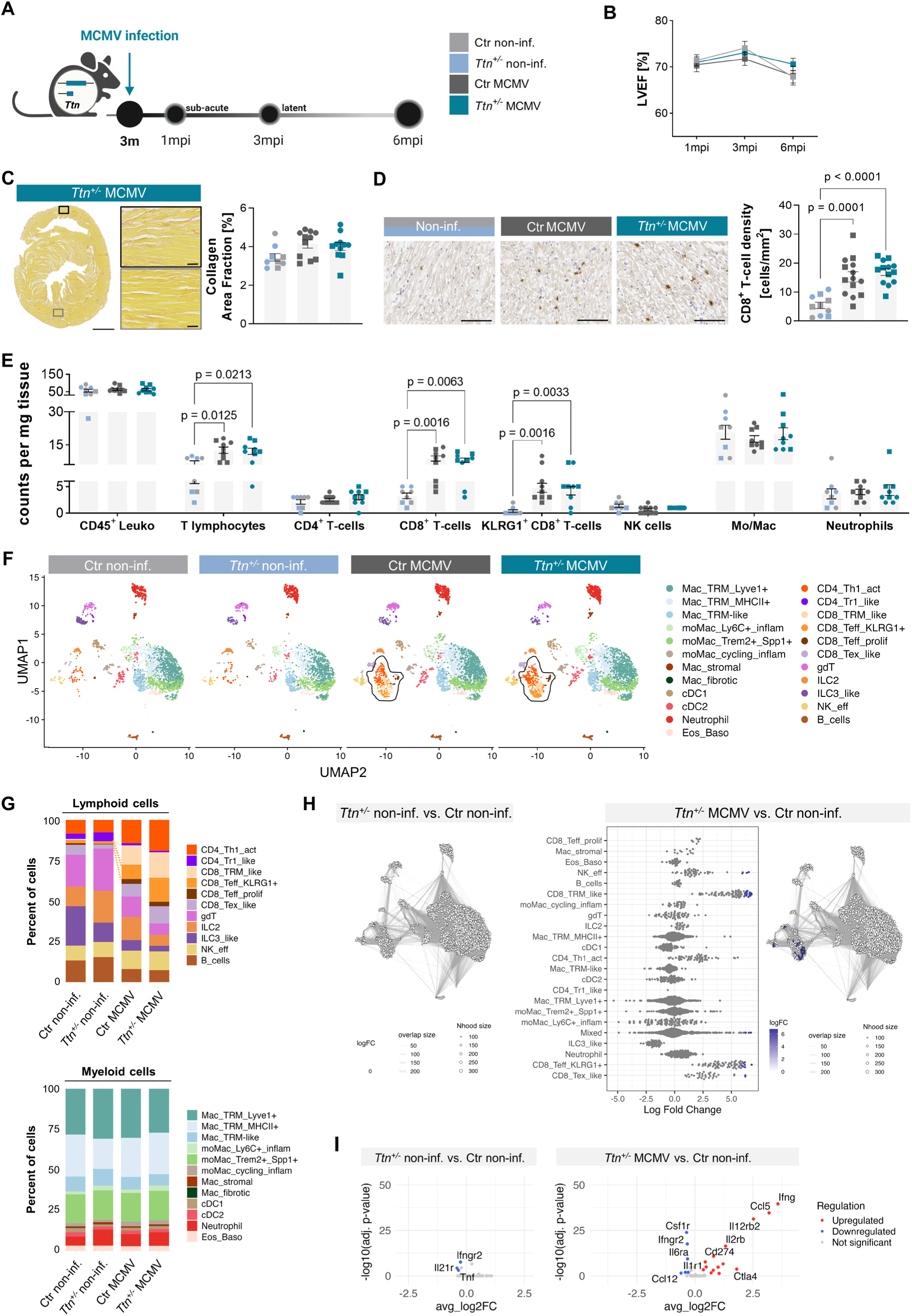
MCMV does not trigger disease onset in a genetic mouse model of dilated cardiomyopathy with heterozygous loss of the sarcomeric gene *Ttn* (*Ttn^+/−^*). **A,** Schematic experimental design. 3-month-old *Ttn^+/−^* and control (Ctr) mice were infected with 5x10^5^ plaque-forming units Δm157-MCMV-eGFP (MCMV) or received phosphate-buffered saline (PBS) and were analyzed at different time points post-infection. Signature colors: Ctr non-infected (light grey), *Ttn^+/−^* non-infected (light blue), Ctr MCMV (dark grey) and *Ttn^+/−^* MCMV (blue). **B**, Left ventricular ejection fraction (LVEF) measured by echocardiography from 1 to 6 months post-infection (mpi); n = 1mpi/3mpi/6mpi: Ctr non-infected n = 5/5/4, *Ttn^+/−^* non-infected n = 5/5/5, Ctr MCMV n = 9/9/9, *Ttn^+/−^* MCMV n = 9/9/9. **C**, Representative picrosirius red stained *Ttn^+/−^* MCMV cardiac section and quantification of collagen area fraction at 6mpi; scale bars: overview 1000µm, detail 50 µm; Ctr non-infected + *Ttn^+/−^* non-infected n = 9, Ctr MCMV n = 11, *Ttn^+/−^* MCMV n = 11. **D**, Representative images of CD8^+^ immunostaining in 6mpi heart sections and corresponding quantification of CD8^+^ T-cell density; scale bar 50µm; Ctr non-infected + *Ttn^+/−^* non-infected n = 10, Ctr MCMV n = 14, *Ttn^+/−^*MCMV n = 13. **E**, Quantification of a lymphoid-tailored flow cytometry panel of cardiac CD45^+^ leukocytes from 6mpi hearts shown as absolute counts per milligram tissue; Ctr non-infected + *Ttn^+/−^* non-infected n = 8, Ctr MCMV n = 9, *Ttn^+/−^* MCMV n = 9. **F**, UMAP representation of 12,660 cardiac leukocytes identified by CITE-seq split by experimental group at 6mpi; Ctr non-infected n = 2, *Ttn^+/−^* non-infected n = 2, Ctr MCMV n = 3, *Ttn^+/−^* MCMV n = 3. **G**, Relative cell type abundances across groups in lymphoid and myeloid cells at 6mpi. **H**, Assessment of statistically relevant changes in abundance using the MiloR package (p-value 0.1) in *Ttn^+/−^*non-infected and *Ttn^+/−^*MCMV versus Ctr non-infected, respectively. **I**, Comparative cytokine expression analysis in cardiac leukocytes (p-value 0.1) in *Ttn^+/−^* non-infected and *Ttn^+/−^*MCMV versus Ctr non-infected, respectively. Data are shown as mean ± SEM. Male: squares; female: circles. Ordinary one-way ANOVA with or without Brown-Forsythe and Welch’s correction and Tukey’s or Dunnett’s T3 multiple comparisons, respectively or Kruskal-Wallis test with Dunn’s multiple comparisons, as appropriate. Ctr – control, UMAP – uniform manifold approximation and projection, CITE-seq – cellular indexing of transcriptomes and epitopes by sequencing, NK cells – natural killer cells.

To capture subtle immune shifts, CITE-seq was performed on CD45⁺ cells at 6mpi (10 hashed samples: Ctr non-inf n=2, *Ttn^+/−^* non-inf n=2, Ctr MCMV n=3, *Ttn^+/−^* MCMV n=3; Figure S9A-C). Relative cell type abundance did not differ between non-infected *Ttn^+/−^*and Ctr hearts, while both infected groups displayed expansion of activated T-cell subsets (CD4_Th1_act, CD8_TRM_like, CD8_Teff_KLRG1+, CD8_Teff_prolif, CD8_Tex_like; Figure 5F+G). Myeloid proportions remained unchanged across all groups, in contrast to the genotype-driven myeloid skewing observed in *Pkp2^+/−^*mice. MiloR^30^ testing confirmed no differences between non-infected genotypes, justifying their pooling in subsequent analyses. MCMV infection (*Ttn^+/−^* MCMV vs. Ctr non-infected) enriched CD8_TRM_like, NK_eff, CD8_Teff_KLRG1+ and CD8_Tex_like clusters, mirroring findings in Ctr MCMV hearts, while there was no difference between both infected genotypes (Figure 5H; Figure S9D+E). Differential cytokine expression analysis supported these results: *Ttn^+/−^* non-infected hearts showed no upregulated inflammatory mediators, while MCMV infection induced *Ifng, Ccl5* and IL-2 and IL-12 receptor components (*Il2rb, Il12rb2*) in both genotypes compared to non-infected animals (Figure 5I, Figure S9F). Thus, in contrast to *Pkp2^+/−^* mice, *Ttn^+/−^*hearts exhibited no genotype-driven immune remodeling. All observed changes were attributable to MCMV infection, underscoring the specificity of immune activation in the *Pkp2^+/−^*model.

### Immune cell remodeling and phenotype development in aged *Pkp2^+/−^* mice

To determine how inflammation, immune responses and cardiac function evolve with age in the absence of viral infection, we longitudinally analyzed *Pkp2^+/−^* and Ctr mice from 3 to 15 months of age (Figure 6A). Aging progressively reshapes the cardiac immune landscape^31,32^. We thus hypothesized that interaction with *Pkp2* deficiency might eventually unmask a cardiac phenotype.

**Figure 6.**
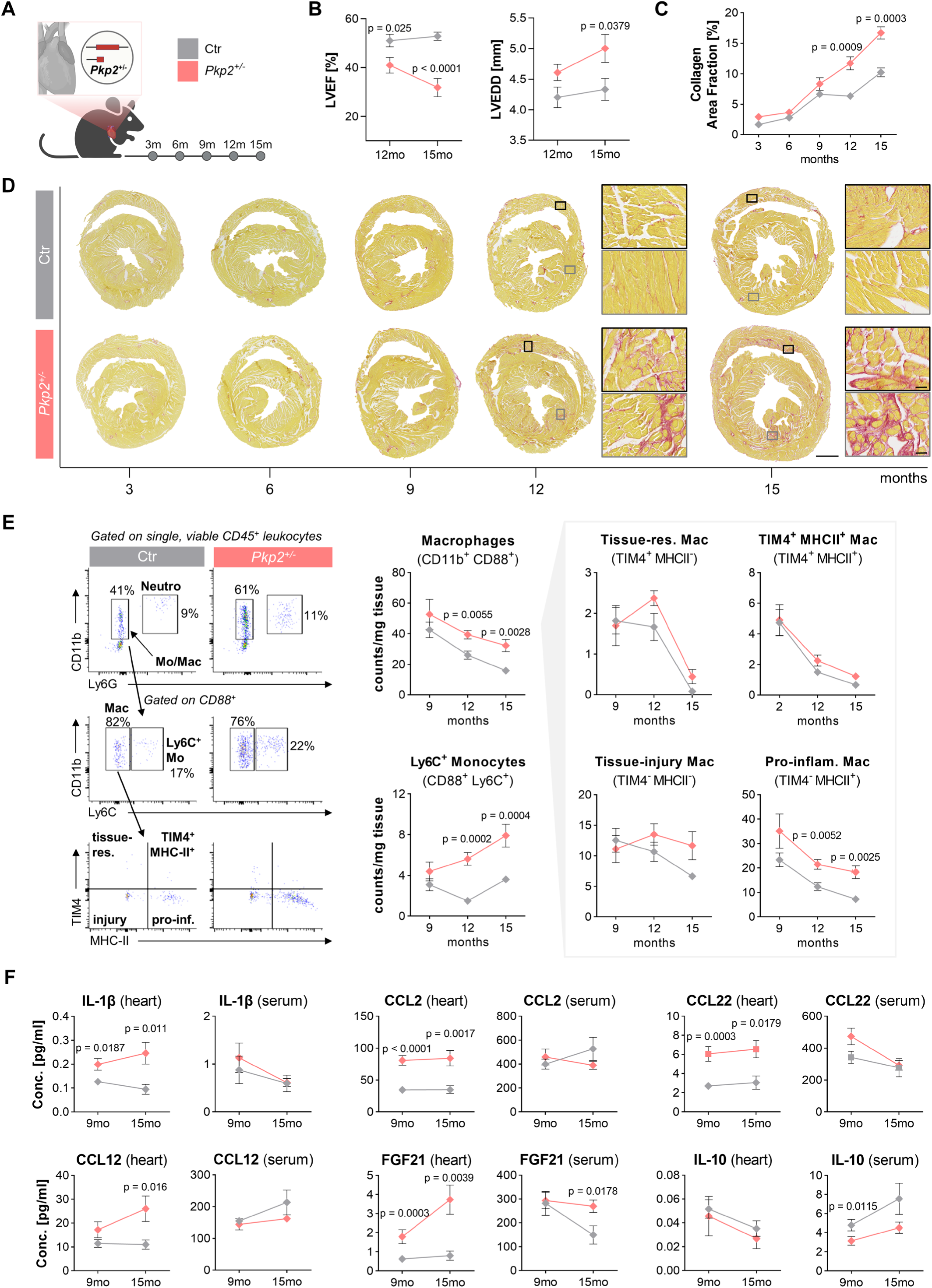
Progressive development of a cardiac phenotype in aged *Pkp2^+/−^* mice. **A**, Schematic overview of the experimental time points in *Pkp2^+/−^*and Ctr mice. **B**, Quantification of echocardiographic parameters at 12 and 15 months (mo); n = 12mo/15mo: Ctr n = 9/8, *Pkp2^+/−^* n = 9/9. **C+D**, Quantification of collagen area fraction and representative picrosirius red staining of cardiac sections from 3 to 15mo; scale bars: overview 1000µm, detail 50µm; n = 3mo/6mo/9mo/12mo/15mo: Ctr n = 4/7/12/6/7, *Pkp2^+/−^* n = 5/6/10/8/10. **E**, Representative gating and quantification of different cardiac CD45^+^ leukocytes at 9, 12 and 15mo by flow cytometry in counts per milligram tissue; n = 9mo/12mo/15mo: Ctr n = 11/6/12, *Pkp2^+/−^* n = 10/8/10. **F**, Cytokine profiling based on proximity extension assay at 9 and 15mo; n = 9mo/15mo: Ctr n = 10/6, *Pkp2^+/−^* n = 10/10. Data are shown as mean ± SEM. Unpaired t test with or without Welch’s correction or Mann-Whitney test, as appropriate for 2 groups. Ordinary one-way ANOVA with or without Brown-Forsythe and Welch’s correction and Tukey’s or Dunnett’s T3 multiple comparisons, respectively or Kruskal-Wallis test with Dunn’s multiple comparisons, as appropriate for >2 groups. P-values for comparisons between groups at a given time point are shown in the graph; p-values for longitudinal comparisons within groups are provided in Table S14-S17. Ctr – control, LVEF – left ventricular ejection fraction, LVEDD – left ventricular end-diastolic diameter, Neutro – neutrophils, Mo/Mac – monocytes/macrophages, Mo – monocytes, Mac – macrophages.

Up to 9 months, *Pkp2^+/−^* mice maintained normal cardiac function despite a myeloid-skewed inflammatory state (Figures 2E+3D+4A; *Pkp2^+/−^* non-infected at 6mpi). From 12 months onward, however, LVEF declined and LVEDD increased, progressing further by 15 months, while Ctr hearts remained stable, indicating worsening systolic function and ventricular remodeling (Figure 6B; Table S14+S15). Collagen content increased with age in both genotypes but rose more steeply in *Pkp2^+/−^* hearts, with fibrosis localized to the LV and RV endocardial layers (Figure 6C+D; Figure S10A). Flow cytometry revealed progressive accumulation of inflammatory Ly6C^+^ monocytes and persistently elevated total macrophages in *Pkp2^+/−^* hearts from 12 months onward. Among macrophage subsets, pro-inflammatory antigen-presenting (TIM4^−^ MHC-II^+^) cells were consistently increased, despite an overall age-related decline in this population (Figure 6E). Cytokine profiling supported these findings: IL-1β, CCL2 and CCL22 were elevated solely in *Pkp2^+/−^*hearts as of 9 months, while CCL12 increased only at 15 months, whereas none of these mediators were altered in serum, indicating a localized rather than a systemic inflammatory response. Cardioprotective FGF21 was already elevated in *Pkp2^+/−^*hearts as of 9 months and additionally increased in serum at 15 months, suggesting an emerging compensatory systemic response. In contrast, IL-10 remained unchanged in cardiac tissue but was reduced in serum, consistent with a shift towards a pro-inflammatory milieu in *Pkp2^+/−^* mice (Figure 6F; Table S16+17). Together, these data show that physiological aging unmasks systolic dysfunction, LV dilation, fibrosis and a sustained cardiac inflammatory state in *Pkp2^+/−^*mice. Between 9 and 12 months, inflammatory markers rise before the onset of overt fibrosis and contractile decline, defining a therapeutic window during which anti-inflammatory intervention may attenuate immune activation and fibrotic remodeling to preserve cardiac function.

To identify signaling networks associated with this early inflammatory state, kinome profiling was performed in 12-month-old *Pkp2^+/−^*versus Ctr hearts. The significantly regulated kinases were uniformly upregulated, indicating a coordinated activation of inflammatory, immune, and compensatory signaling pathways (Figure 7A; Figure S10B). IKBKE, PRKD1 and CSNK2A1 reflect activation and regulation of the NF-κB pathway^33,34^ sustaining chronic low-grade inflammation. FES, FGR, LCK and EPHA1 indicate enhanced immune-cell signaling and cell-cell communication, supporting the immune activation observed at this stage. In parallel, MET, AKT3, PRKG1/2, CSNK2A1 and RPS6KB2 suggest engagement of pro-survival and metabolic stress-response pathways via PI3K-AKT-mTOR signaling^35^. STRING network analysis showed connections between these immune-related and compensatory kinases (Figure 7B), while KEGG pathway enrichment also reflected the dual activation of different pro-inflammatory signaling pathways alongside pro-survival cascades, indicating cardiac stress adaptation (Figure 7C). To assess whether kinase activation translated into downstream pathway engagement at later stages, we analyzed hearts from 15-month-old mice by Western blot (Figure 7D). Consistent with persistent NF-κB activation, both total and phosphorylated p65 (p_i_-p65) were increased in *Pkp2^+/−^* hearts compared with Ctr. These molecular changes coincide with the stage of functional decline and fibrosis in aged *Pkp2^+/−^* hearts, confirming sustained NF-κB signaling and chronic inflammatory stress.

**Figure 7.**
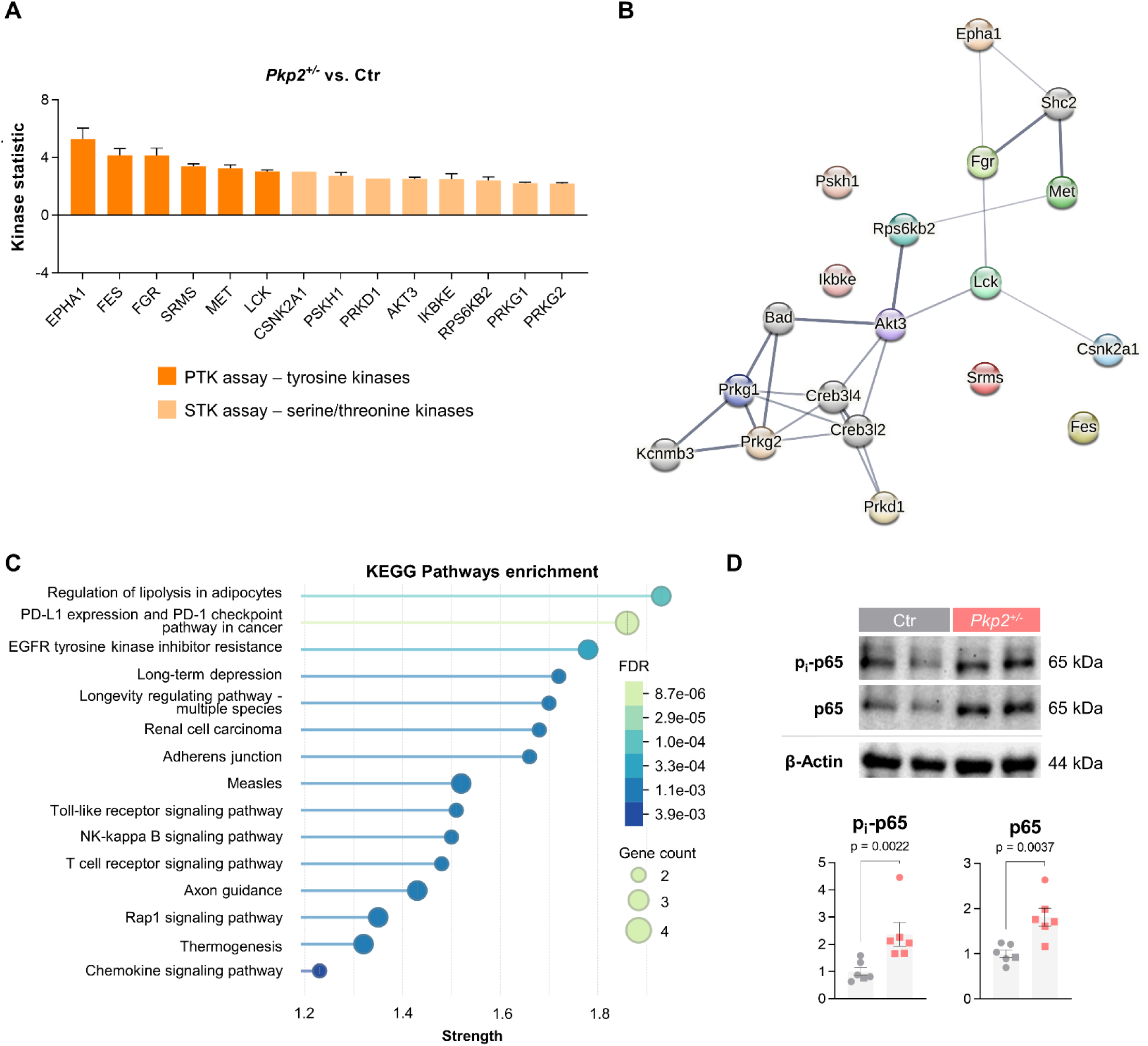
Activation of inflammatory and compensatory signaling in *Pkp2^+/−^* hearts at 12 months. **A**, Comparative PamChip kinome profiling of *Pkp2^+/−^* hearts versus Ctr hearts at 12 months; Ctr n = 4, *Pkp2^+/−^* n = 3. Bar plot representation of altered tyrosine (PTK) and serine/threonine (STK) kinases ranked by mean kinase statistic; Ctr n = 4, *Pkp2^+/−^* n = 3. **B**, STRING interactions at medium confidence level. Line thickness correlates with predicted interaction strength. **C**, KEGG pathway enrichment of identified signaling networks. **D**, Western blot quantification and representative blot snips of phosphorylated p65 (p_i_-p65, Ser536) and p65 protein levels in 15-month-old hearts normalized to β-Actin levels; Ctr n = 6, *Pkp2^+/−^* n = 6. Data are shown as mean ± SEM. Male: squares; female: circles. Unpaired t test with or without Welch’s correction or Mann-Whitney test, as appropriate. Ctr – control, KEGG – Kyoto Encyclopedia of Genes and Genomes.

### Early inflammatory signatures in homozygous *Pkp2^−/−^* hearts point towards infiltrating monocytes as a therapeutic target

The overall phenotype of heterozygous *Pkp2^+/−^* mice was moderate and became apparent only upon the inflammatory trigger and at advanced age. To model a more pronounced phenotype, we analyzed homozygous *Pkp2^−/−^*mice. Because 6-8-week-old *Pkp2^−/−^* hearts already displayed extensive fibrosis (Figure 1C), mice were examined at 3-4 weeks to capture earlier inflammatory stages (Figure 8A). However, even at this age, echocardiography revealed functional impairment, albeit less pronounced than at 6-8 weeks (Figure 8B). Interestingly, HE-stained hearts showed clear immune cell accumulation between cardiomyocytes in both ventricles, which stained positive for the macrophage marker CD68 (Figure 8C+D). Flow cytometry confirmed significant infiltration of F4/80^+^ macrophages and Ly6C^+^ monocytes in *Pkp2^−^_/-_* hearts. Sub-characterization revealed expansion of recruited pro-inflammatory (TIM4^−^MHC-II^+^) and tissue-injury-associated macrophages (TIM4^−^ MHC-II^−^), alongside reduced tissue-resident macrophages (TIM4^+^ MHC-II^−^), indicating a monocyte-derived inflammatory response and identifying these cells as a potential therapeutic target (Figure 8E). Cytokine analysis showed broad induction of inflammatory mediators in *Pkp2^−/−^*hearts, including monocyte/macrophage chemoattractants (CCL2, CCL12, CCL4, CCL22, CCL5), neutrophil chemokines (CXCL1, CXCL2), eosinophil-recruiting CCL11 and pro-inflammatory cytokines (IL-1α, IL-1β). Upregulation of immune checkpoints PD-L2 and CTLA4 suggested active immunoregulation within the myocardium. These changes were largely heart-restricted, with serum mediators mainly unaltered (Figure 8F). Recruitment of CCR2^+^ monocytes via CCL2 and CCL12 thus represents a central feature of the *Pkp2^−/−^* model.

**Figure 8.**
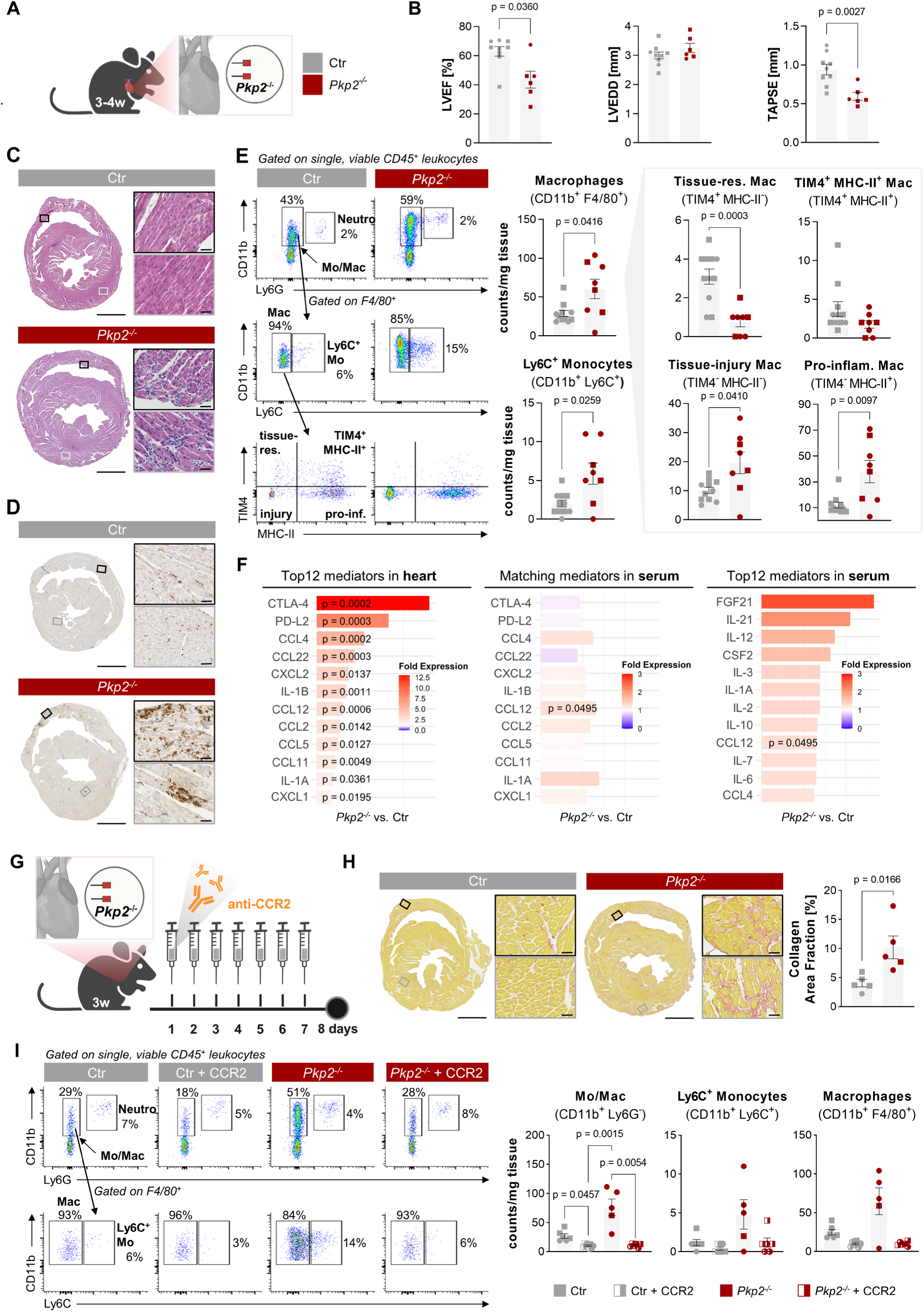
Anti-CCR2 intervention reduced numbers of recruited inflammatory monocytes and macrophages in more severely affected homozygous *Pkp2^−/−^*mice. **A,** Schematic representation of homozygous cardiac-specific *Pkp2* knockout (*Pkp2^−/−^*) mice at 3-4 weeks of age; signature colors: Ctr - grey, *Pkp2^−/−^* - dark red. **B**, Quantification of echocardiographic parameters at 3-4 weeks; Ctr n = 9, *Pkp2^−/−^* n = 6. **C**, Representative images of cardiac sections stained with hematoxylin-eosin (HE); scale bars: overview 1000µm, detail 50µm. **D**, Representative images of cardiac immunostaining against the macrophage surface marker CD68; scale bars: overview 1000µm, detail 50µm. **E**, Representative gating and quantification of different cardiac CD45^+^ leukocytes at 3-4 weeks by flow cytometry, shown in counts per milligram tissue; Ctr n = 11, *Pkp2^−/−^*n = 8. **F**, Cytokine profiling based on proximity extension assay at 3-4 weeks comparing expression ratios of *Pkp2^−/−^* versus Ctr heart and serum samples. Shown are the top 12 differentially expressed heart mediators with corresponding serum levels along with the top 12 most altered serum-specific mediators; n = heart/serum: Ctr n = 10/10, *Pkp2^−/−^* n = 10/10. **G**, Schematic representation of anti-CCR2 antibody treatment of *Pkp2^−/−^* and Ctr mice at 3 weeks of age. **H**, Representative picrosirius red staining of cardiac sections and quantification of collagen area fraction without anti-CCR2 treatment; scale bars: overview 1000µm, detail 50µm; Ctr n = 5, *Pkp2^−/−^* n = 5. **I**, Representative flow cytometry gating and quantification of different cardiac CD45^+^ leukocytes following anti-CCR2 treatment represented in counts per milligram tissue; untreated – filled symbol, anti-CCR2 treated – half-filled symbol; Ctr n = 6, Ctr+CCR2 n = 8, *Pkp2^−/−^* n = 5, *Pkp2^−/−^*+CCR2 n = 6. Data are shown as mean ± SEM. Male: squares; female: circles. Unpaired t test with or without Welch’s correction or Mann-Whitney test, as appropriate for 2 groups. Ordinary one-way ANOVA with or without Brown-Forsythe and Welch’s correction and Tukey’s or Dunnett’s T3 multiple comparisons, respectively or Kruskal-Wallis test with Dunn’s multiple comparisons, as appropriate for >2 groups. w – weeks, Ctr – control, LVEF – left ventricular ejection fraction, LVEDD – left ventricular end-diastolic diameter, TAPSE – tricuspid annular plane systolic excursion, Neutro – neutrophils, Mo/Mac – monocytes/macrophages, Mo – monocytes, Mac – macrophages.

To examine whether this monocyte-driven inflammatory state can be modulated, we treated 3-week-old *Pkp2^−/−^* and Ctr mice with an anti-CCR2 antibody (Figure 8G). Treatment duration was limited to 7 days to prevent the formation of anti-drug antibodies, which could compromise target engagement^36^. At the beginning of treatment, mild fibrotic remodeling was already evident in *Pkp2^−/−^* hearts (Figure 8H). Despite this, CCR2 blockade markedly reduced total cardiac monocytes/macrophages, with similar trends toward decreases in Ly6C^+^ monocytes, F4/80^+^ macrophages (Figure 8I), tissue-injury-associated and pro-inflammatory subsets (Figure S11A+B), confirming the monocytic origin of the likely disease-driving myeloid populations in this model. Fibrosis and cardiac function remained unchanged due to the short treatment period (data not shown). These data confirm effective depletion of CCR2^+^ myeloid populations *in vivo* and indicate that the cellular and tissue state at treatment onset likely shapes the measurable downstream effects.

Across genetic, viral and aging models, our data show that, in particular, the *Pkp2^+/−^*genotype and therefore disturbed desmosomal proteins predispose to immune remodeling that accelerates with secondary stressors, whereas complete loss of *Pkp2* rapidly triggers monocyte-driven inflammation and cardiac dysfunction, highlighting inflammatory monocytes as a therapeutic target population.

## 6. Discussion

In this study, we demonstrate that murine cytomegalovirus (MCMV) infection acts as a potent, genotype-dependent environmental modifier, unmasking early disease in phenotypically unaffected cardiac-restricted *Pkp2^+/−^* mice, but not in *Ttn^+/−^*mice. While both models exhibited persistent effector-memory CD8⁺ T-cells in the myocardium during latent infection, only *Pkp2^+/−^* MCMV mice developed progressive systolic dysfunction accompanied by increased recruitment of monocytes and macrophages - findings that were absent in *Ttn^+/−^* MCMV mice. These results reveal that MCMV does not trigger a generic response in genetic cardiomyopathies, but instead amplifies desmosome-related inflammation, driving ACM-like pathology. Even in the absence of infection, the *Pkp2^+/−^* genotype alone induced a subclinical pro-inflammatory state, which is mediated by the CCL2/CCR2 signaling pathway. With aging, *Pkp2^+/−^* mice developed fibrosis and overt contractile dysfunction in the absence of MCMV, marking a transition from compensated inflammation to pathological remodeling. Together, these findings establish viral infection as a disease-modifying factor in ACM, delineate mutation-specific inflammatory mechanisms distinguishing ACM from other cardiomyopathies, and identify the subclinical inflammatory phase as a promising therapeutic window for anti-inflammatory intervention.

As a cardiotropic virus with high seroprevalence in the human population, CMV establishes lifelong latency and can reactivate, for instance, under inflammatory stress^37,38^. MCMV closely mirrors the human infection, reproducing key virological and immunological features, including reshaping of the T-cell compartment through memory inflation, which maintains chronic low-grade inflammation^21^. To prevent rapid NK cell-mediated clearance typical for wild-type MCMV in C57BL/6 mice, we used a m157-deficient strain (Δm157-MCMV-eGFP) that bypasses recognition by the activating NK cell receptor Ly49H and shifts the antiviral defense toward a sustained CD8^+^ T-cell response^20,39^. Consistently, we observed prominent CD8^+^ T-cell plaques in acutely infected hearts that declined but persisted up to 6mpi. A subset of these cells expressed KLRG1, indicating an effector-memory phenotype characteristic of CMV-driven memory inflation and a chronic pro-inflammatory milieu^40^. Importantly, Ctr MCMV mice in both genetic models did not develop systolic dysfunction or fibrosis, reflecting asymptomatic latent infection in immunocompetent hosts. In contrast, *Pkp2^+/−^*mice with latent MCMV infection developed progressive systolic dysfunction without structural remodeling, establishing MCMV as a potent second hit that unmasks disease in a genetically primed background. Infection amplified the pre-existing inflammatory state in *Pkp2^+/−^* mice, marked by early cytokine activation and myeloid cell recruitment. At the same time, additional CD8^+^ T-cell infiltration further enhanced cytokine signaling and was associated with functional decline. Previous studies have shown that chronic inflammatory mediators can exert negative inotropic effects^41^ and destabilize desmosomal integrity^42^, thereby impairing cardiac contractility, providing a possible explanation for the observed phenotype in our model. However, this MCMV-induced phenotype was genotype-specific. *Ttn^+/−^* mice carrying a truncating titin variant^11^, despite mounting a similar CD8^+^ T-cell response, showed preserved cardiac function and no signs of additional inflammation throughout the entire time course. Previous studies demonstrated that stimulation with angiotensin II or isoproterenol induced systolic failure and fibrosis in phenotypically silent *Ttn^+/−^* mice, indicating that the described pathology is linked to biomechanical rather than inflammatory stress^11,29^. Thus, the mechanisms by which *Pkp2* and *Ttn* mutations lead to cardiomyopathy diverge: desmosomal disruption generates an intrinsic pro-inflammatory state that viral infection can amplify, whereas titin mutation-driven disease appears to be dominated by stress-responsive or possibly autoimmune mechanisms. Together, these findings establish CMV as a context-dependent modifier of genetic cardiomyopathy in mice, revealing that viral infection amplifies mutation-specific immune mechanisms rather than acting as a universal pathogenic trigger across genotypes and model systems.

In the absence of external triggers, *Pkp2^+/−^* mice exhibited a slowly progressive phenotype that mirrors the clinical trajectory of ACM, transitioning from a concealed phase to overt cardiac dysfunction. Cardiac function remained preserved up to 9 months of age, followed by the onset of interstitial fibrosis and contractile impairment from 12 months onward. Molecularly, this progression was preceded by early upregulation of CCL2, a chemoattractant for cells expressing its receptor CCR2, along with increased IL-1β signaling in *Pkp2^+/−^* hearts. These changes promoted the recruitment of Ly6C⁺ CCR2⁺ monocytes and, ultimately, pro-inflammatory macrophages. These events parallel findings in *Dsg2*-based ACM models, where *Ccl2/Ccr2* upregulation and macrophage influx were identified as an early sign of the disease process^18,43^. In addition, we identified *Trem2^+^Spp1^+^* macrophages resembling the reparative *Trem2^hi^Spp1^hi^*population described after myocardial infarction (MI)^44^. Unlike the sequential inflammatory-to-reparative transition seen after MI, Ly6C^+^ CCR2^+^ monocytes and *Trem2^+^Spp1^+^*macrophages coexisted in *Pkp2^+/−^* hearts, indicating persistent low-grade signaling that sustains both inflammatory and pro-fibrotic programs. In a model of acute viral myocarditis, *Spp1^+^* macrophages instead adopted a pro-inflammatory role by inducing fibroblast *Ccl2/Ccl7* expression and promoting CCR2^+^ cell recruitment^45^. These findings highlight the environment-dependent heterogeneity of macrophage function in acute and chronic settings. At 9 months, apart from pro-inflammatory molecules, several immunoregulatory mediators were elevated, including CCL22, which attracts regulatory T cells^46^; PD-L2, a ligand of the PD-1 checkpoint receptor that dampens T-cell activation^47^; and FGF21, a cytokine with cardioprotective effects in mice limiting cardiac damage in hypertension and oxidative stress^27,28^. These changes suggest an early balance between pro- and anti-inflammatory signals. With age, this equilibrium shifted toward chronic inflammation, reflected by CCL12 expression at 15 months and also in *Pkp2^+/−^* MCMV hearts at 6mpi. A similar *Ccl12* induction has been reported in *Dsg2^cKO^* mice during late stages of disease, consistent with a shared inflammatory program driving CCR2⁺ cell recruitment^43^. Moreover, in the setting of MI combined with chronic periodontal inflammation, *Ccl12* was shown to prolong inflammation and impair cardiac wound healing, underscoring its role in sustaining maladaptive immune responses^48^. Kinome profiling at 12 months revealed concurrent activation of immune-cell kinase networks and stress-response pathways. Upregulated kinases included IKBKE and PRKD1, both activators of NF-κB^33,34^, and FGR, FES and LCK, linked to myeloid and lymphoid signaling^49–51^, together indicating sustained innate immune activation. Correspondingly, phosphorylated p65 and total p65 were elevated in 15-month-old *Pkp2^+/−^* hearts, confirming persistent NF-κB activity. While NF-κB-mediated monocyte recruitment has been described in *Dsg2^mut/mut^* models^52^, our data extend this mechanism to heterozygous *Pkp2* deficiency. Interestingly, Pérez-Hernández et al. also identified NF-κB as a transcription factor associated with PKP2, implying that PKP2 deficiency makes cardiomyocytes more prone to sterile myocarditis, which can be worsened by inflammatory or viral triggers^53^, as shown in our study. Integrating these findings, we propose that the *Pkp2^+/−^* genotype intrinsically activates NF-κB-dependent signaling in cardiomyocytes, driving sustained release of CCL2 and, at later stages, CCL12. This chemokine profile promotes recruitment of CCR2^+^ monocytes and impairs resolution of inflammation, resulting in the persistence of macrophages with both inflammatory and pro-fibrotic properties. Such inflammatory activation was absent in *Ttn^+/−^*hearts, underscoring the genotype-specific nature of this response. In *Pkp2^+/−^*hearts, sustained macrophage activity likely drives progressive injury by releasing proteases and reactive oxygen species^54^, which promote extracellular matrix remodeling and systolic dysfunction. Identifying this inflammatory phase before irreversible fibrosis develops defines a promising window for therapeutic intervention.

Building on this CCR2-centered mechanism, studies in *Dsg2^mut/mut^*-based ACM have demonstrated that genetically blocking CCR2-dependent monocyte recruitment reduces macrophage-driven fibrosis^52^, supporting the idea that CCR2^+^ cells directly contribute to desmosomal injury. In our *Pkp2^−/−^* model, CCR2^+^ macrophages were similarly the predominant expanded population, and short-term anti-CCR2 treatment showed effective depletion *in vivo*. Even though the intervention was initiated when fibrotic changes were already present, the reduction in CCR2^+^ myeloid cells suggests that blocking this pathway at an earlier, inflammation-dominated stage would probably exert a more pronounced effect - consistent with findings from the *Dsg2^mut/mut^* studies. However, sustained depletion of CCR2^+^ cells is not feasible in human patients because of their critical role in immune defense. Thus, although CCR2^+^ monocytes represent a shared disease-driving population across desmosomal cardiomyopathy models, these findings suggest that precisely timed modulation of early inflammatory signaling, rather than global depletion of myeloid cells, is the more promising therapeutic strategy.

Currently, no approved disease-specific targeted therapy exists for ACM, and anti-inflammatory or immunosuppressive agents such as glucocorticoids (e.g., dexamethasone) and azathioprine are used only occasionally and off-label in symptomatic patients^5,55,56^. Such treatments are likely to be most beneficial when applied during early or acute inflammatory disease phases, before irreversible remodeling develops.

More targeted approaches, including IL-1β antagonists, have shown promising results in related inflammatory cardiac conditions^57,58^. In *Pkp2^+/−^* mice, the therapeutic window spans 9-12 months, whereas in patients, the optimal timing remains to be determined and may vary between individuals. Our data align with retrospective clinical studies suggesting that anti-inflammatory therapies could benefit desmoplakin mutation carriers presenting with myocarditis-like symptoms^57,58^. Beyond therapeutic modulation, our findings highlight the influence of secondary triggers on the course of genetic cardiomyopathies. The absence of a cardiac phenotype following MCMV infection in *Ttn^+/−^* mice suggests that the timing and sequence of second hits are critical for disease manifestation. In humans, *TTN* variants are frequently detected in patients with complicated myocarditis^7^, which may reflect a scenario in which pre-existing, subclinical myocardial impairment serves as a first hit that limits recovery after infection and accelerates cardiomyopathy progression. In contrast, the viral challenge in our mouse model, applied before structural changes developed, may have preceded this window of susceptibility. It is conceivable that combining stressors, such as pregnancy, alcohol abuse or toxic substances, with viral infection could unmask such vulnerability in *Ttn*-deficient hearts^10^. In *Pkp2*-deficient hearts, however, latent MCMV proved to be a potent trigger of cardiac dysfunction, a finding with direct translational relevance: patients with PKP2 or other desmosomal mutations may particularly benefit from avoiding physical exertion or promptly treating systemic infections. Preventive measures, including vaccination and early antiviral therapy, should therefore be applied for individuals with a genetic predisposition. Although no CMV vaccine is currently available, antiviral agents such as ganciclovir, routinely used in immunocompromised patients^59^, could be explored as adjuncts to anti-inflammatory treatment during early myocarditis-like episodes. In conclusion, our findings emphasize infection control and timely immunomodulation as promising strategies to delay or prevent disease progression in ACM.

This study also has limitations that should be considered when interpreting the results. As with any mouse model, the immune and inflammatory responses characterized here may not fully capture the complexity of human ACM, where genetic diversity, lifelong antigen exposure and comorbidities shape disease expression. In addition, only male mice were included in the MCMV-infected *Pkp2^+/−^*cohort, reflecting the known male predominance in ACM, but future work should examine whether sex-specific immune responses modify the disease trajectory. Our findings in *Ttn^+/−^* mice represent a single truncating variant located in the A-band and therefore do not exclude the possibility that other variants and location in titin and in particular additional environmental triggers could interact with inflammation differently. Finally, while CCR2^+^ monocytes emerged as a shared disease-driving population across desmosomal cardiomyopathy models, their essential role in human host defense limits the direct translational applicability of cell-depletion strategies.

Despite these considerations, the convergence of genetic, immunological, and virological evidence across models provides strong support for early inflammatory signaling as a central driver of desmosomal cardiomyopathy and highlights the importance of defining clinically meaningful windows for therapeutic intervention.

## 7. Acknowledgements

The authors thank Anne Kilinç, Susanne Schraut and Daniela Urlaub for their dedicated technical assistance with mouse scoring, genotyping, and experimental procedures; Arnhild Grothey for providing the viral stocks; and Berkan Arslan for performing and analyzing echocardiographic measurements. Alicia Bender and Jakob Weilbach are acknowledged for their support during tissue collection, Sonja Luckhardt for expert technical assistance with cytokine profiling, and Alanis Barbosa Gulde for conducting the kinase activity analyses. We acknowledge the contributions of service projects PS1 and PS2 within the Collaborative Research Centre 1525 for their support in conducting echocardiography and supporting CITE-seq and single-cell transcriptomic profiling, respectively. Schematic subfigures were created in BioRender; Gerull, B. (2025) https://BioRender.com/hkmxgxg.

## 10. Supplemental Material

Supplemental Methods

Supplemental Figures

Supplemental Figures

References^11,30,44,60–78^

## List of nonstandard abbreviations

ARVC/ACM: Arrhythmogenic right ventricular cardiomyopathy/Arrhythmogenic cardiomyopathy
PKP2: plakophilin-2
DCM: Dilated cardiomyopathy
TTN: titin
MCMV: Murine cytomegalovirus
CCL2: C-C motif chemokine ligand 2
CCR2: C-C chemokine receptor type 2

