## Supplemental Material for "Murine CMV infection unmasks macrophage-driven inflammatory cardiomyopathy in *Pkp2*, but not in *Ttn* mutant mice"

### A. Supplemental Methods

#### Study approval

All animal studies and animal numbers used were in compliance with the Directive 2010/63/EU of the European Parliament, approved by the local ethics committee (RUF-55.2.2-2532-2-663, RUF-55.2.2-2532-2-962, and RUF-55.2.2-2532-2-1244) and conducted in accordance with the institutional guidelines.

#### Mouse models and housing conditions

C57BL/6J.129.Pkp2$^{tm1Mdcb}$ (*Pkp2$^{fl/fl}$*) mice were generated on a C57BL/6J genetic background as previously described[60] and kindly provided by Dr. Arnd Heuser (MDC, Berlin, Germany). In brief, a targeting vector was designed to introduce a *loxP* site upstream of exon 4 of the *Pkp2* gene and a *frt*-flanked neomycin (neo) resistance cassette together with a second *loxP* site downstream of exon 4. The construct, derived from the RP23-463A2 BAC clone, was linearized and electroporated into 129/Ola embryonic stem cells (ES). Clones with correctly targeted alleles were selected, expanded and injected into blastocysts from C57BL/6J mice. The neomycin cassette was selectively removed by crossing founder mice with FLP recombinase-expressing mice (B6;SJL-Tg(ACTFLPe)9205Dym/J; JAX stock #003800), resulting in *Pkp2* floxed alleles. Heterozygous *Pkp2$^{fl/-}$* mice were backcrossed to C57BL/6J for nine generations to establish a stable colony and then intercrossed to attain homozygosity (*Pkp2$^{fl/fl}$*). To achieve cardiac-specific deletion of exon 4, subsequent frame-shift and a premature stop codon, *Pkp2$^{fl/fl}$* mice were mated with C57BL/6J.αMHC-Cre mice (JAX stock #011038), which express Cre under the control of the cardiac α myosin heavy chain promoter, producing heterozygous *Pkp2$^{+/-}$*animals. These were crossed with the parental *Pkp2$^{fl/fl}$* strain to obtain homozygous *Pkp2$^{-/-}$* mice.

Generation of heterozygous C57BL/Cg-Ttn$^{tm1Brge}$ mice with a 2bp insertion causing a frameshift and premature stop codon (*Ttn$^{+/-}$*) were previously described[11] and maintained on a C57BL/6J background by breeding hemizygote carriers with noncarrier littermates.

Mice of both sexes were housed in groups under standard laboratory conditions in temperature- and humidity-controlled rooms. Animals had *ad libitum* access to standard rodent chow and water. A 12-hour light/dark cycle was maintained to reflect the natural day-night rhythm. To minimize stress and promote well-being, all cages were equipped with bedding, nesting material and an environmental enrichment structure. Experiments were conducted in *Pkp2$^{+/-}$*, *Pkp2$^{-/-}$* and Cre-negative *Pkp2$^{fl/fl}$* or *Pkp2$^{fl/-}$* mice or *Ttn$^{+/-}$* and wildtype control mice, with age and sex distributions as specified in the corresponding figure legends and results section. Whenever possible, littermate animals were used; otherwise, age-matched animals from different litters were included.

**Mouse genotyping**

Genotyping was conducted using PCR with genomic DNA extracted from ear punches and the following primers:

| | |
|---|---|
| Pkp2_fwd: | 5'- CTG ACC TGT GGG TAG AGT GGA -3' |
| Pkp2_rev: | 5'- GGA GAC AGG GAG ATA CCA ACC -3' |
| MHC-Cre_fwd | 5'- ATG ACA GAC AGA TCC CTC CTA TCT CC -3' |
| MHC-Cre_rev: | 5'- CTC ATC ACT CGT TGC ATC ATC GAC -3' |
| Ttn-WT_fwd | 5'- CAT TCG ACC ACC AAG CGA AAC ATC -3' |
| Ttn-WT_rev | 5'- ATA TCA CGG GAT GCC AAC GCT ATG -3' |
| Ttn-tg_fwd: | 5'- GAT GAC CAT CGA AAA CCC AGC -3' |
| Ttn-tg_rev: | 5'- GAC TAG CTT CTG ATA GAT GCC ACG -3' |

The wildtype (WT) and floxed *Pkp2* PCR products correspond to 257bp and 357bp, respectively; the αMHC-Cre PCR product to 300bp; and *Ttn* WT and transgene PCR products correspond to 289bp and 646bp, respectively.

**Echocardiography**

Cardiac function was assessed using high-resolution transthoracic echocardiography with two different protocols. Anesthesia was induced with inhaled isoflurane (3%) in 97% oxygen at a flow rate of 1l/min and maintained at 1–2%, depending on respiration rate. Mice were placed supine on a heated platform with embedded electrodes to monitor ECG and heart rate and to keep body temperature between 36.0-37.5°C throughout the procedure. Echocardiography was performed using a Vevo 1100 micro-ultrasound system (VisualSonics Inc.) equipped with 38MHz and 55MHz linear array transducers. In the first protocol, both two-dimensional B- and one-dimensional M-mode images were acquired in parasternal long- and short-axis views to assess cardiac morphology and function. Short-axis M-mode recordings at mid-papillary and apical levels of the left ventricle (LV) were used to calculate fractional shortening (FS) and to measure the thickness of the anterior, posterior, and free walls during diastole and systole. Additional B-mode images at maximum chamber diameter and different short-axis levels (base, mid-ventricle, and apex), along with end-diastolic and end-systolic lengths from the long-axis view, helped estimate LV ejection fraction (LVEF), cardiac output (CO), and LV volumes using Simpson's method. Four-chamber M-mode imaging at the tricuspid annulus was used to determine tricuspid annular plane systolic excursion (TAPSE). Pulsed-wave Doppler at the mitral valve in the apical four-chamber view was recorded to assess isovolumic contraction and relaxation times. Each parameter was averaged over at least three consecutive cardiac cycles recorded under stable anesthesia. The second protocol involved similar anesthesia, positioning, and monitoring. Two-dimensional B-mode imaging was performed in both parasternal long- and short-axis views, with guided M-mode tracings at the mid-ventricular level, distal to the posteromedial papillary muscle in the parasternal long-axis view for standard measurements of interventricular septum (IVS) thickness, LV internal diameter (LVID), and LV posterior wall (LVPW) thickness over 3-4 cardiac cycles. LV anterior wall (LVAW) thickness was measured from short-axis M-mode recordings as described previously[61]. LV volume and systolic function parameters, including FS, LVEF, stroke volume (SV) and CO, were obtained from a B-mode LV trace. Four-chamber views were used to record mitral valve flow velocity (E/A ratio) with pulse-wave Doppler and TAPSE in M-mode. To further evaluate right ventricular (RV) function, RV outflow tract (RVOT) recordings were performed and pulmonary valve size measured, followed by pulsed-wave Doppler of the pulmonary flow. To minimize motion artifacts, ECG and respiratory gating were applied during cine loop and M-mode acquisitions. Cine acquisition and analysis were performed blinded to genotype and experimental groups, using Vevo LAB software (version 3.2.0, VisualSonics Inc.).

**Δm157-MCMV-eGFP stock**

NIH-3T3 (ATCC CRL-1658) mouse embryonic fibroblasts were cultured in DMEM (Gibco) supplemented with 100IU/mL penicillin, 100µg/mL streptomycin, and 10% newborn calf serum (NCS). M2-10B4 (ATCC CRL-1972) fibroblasts were cultured in RPMI-1640 (Gibco) supplemented with 100IU/mL penicillin, 100µg/mL streptomycin and 10% fetal calf serum (FCS). All cells were maintained at 37°C in 5% $CO_2$. Δm157-MCMV-eGFP was previously generated[62] using en passant BAC mutagenesis[63]. Virus was reconstituted by transfection into NIH-3T3 cells using TransIT-X2 (Mirus). Crude virus supernatant was used to infect M2-10B4 cells to generate sucrose cushion-purified virus stocks, which were then titrated by standard plaque assays on NIH-3T3 cells as described previously[64].

**Δm157-MCMV-eGFP infection**

Mice were infected intraperitoneally (*i.p.*) with $5x10^5$ plaque-forming units (PFU) of cell culture-derived Δm157-MCMV-eGFP, while control mice received PBS. Groups of mice were infected and sacrificed at ages and days post-infection as indicated in the results and respective figure legends.

**$CCR2^+$ monocyte depletion**

To specifically inhibit the recruitment and infiltration of $CCR2^+$ monocytes into the heart, 3-week-old *Pkp2*$^{-/-}$ and Ctr mice received 20µg of a neutralizing anti-CCR2 monoclonal antibody (clone MC-21, kindly provided by Prof. Mack) *i.p.* for seven consecutive days[65]. Untreated mice of either genotype served as controls.

**Tissue collection**
After euthanasia and thoracotomy, EDTA whole blood was collected and stored at room temperature until further processing. Vascular perfusion was performed with PBS containing heparin (50IU/mL) to remove circulating leukocytes and flush the coronary vessels. Hearts were excised, arrested in diastole in ice-cold 37mM KCl in 0.9% NaCl, and divided into three transverse segments after the atria were removed. The base slice for histology was either frozen in optimal cutting temperature (OCT) compound or fixed in 4% formalin for paraffin embedding and stored at -20°C or 4°C, respectively. Mid-ventricular tissue for flow cytometry was immersed in RPMI-1640 (Gibco) and kept on ice until further processing. Apex tissue for biochemical and molecular analyses was snap-frozen in liquid nitrogen and stored at -80°C. For CITE sequencing, the atria and the valvular plane of the hearts were removed, and the remaining parts of the right and left ventricles were immersed in RPMI-1640 medium and kept on ice until preparation of the single-cell suspension.

**Histology**
Frozen hearts were transversely sectioned at 7µm thickness using a cryostat (Leica), while paraffin-embedded hearts were sliced into 10µm sections with a microtome (Leica). For assessment of overall cardiac morphology and fibrotic remodeling, deparaffinized sections were stained with hematoxylin and eosin[66] (HE; Morphisto) or picrosirius red (PSR; Morphisto, https://www.statlab.com/pdfs/ifu/KTPSR.pdf), respectively. Entire tissue cross-sections were imaged at high resolution using a Keyence microscope (BZ-X800 series). For quantification of collagen area fraction, ImageJ2 software[67] was used to analyze fibrotic areas in PSR-stained sections under polarized light as recently described[68].

**Immunohistochemistry (IHC)**
IHC staining was performed using the Avidin/Biotin Blocking and Vectastain® Elite-HRP ABC kits (Vector Labs) with minor modifications. In brief, 7µm cryosections were thawed at room temperature (RT), fixed in 4% formalin for 15min, washed twice in PBS, treated for 10min with 0.3% peroxide/methanol and for 2min with 50% ethanol, then washed twice with PBS. After blocking in 10% goat serum containing avidin (4drops/ml) for 1h and being washed twice with PBS, sections were incubated overnight at 4°C with the following primary antibodies: rabbit anti-CD4 (Abcam, clone 19514), rabbit anti-CD8 (Abcam, clone 21769), or rabbit anti–CD68 (Bosterbio, #PA1518), each diluted in 10% goat serum containing biotin (4drops/ml). For anti-CD45 (BioLegend, clone 30-F11) and anti-CD3 (BioLegend, clone 17A2) staining, sections were pretreated using the M.O.M.® Kit (Vector Labs) prior to antibody application. Following antibody incubation, slides were rinsed three times with PBS and subjected to biotinylated goat anti-rabbit IgG (Vector Labs, BA-1000-1.5) or goat anti-rat IgG (Vector Labs, BA-9401-.5) in 10% goat serum or M.O.M. ® diluent for 30min, respectively. After washing, sections were

incubated with ABC reagent, washed again, and then incubated in a 3,3'-diaminobenzidine (DAB) substrate solution (1drop/ml PBS). Afterwards, sections were counterstained with hematoxylin, cleared, and mounted. To validate antibody specificity, secondary antibody-only control sections were used. Entire tissue cross-sections were imaged at high resolution using a Keyence microscope (BZ-X800 series). Images were analyzed using a custom ImageJ2 macro that performed background correction, color deconvolution, and threshold-based segmentation to identify DAB-positive (brown) immune cells to quantify cell density per $mm^2$ tissue.

**Flow cytometry**

For preparation of cardiac single-cell suspension, pre-weighted perfused mid-ventricular tissue (approximately 50mg) was minced in 0.5ml RPMI-1640 (Gibco) and digested for 30min at 37°C with an equal volume of 4000IU/ml collagenase II in RPMI-1640 under agitation (600-1200rpm). The resulting suspension was passed through a 70µm cell strainer, washed with HBSS containing 1% BSA, and centrifuged (5min, 400g, 4°C). Cell pellets were resuspended in 300µl PBS and were divided for staining with antibody panels for monocyte/macrophage subsets (200µl) and T-cell subsets (100µl). For blood processing, 100µl of EDTA blood was mixed with 2ml of 1x RBC lysis buffer (BioLegend) and incubated for 10min at RT in the dark. After centrifugation (5min, 400g, 4°C), the cell pellet was resuspended in 200µl PBS and kept on ice until staining. Prior to antibody surface staining, cells were incubated with Zombie Aqua™ Fixable Viability dye (BioLegend) for 15min at RT in the dark to exclude dead cells later. Antibody surface staining was then performed with predefined antibody mixes for 15min at 4°C in the dark. After washing, cells were resuspended in 2% FCS in PBS and acquired on an Attune NxT Flow Cytometer (Thermo Fisher) and analyzed using FlowJo™ v10.8 (BD Life Sciences).

*Antibody mix for identification of cardiac monocyte and macrophage subsets:* either CD45-A700 (BioLegend, clone I3/2.3), CD11b-APC/Fire750 (BioLegend, clone 101262), Ly6G-Pacific Blue (BioLegend, clone 127612), MHC-II-A488 (BioLegend, clone M5/114.15.2), Ly6C-PerCP/Cy5 (BioLegend, clone HK1.4), TIM4-PE (BioLegend, clone RMT4.54) and F4/80-PE/Cy7 (BioLegend, clone BM8) or CD45-A700 (BioLegend, clone I3/2.3), CD11b-APC/Fire750 (BioLegend, clone 101262), CD88-PE/Cy7 (BioLegend, clone 135809), CD26-APC (BioLegend, clone 137807), Ly6G-PacificBlue (BioLegend, clone 127612), MHC-II-A488 (BioLegend, clone M5/114.15.2), Ly6C-PerCP/Cy5.5 (BioLegend, clone HK1.4), and TIM4-PE (BioLegend, clone RMT4.54).

*Antibody mix for identification of cardiac T-cell subsets:* CD45-BV421 (BioLegend, clone 103134), CD11b-A488 (BioLegend, clone 101217), CD8a-PerCP/Cy5.5 (BioLegend, clone

100734), CD4-PE (BioLegend, clone 100512), KLRG1-PE/Cy7 (BioLegend, clone 138416), TCRβ-A647 (BioLegend, clone 109218), Ly6G-A700 (BioLegend, clone 127622), and NK1.1-APC/Fire750 (BioLegend, clone 156516).

*Antibody mix for characterization of blood leukocytes:* CD45-A700 (BioLegend, clone I3/2.3), CD11b-A488 (BioLegend, clone M1/70), Ly6G-PacificBlue (BioLegend, clone 1A8), Ly6C-PerCP/Cy5 (BioLegend, clone HK1.4), and CD115-APC (BioLegend, clone AFS98).

To minimize nonspecific binding to Fc receptors, anti-CD16/32 (BioLegend, clone 62) was additionally added to each antibody mix.

**Quantitative real-time PCR of viral nucleic acid**

For detection and quantification of viral nucleic acid, DNA was isolated from snap frozen heart tissue using the DNeasy blood & Tissue kit with proteinase K digestion according to the manufacturer’s protocol (Qiagen, #69504). DNA concentration was determined by Nanodrop One (Thermo Scientific). 600ng of heart tissue DNA/animal was used to detect the MCMV-specific immediate early 1 gene (*ie1*). *Actb*, coding for β-actin, was chosen as reference. The quantitative transcription profile of *ie1* and *Actb* was determined by real-time PCR using the following primers/fluorogenic probe mix (Integrated DNA Technologies): $ie1_{fwd}$: 5´-GAG GAT CTT GTT GCG ACA T-3´ and $ie1_{rev}$: 5´-GAC ACA TTT CAT GAC TCT GCA TTT A-3´ with 6-FAM-probe: 5´- CTC TTG TTC TCT CAG GGT GCC AGC; β-$act_{fwd}$: 5´-AAA CCT AGA AGG TTG CTC TGA C-3´ and β-$act_{rev}$: 5´-GAC AGG ATG CAG AAG GAG ATT AC-3´ with Cy5-probe: 5´-AGC ATC CTT AGC TTG GTG AGG GT-3´. Reactions were set up in duplicates for each gene and run in a CFX96 Real-Time PCR Detection System (BioRad Technologies). Viral load was interpreted from quantification cycle values ($C_q$).

**Western blot**

Cardiac tissue homogenates were prepared in lysis buffer consisting of 50% (v/v) 2x Tris-Saline-EDTA (100mM Tris-HCl; 600mM NaCl, 10mM EDTA, 0.02% (m/v) $NaN_3$, pH 7.4), 10% (v/v) IBX phosphatase inhibitor buffer (500mM NaF; 50mM $Na_4P_2O_7$; 1mM $Na_3VO_4$; 0.02% (m/v) $NaN_3$), 1% (v/v) Triton-X-100, 2% (v/v) 50x EDTA-free Protease-Inhibitor Cocktail (Roche #11873580001), and 1% (v/v) 100x PMSF (100mM) using the Precellys 24 bead-based homogenizer according to the manufacturer’s protocol (Bertin Technologies). Protein concentration was determined using the Pierce™ BCA Protein Assay kit according to the manufacturer´s instructions (Thermo Scientific™, #23227). 10µg protein were mixed with 4x SDS sample buffer (BioRad Technologies, #161-0747) containing β-mercaptoethanol (Gibco™, #31350-010), boiled at 95°C for 5min, and subjected to SDS-PAGE on AnykD precast PAA gels (BioRad Technologies, #456-9036). Proteins were transferred onto nitrocellulose membranes (BioRad Technologies, #1704159), blocked in 5% BSA in TBST and immunoblotted overnight at 4°C with the primary antibodies PKP2 (Progen, #651101), GAPDH (Abcam, ab125247), phospho-p65 (Ser536, Cell Signaling, #3031), p65 (Abcam, #ab16502), β-Actin (Santa Cruz, #sc-81178). Subsequently, membranes were incubated with the respective HRP-conjugated anti-mouse (CST, #7076) or anti-rabbit (CST, #7074) antibodies for 1h at RT. For visualization, Amersham™ ECL western blotting detection reagents were used according to the manufacturer's protocol (GE Healthcare, #RPN2106). ECL signals were visualized with the ChemiDoc™ Touch imaging system (BioRad). Relative protein expression levels were quantified by densitometry and were normalized to the housekeeping proteins GAPDH or β-Actin using Image Lab 6.1 Software (BioRad, Version 6.1).).

**Kinome analysis**

Fresh frozen cardiac tissue samples were lysed for 30min in M-PER Mammalian Protein Extraction reagent (Thermo Scientific, #78503) supplemented with phosphatase (Thermo Scientific, #78420) and protease inhibitor cocktails (Thermo Scientific, #87785) at 1:80 dilutions, respectively. To remove debris, the lysates were centrifuged at 17.500xg for 15min at 4°C. Supernatants were collected, and their protein concentration was determined using the Pierce BCA Protein Assay Kit (Thermo Scientific, #23227) according to the manufacturer's instructions. Samples were aliquoted at 15µl each, snap-frozen in liquid nitrogen and stored at -80°C until further processing. Kinase activity profile analysis was performed at the Institute for Experimental Dermatology of the University Luebeck (https://www.lied.uni-luebeck.de/home). In brief, kinome activity profiling was performed using the PamGene PamChip® 4 protein tyrosine kinase (PTK) and serine/threonine kinase (STK) arrays (PamGene International B.V.) according to the manufacturer's instructions. Fluorescence signals corresponding to phosphorylation of phosphosites by protein kinases were monitored using Evolve software (PamGene International B.V.). Image quantification and data normalization were performed with BioNavigator software (PamGene International B.V.) as previously described[69]. Kinase activity scores were derived from phosphosite phosphorylation profiles. Kinases were considered regulated (either activated or inhibited) if the mean specificity score (the negative decadic logarithm of the likelihood of observing a higher difference between groups by random peptide-to-kinase assignment) was ≥1, and the significance score (the likelihood of observing a higher difference by random assignment of samples to treatment and control) was ≥0.5. The kinase statistic, calculated from the peptide statistics of the peptides phosphorylated by a specific kinase, indicates the directionality of the regulatory effect. When comparing *Pkp2*$^{+/-}$ to Ctr mice, values <0 suggest inhibition, while values >0 indicate activation in mutant versus control. Regulated kinases were subjected to network and pathway enrichment analyses using the STRING database[70]. Additionally, human homologues of the regulated murine kinases were superimposed on a kinome tree for visualization using Coral[71].

**Cytokine analysis**

Tissue homogenates for cytokine analysis were prepared from fresh frozen cardiac tissue lysed in RIPA lysis buffer (Millipore, #20-188) containing EDTA-free protease inhibitors (Roche, #118735800001) using the Precellys 24 bead-based homogenizer according to the manufacturer's instructions (Bertin Technologies). The Pierce BCA Protein Assay Kit (Thermo Scientific, #23227) was used to determine protein concentrations in duplicate. 20µg of protein were diluted to 1µg/µl in RIPA buffer. Protein profiles were measured at the Fraunhofer Institute for Translational Medicine and Pharmacology in Frankfurt am Main, Germany using Olink® Target 48 Mouse Cytokine assay (Thermo Fisher Scientific) according to the manufacturer´s instructions, capturing 43 proteins by high-throughput proximity extension assay (PEA) technology. Absolute cytokine concentrations (pg/ml) were calculated based on standard curves from internal calibration samples. Comparative barplot visualizations were generated in R (v4.5.1)[72] .

**Sample processing for single-cell RNA and CITE-seq**

To perform cellular indexing of transcriptomes and epitopes by sequencing[73] (CITE-seq) of ventricular $CD45^+$ leukocytes, 2.5µg of anti-CD45.2 APC antibody (BioLegend, clone 104) was injected intravenously into the tail vein prior to euthanasia to label all circulating immune cells. After thoracotomy, hearts were perfused with PBS containing heparin (50IU/mL) and excised. Atria and the heart base were removed and prepared for histology, while the remaining tissue was weighed and processed as recently described[44] with minor modifications. In brief, tissue was digested in RPMI-1640 containing 450U/ml collagenase I (Sigma-Aldrich, #C0130), 125U/ml collagenase XI (Sigma-Aldrich, C7657), 60U/ml hyaluronidase (Sigma-Aldrich, H3506), and 60U/ml DNase (Roche, #11284932001) under agitation at 37°C. After 45min, ~10% of the cell suspension was labeled with eBioscience™ Fixable Viability Dye eFluor™ 780 (ThermoFisher, #65-0865-14), CD45 PECy7 (BioLegend, Clone 30-F11), and CD3 PE (BD Biosciences, clone 17A2). Absolute cell counts of viable cardiac immune cells were assessed by flow cytometry using counting beads (BioLegend #424902), and normalized to the weight of the processed heart sample to obtain $CD45^+$ cells/mg tissue. The remaining 90% of the single cell suspension was washed twice in MACS buffer (PBS, 0.5% (m/v) BSA, 2mM EDTA) and was incubated on ice with TruStain FcX™ (anti-mouse CD16/32, BioLegend #101320) to block non-specific binding of immunoglobulins to the Fc receptors. After 5min, cells were stained with MicroBeads conjugated to monoclonal CD45 antibody (Miltenyi, clone 30F11.1 #130-052-301) and anti-CD45.2 A488 (BioLegend, clone 104) for 10min at 4°C. Subsequently, samples were incubated for another 15min at 4°C with a panel of 10 TotalSeq-A hashtag antibodies (BioLegend) to enable sample multiplexing[74] (see Supplemental Methods Table 1). In each experiment, 10 samples were pooled for single-cell analysis of either the *Pkp2*$^{+/-}$ model

(Ctr non-infected n=2, *Pkp2*$^{+/-}$ non-infected n=2, Ctr MCMV n=3 and *Pkp2*$^{+/-}$ MCMV n=3) or the *Ttn*$^{+/-}$ model (Ctr non-infected n=2, *Ttn*$^{+/-}$ non-infected n=2, Ctr MCMV n=3 and *Ttn*$^{+/-}$ MCMV n=3). In total, two independent *Pkp2*$^{+/-}$ experiments and one *Ttn*$^{+/-}$ experiment were performed. Following incubation, samples were washed twice in MACS buffer and were pooled before magnetic purification was performed using three separate LS Columns (Miltenyi, #130-042-401) and MidiMACS separators (Miltenyi, #130-042-302) to avoid clogging. Positive cell fractions were pooled, washed twice in 1% (w/v) BSA in PBS, and incubated in 1% (w/v) BSA in PBS containing eBioscienceTM Fixable Viability Dye eFluorTM 780 (ThermoFisher, #65-0865-14), anti-CD3-PE (BD Biosciences, clone 17A2), and anti-mouse TotalSeq-A Antibodies (all diluted at 1:400) used for surface marker detection during sequencing (see Supplemental Methods Table 2) for 25min at 4°C. Subsequently, cells were washed in 1% (w/v) BSA in PBS, resuspended, and viable CD45.2-APC(*i.v.*)$^{neg}$CD45-Alexa488$^{+}$ cells were sorted using a BD FACS Aria III with a 100µm nozzle. Finally, 23,000 sorted cells were loaded into the 10x Genomics Chromium in duplicate lanes for droplet encapsulation and barcoding using the Chromium Next GEM Single Cell 3′ Reagent Kit v3.1, aiming to recover 15,000 cells per lane.

| ***Pkp2*$^{+/-}$ cohort experiment 1** | | ***Pkp2*$^{+/-}$ cohort experiment 2** | | ***Ttn*$^{+/-}$ cohort experiment 1** | |
|---|---|---|---|---|---|
| Animal | Name | Group | Name | Group | Name |
| *Pkp2*$^{+/-}$ non-inf. 1 | Hashtag1_TotalA | *Pkp2*$^{+/-}$ non-inf. 3 | Hashtag1_TotalA | *Ttn*$^{+/-}$ non-inf. 1 | Hashtag1_TotalA |
| Ctr non-inf.1 | Hashtag2_TotalA | *Pkp2*$^{+/-}$ non-inf. 4 | Hashtag2_TotalA | *Ttn*$^{+/-}$ non-inf. 2 | Hashtag2_TotalA |
| Ctr MCMV 1 | Hashtag3_TotalA | Ctr MCMV 4 | Hashtag3_TotalA | Ctr MCMV 1 | Hashtag3_TotalA |
| Ctr MCMV 2 | Hashtag4_TotalA | Ctr MCMV 5 | Hashtag4_TotalA | *Ttn*$^{+/-}$ MCMV 1 | Hashtag4_TotalA |
| Ctr MCMV 3 | Hashtag5_TotalA | Ctr MCMV 6 | Hashtag5_TotalA | *Ttn*$^{+/-}$ MCMV 2 | Hashtag5_TotalA |
| *Pkp2*$^{+/-}$ MCMV 1 | Hashtag6_TotalA | Ctr non-inf.3 | Hashtag6_TotalA | Ctr non-inf.1 | Hashtag6_TotalA |
| *Pkp2*$^{+/-}$ MCMV 2 | Hashtag7_TotalA | Ctr non-inf.4 | Hashtag7_TotalA | Ctr non-inf.2 | Hashtag7_TotalA |
| *Pkp2*$^{+/-}$ MCMV 3 | Hashtag8_TotalA | *Pkp2*$^{+/-}$ MCMV 4 | Hashtag8_TotalA | Ctr MCMV 2 | Hashtag8_TotalA |
| Ctr non-inf.2 | Hashtag9_TotalA | *Pkp2*$^{+/-}$ MCMV 5 | Hashtag9_TotalA | *Ttn*$^{+/-}$ MCMV 3 | Hashtag9_TotalA |
| *Pkp2*$^{+/-}$ non-inf. 2 | Hashtag10_TotalA | *Pkp2*$^{+/-}$ MCMV 6 | Hashtag10_TotalA | Ctr MCMV 3 | Hashtag10_TotalA |

**Supplemental Methods Table 1** Hashtag-Sample relationship

| | | | | | |
|---|---|---|---|---|---|
| BTLA | CD152 | CD300cd | CD68 | IAIE | Podoplanin |
| CCR2 | CD163 | CD304 | CD80 | ICAM1 | Rat_IgG1 |
| CCR3 | CD169 | CD36 | CD81 | IgM | Rat_IgG2a |
| CCR5 | CD178 | CD39 | CD83 | IL1RL1 | Rat_IgG2b |
| CD103 | CD19 | CD4 | CD86 | ITGB7 | Sca1 |
| CD115 | CD21_35 | CD40 | CD88 | JAML | SiglecF |
| CD117 | CD223 | CD43 | CD8a | KLRG1 | SiglecH |
| CD11a | CD226 | CD44 | CD9 | Ly6C | Sirpa |
| CD11b | CD24 | CD47 | CD95 | Ly6G | TCRbeta |
| CD11c | CD25 | CD49d | CX3CR1 | Mac2 | TCRgd |
| CD127 | CD26 | CD5 | CXCR4 | MGL1 | TIGIT |
| CD135 | CD274 | CD55 | CXCR5 | MGL2 | TIM3 |
| CD137 | CD278 | CD62L | F480 | MSR1 | TIM4 |
| CD137L | CD279 | CD63 | FCeRIa | NK11 | XCR1 |
| CD14 | CD3 | CD64 | Hamster | PIRAB | |

**Supplemental Methods Table 2** CITE-seq marker panel

**Single-cell RNA and CITE-seq library preparation and sequencing**

Libraries were prepared using the Chromium Single Cell 3’ Reagents Kit v3 following the manufacturer’s instructions. Sample processing for CITE-seq and hashing was carried out according to the recommended protocol up to the cDNA amplification step, where additionally 1µl of ADT and HTO PCR primers for capturing antibody-derived tags (ADTs) and hashtag oligos (HTOs) were included, respectively. Following amplification, 0.6x SPRI beads (Beckman Coulter) were added to separate the ADT/HTO-derived cDNA (<180bp) containing supernatant from the bead-bound fraction harboring mRNA-derived cDNA (>300bp). While the mRNA-derived cDNA-containing bead fraction was handled according to the standard 10x Genomics protocol, the ADT-HTO-enriched supernatant underwent two 2x SPRI purifications, was split equally, and either amplified for ADTs or for HTOs. Indexing of ADT libraries was performed using TruSeq Small RNA primers, whereas hashtags were indexed with modified TruSeq DNA primers within the same PCR reactions[73,74]. After a final purification step using 1.6x SPRI beads, library concentrations were quantified with a Qubit™ 3.0 Fluorometer (Thermo Scientific) and quality assessed using an Agilent 2100 Bioanalyzer system and Agilent High Sensitivity DNA Kit reagents (Agilent, #5067-4626). Sequencing was performed on an Illumina NovaSeq 6000 platform using S1 or S2 flow cells with 100bp paired-end reads. Allocation of sequencing reads for CITE-seq and hashing libraries was set to approximately 85% for mRNAs, 10% for ADTs, and 5% for hashtags. Comprehensive protocols, including oligonucleotide sequences, are available at the CITE-seq resource website.

**Single-cell RNA and CITE-seq data analysis**

10x Genomics data, including HTO and ADT libraries, were demultiplexed using Cell Ranger software (v7.0.1) with the “intron mode” enabled, which incorporates intronic reads. For alignment and counting, the mouse reference genome GRCm38/mm10 was used. To simultaneously assess cell surface protein expression alongside the transcriptome in our main dataset, we used the -feature-ref flag in Cell Ranger. This generates a combined matrix containing both gene expression counts and the expression levels of cell surface proteins, which was further analyzed using Seurat (v5.3.0)[75]. For each experiment, two libraries were processed separately in the initial steps. In total, data were obtained from two *Pkp2* experiments and one *Ttn* experiment, each containing 10 hashed samples and 89 CITE-seq antibodies. RNA expression and antibody-derived tag (ADT) data were separated, and hashtag oligonucleotides (HTOs) were extracted from the antibody capture data. Separate Seurat objects were created for each library and log-normalized. An HTO assay was added and CLR-normalized, and HTODemux with an adjusted positive.quantile parameter was used to demultiplex HTOs and assign cells to samples; only singlets were retained. The ADT assay was likewise CLR-normalized. Quality control was performed per library, excluding cells with > 7.5% mitochondrial transcripts or with nCount_RNA > 30,000-40,000 (threshold depending on dataset). Potentially remaining doublets were identified based on transcription data and removed using DoubletFinder (v2.0.6)[76]. The resulting singlets comprised *Pkp2* Exp1Lib1: 6186 cells; *Pkp2* Exp1Lib2: 6633; *Pkp2* Exp2Lib1: 7063; *Pkp2* Exp2Lib2: 6774; *Ttn* Exp1Lib1: 6479; *Ttn* Exp1Lib2: 6181 cells. Datasets were pooled and normalized with SCTransform(), followed by principal component analysis using 25 components. Integration across experiments was performed using Harmony (v1.2.3)[77]. Louvain clustering was applied (resolution = 0.2, dimensions 1–25), yielding 16 clusters, and UMAP embedding was computed on 25 dimensions. Clusters were annotated based on surface proteins and differentially expressed genes. Myeloid cell subsets were re-clustered using 17 principal components and resolution=0.3 without recalculating UMAP, producing 13 clusters (two merged, one contaminant removed, final = 11). Lymphoid cells were similarly re-clustered using dimensions 1–16 and resolution=0.25, resulting in 12 clusters; a small contaminating population with myeloid and proliferative signatures was retained and labeled moMac_cycling_inflam. Refined subcluster annotations were merged back into the integrated dataset for the global refined annotation.

Neighborhood differential abundance was assessed using MiloR (v2.2.0)[30] on the integrated Seurat object (Refined_annotation). The object was converted to SingleCellExperiment, preserving Sample, Condition, and Refined_annotation metadata, with PCA and Harmony-corrected UMAP embeddings stored in reducedDims. A Milo object was created, and a KNN graph built with buildGraph(). Neighborhoods were defined using makeNhoods(), and cell

counts per sample obtained with countCells(). Differential abundance testing was performed with testNhoods() using the model ~Condition, comparing two groups (e.g. Pkp2_Ctr_MCMV vs Pkp2_Ctr_noninf). Significant neighborhoods were defined by SpatialFDR<0.1, annotated with annotateNhoods(), and labeled “Mixed” if annotation purity<0.7. SpatialFDR reports p-values corrected for multiple testing, taking into account neighborhood overlap. Results were visualized as UMAP-based neighborhood maps and beeswarm plots.
Cytokine and receptor expression was analyzed with a pre-defined panel of cytokine ligands and receptors (see Supplemental Methods Table 2). Differential expression was tested across experimental contrasts (e.g. Pkp2_HetKO_noninf vs Pkp2_Ctr_noninf) and was computed with FindMarkers (Wilcoxon test; logfc.threshold = 0, min.pct = 0.05), and genes with adj. p < 0.05 and |avg_log2FC| > 0.25 were considered significant. Results were visualized as volcano plots.

| | |
|---|---|
| Cytokine ligands | *Ccl2, Ccl4, Ccl5, Ccl11, Ccl12, Ccl17, Ccl22, Cxcl1, Cxcl2, Cxcl9, Cxcl12, Il1a, Il1b, Il2, Il3, Il4, Il5, Il6, Il7, Il9, Il10, Il12a, Il12b, Il16, Il17a, Il17f, Il21, Il22, Il27, Il31, Il33, Ifna2, Ifng, Ifnl2, Tnf, Csf1, Csf2, Csf3, Hgf, Fgf21* |
| Cytokine receptors | *Ccr1, Ccr2, Ccr3, Ccr4, Ccr5, Ccr8, Cxcr1, Cxcr2, Cxcr3, Cxcr4, Ackr3, Il1r1, Il1rap, Il1rl1, Il2ra, Il2rb, Il2rg, Il3ra, Il4ra, Il5ra, Il6ra, Il6st, Il7r, Il9r, Il10ra, Il10rb, Il12rb1, Il12rb2, Il17ra, Il17rc, Il21r, Il22ra1, Il27ra, Il31ra, Osmr, Ifnar1, Ifnar2, Ifnlr1, Ifngr1, Ifngr2, Tnfrsf1a, Tnfrsf1b, Csf1r, Csf2ra, Csf2rb, Csf3r, Cd4, Cd80, Cd86, Ctla4, Pdcd1, Pdcd1lg2, Cd274, Met, Fgfr1, Klb* |

**Supplemental Methods Table 3** Genes used for differential cytokine analysis

For pseudobulk analysis, raw RNA counts from the Refined_annotation Seurat object were aggregated by sample and cell type using AggregateExpression(). For each cell type, genes with fewer than 10 total counts were removed, and differential expression was tested with DESeq2. Genes with adjusted p < 0.05 were considered significant. Significantly up- and downregulated genes were analyzed with enrichR (v3.4)[78], querying the MSigDB Hallmark 2020 database. Enrichment results were ranked by adjusted p-value and Combined Score, a metric integrating the significance and magnitude of enrichment, and the top 15 pathways per cell type were visualized as bar plots.

#### Statistical analyses

Statistical significances were calculated using GraphPad Prism version 9.5.1 (for Windows, GraphPad Software, www.graphpad.com). Results are represented as mean ± SEM. For two-group comparisons, normality was assessed with the Anderson-Darling, D’Agostino-Pearson omnibus, and Shapiro-Wilk tests, while equal variances were evaluated using the F test. If the data met the assumptions of normality and equal variances, a parametric unpaired t test was used. When the assumption of equal variances was violated, Welch’s correction was applied. If any of the normality tests failed, the nonparametric Mann-Whitney test was used. For more than two groups, the same normality tests were performed, with Bartlett’s and Brown-Forsythe tests used to assess equal variances. When data were normally distributed, with or without equal variances, an ordinary one-way ANOVA with or without Brown-Forsythe and Welch's correction and Tukey's or Dunnett's T3 multiple comparisons were conducted. When data did not follow a normal distribution, the nonparametric Kruskal-Wallis test with Dunn's multiple comparisons was performed.

**Statistical analyses**

Statistical significances were calculated using GraphPad Prism version 9.5.1 (for Windows, GraphPad Software, www.graphpad.com). Results are represented as mean ± SEM. For two-group comparisons, normality was assessed with the Anderson-Darling, D'Agostino-Pearson omnibus, and Shapiro-Wilk tests, while equal variances were evaluated using the F test. If the data met the assumptions of normality and equal variances, a parametric unpaired t test was used. When the assumption of equal variances was violated, Welch's correction was applied. If any of the normality tests failed, the nonparametric Mann-Whitney test was used. For more than two groups, the same normality tests were performed, with Bartlett's and Brown-Forsythe tests used to assess equal variances. When data were normally distributed, with or without equal variances, an ordinary one-way ANOVA with or without Brown-Forsythe and Welch's correction and Tukey's or Dunnett's T3 multiple comparisons were conducted. When data did not follow normal distribution, the nonparametric Kruskal-Wallis test with Dunn's multiple comparisons was performed.

1 # B. Supplemental Figures

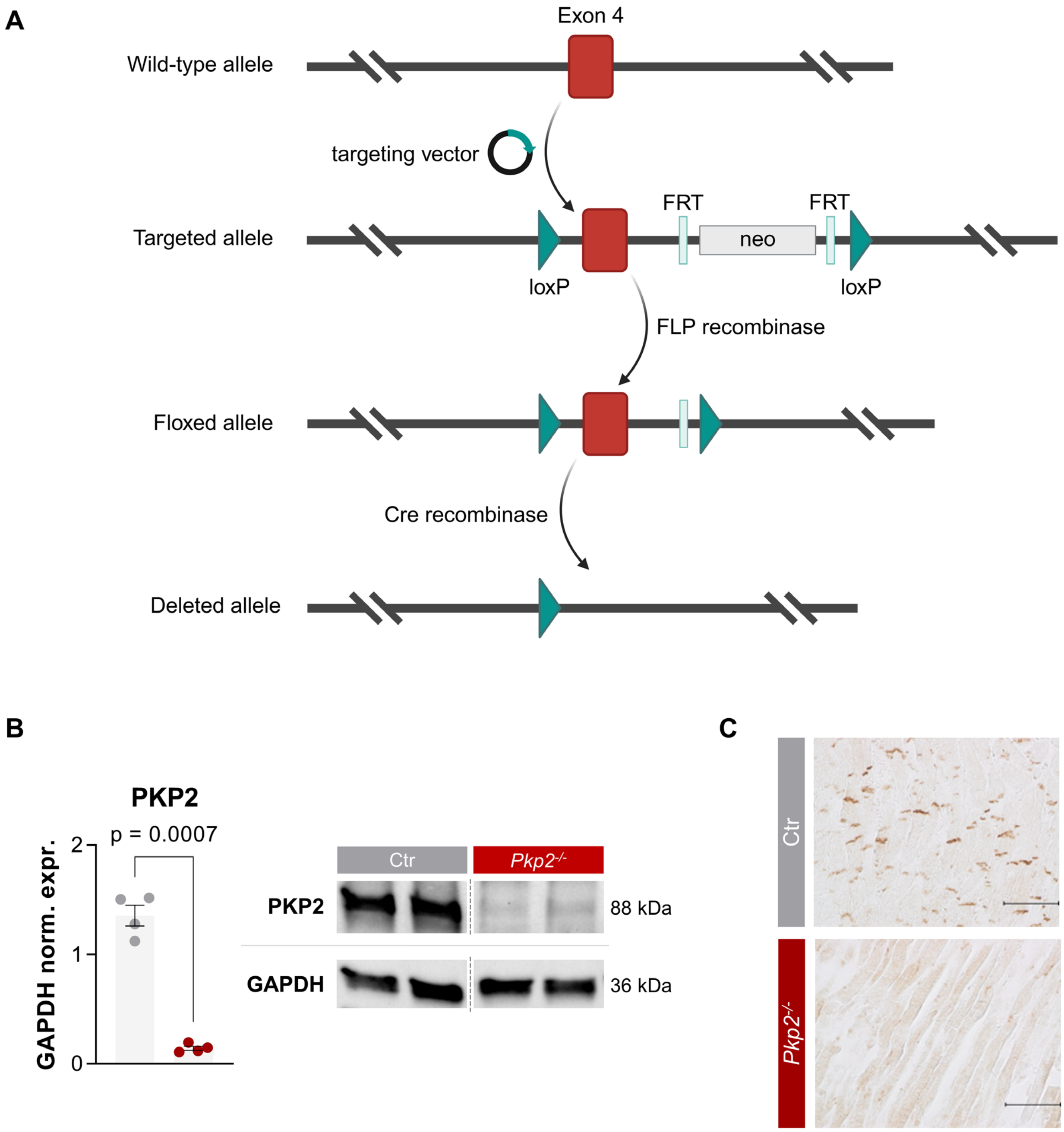

**Figure S1**

**A**, Schematic overview of the wildtype, targeted, floxed and deleted *Pkp2* allele. Created with Biorender.com **B**, Western blot quantification and representative blot snips PKP2 protein levels in 3-4-week-old *Pkp2*$^{-/-}$ hearts normalized to GAPDH levels; Ctr n = 4, *Pkp2*$^{-/-}$ n = 4. **C**, Representative images of PKP2 immunostaining in *Pkp2*$^{-/-}$ hearts; scale bar 50µm. Data are shown as mean ± SEM. Male: squares; female: circles. Unpaired t test with or without Welch's correction or Mann-Whitney test, as appropriate. Neo – neomycin cassette.

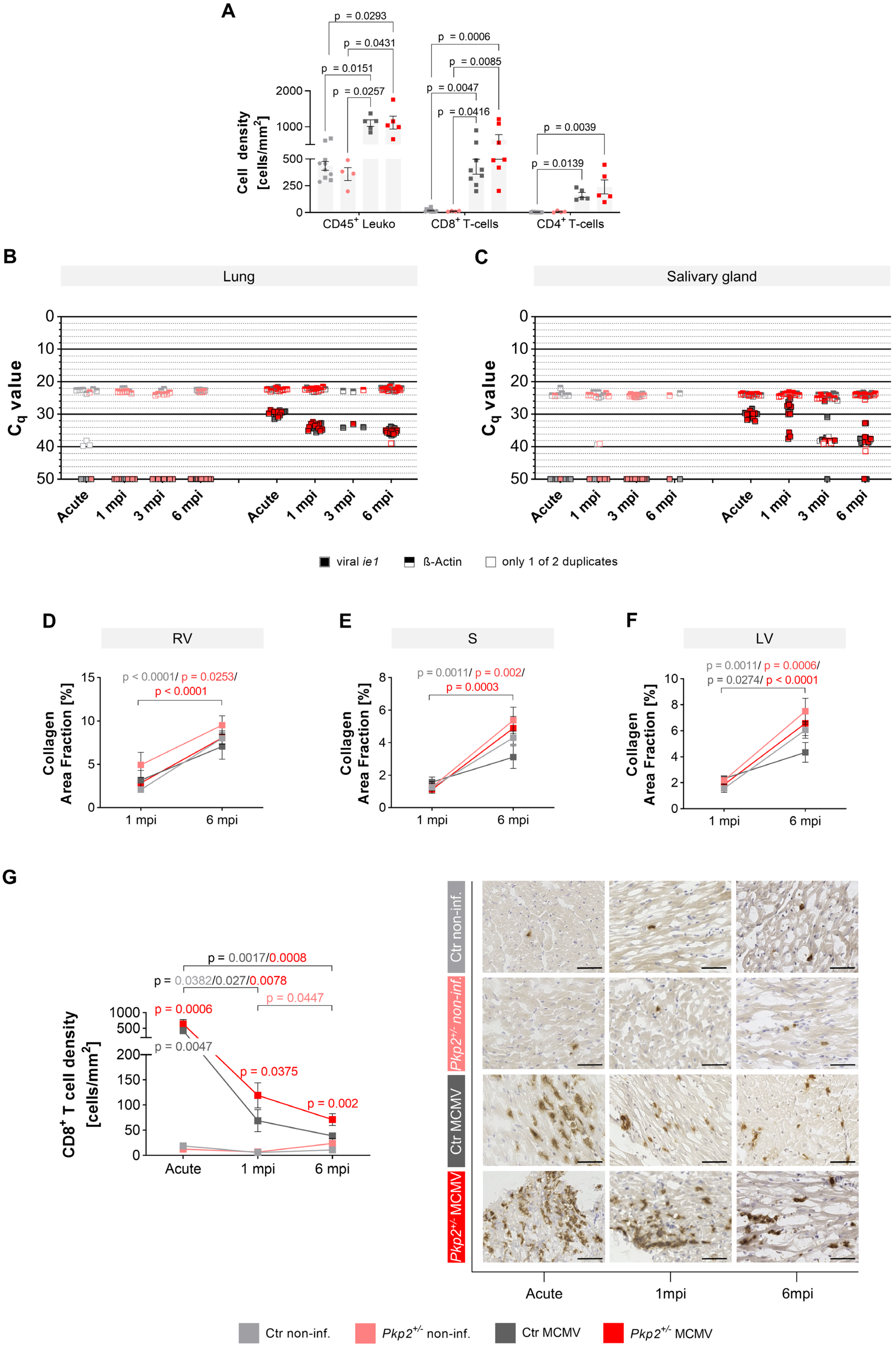

A
p = 0.0293
p = 0.0431
p = 0.0151
p = 0.0257
p = 0.0006
p = 0.0085
p = 0.0047
p = 0.0416
p = 0.0039
p = 0.0139
Cell density [cells/mm²]
2000
1000
500
250
0
CD45⁺ Leuko
CD8⁺ T-cells
CD4⁺ T-cells
B
Lung
C
Salivary gland
Cq value
0
10
20
30
40
50
Acute
1 mpi
3 mpi
6 mpi
viral ie1
ß-Actin
only 1 of 2 duplicates
D
RV
p < 0.0001/ p = 0.0253/
p < 0.0001
Collagen Area Fraction [%]
1 mpi
6 mpi
E
S
p = 0.0011/ p = 0.002/
p = 0.0003
F
LV
p = 0.0011/ p = 0.0006/
p = 0.0274/ p < 0.0001
G
p = 0.0017/0.0008
p = 0.0382/0.027/0.0078
p = 0.0447
p = 0.0006
p = 0.0047
p = 0.0375
p = 0.002
CD8⁺ T cell density [cells/mm²]
Acute
1 mpi
6 mpi
Ctr non-inf.
Pkp2+/- non-inf.
Ctr MCMV
Pkp2+/- MCMV
Acute
1mpi
6mpi
Ctr non-inf.
Pkp2+/- non-inf.
Ctr MCMV
Pkp2+/- MCMV

### Figure S2

**A**, Quantification of cell densities of $CD45^+$ leukocytes, $CD8^+$, and $CD4^+$ T-cells in cardiac sections from immunostaining during acute MCMV infection; n = $CD45^+$/$CD8^+$/ $CD4^+$; Ctr non-infected n = 10/11/10, *Pkp2*$^{+/-}$ non-infected n = 4/4/4, Ctr MCMV n = 4/9/4, *Pkp2*$^{+/-}$ MCMV n = 5/7/5. **B+C**, Viral *ie1* (*immediate early 1*, filled squares) and murine ß-actin (half-top-filled squares) DNA copies detected by quantitative polymerase-chain reaction in lung (B) and salivary gland (C) samples from acute to latent infection represented by quantification cycle values ($C_q$). Measurements were performed in technical duplicates. It is indicated by an unfilled symbol when only one value was usable; n(lung) = Acute/1mpi/3mpi/6mpi: Ctr non-infected n = 6/3/2/410, *Pkp2*$^{+/-}$ non-infected n = 1/7/6/10, Ctr MCMV n = 6/5/2/10, *Pkp2*$^{+/-}$ MCMV n = 8/10/1/10; n(salivary gland) = Acute/1mpi/3mpi/6mpi: Ctr non-infected n = 6/5/6/1, *Pkp2*$^{+/-}$ non-infected n = 1/5/9/1, Ctr MCMV n = 6/3/9/11, *Pkp2*$^{+/-}$ MCMV n = 8/9/7/9. **D-F**, Quantification of collagen area fraction in picrosirius red stained cardiac sections at 1 and 6mpi separately analyzed in the right ventricle (D), septum (E) and left ventricle (F). **G**, Representative images of $CD8^+$ immunostaining and quantification of $CD8^+$ T-cell density from acute to latent infection; n = Acute/1mpi/3mpi/6mpi: Ctr non-infected n = 11/4/11/8, *Pkp2*$^{+/-}$ non-infected n = 4/6/11/9, Ctr MCMV n = 9/5/9/5, *Pkp2*$^{+/-}$ MCMV n = 7/11/6/7. Data are shown as mean ± SEM. Only male mice were used (square symbol). Ordinary one-way ANOVA with or without Brown-Forsythe and Welch's correction and Tukey's or Dunnett's T3 multiple comparisons, respectively or Kruskal-Wallis test with Dunn's multiple comparisons, as appropriate for >2 groups. Unpaired t test with or without Welch's correction or Mann-Whitney test, as appropriate for 2 groups. P-values in G are shown for one group over time (bracket) and across groups at one time point compared to Ctr non-infected. All other p-values can be found in Table S3-S6. Leuko – leukocytes, RV – right ventricle, S – septum, LV – left ventricle.

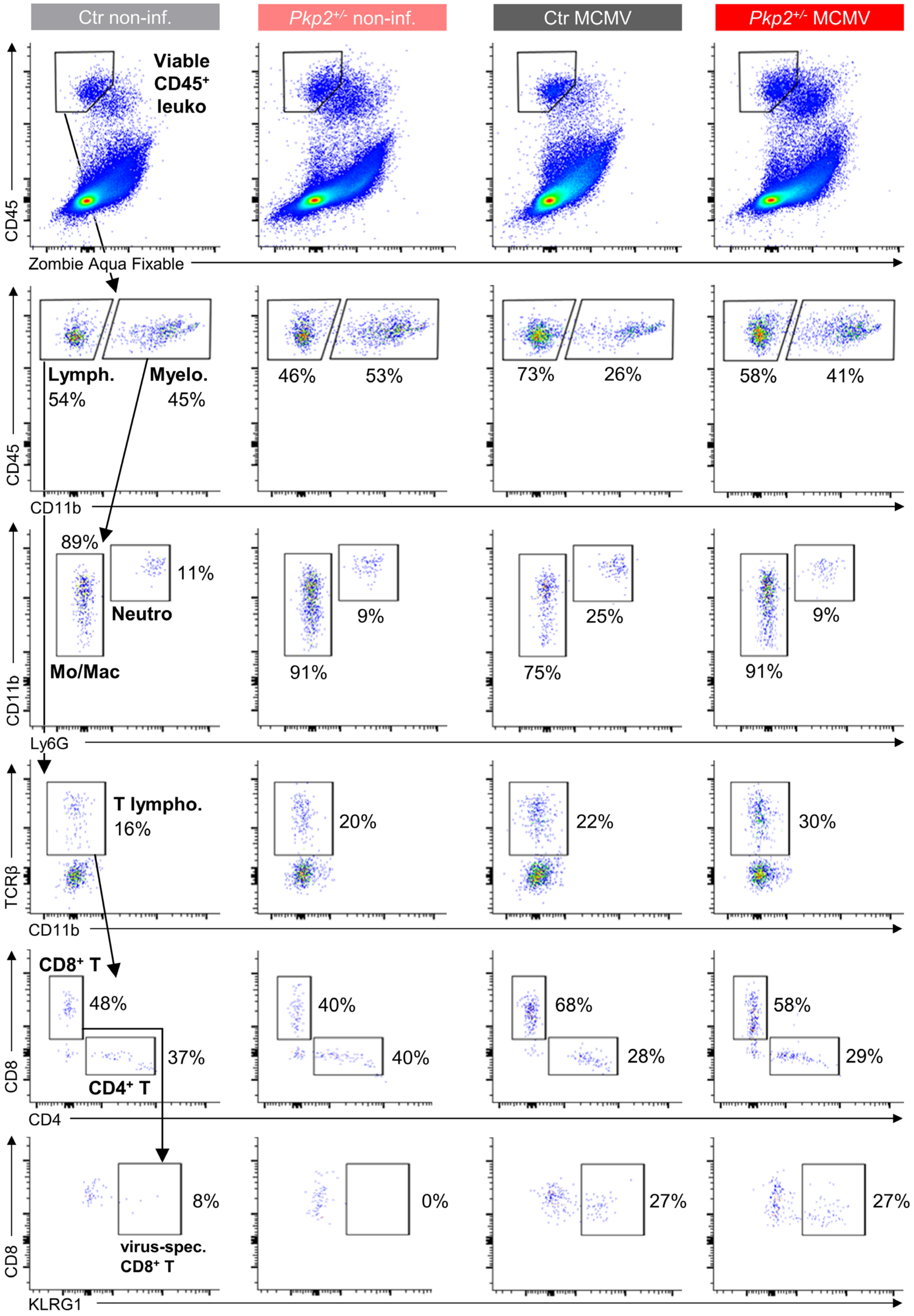
Ctr non-inf.
Pkp2+/- non-inf.
Ctr MCMV
Pkp2+/- MCMV
Viable CD45+ leuko
CD45
Zombie Aqua Fixable
Lymph. 54%
Myelo. 45%
46%
53%
73%
26%
58%
41%
CD11b
89%
11%
Neutro
Mo/Mac
9%
91%
25%
75%
9%
91%
Ly6G
T lympho. 16%
20%
22%
30%
TCRβ
CD8+ T
48%
37%
CD4+ T
40%
40%
68%
28%
58%
29%
CD8
CD4
8%
virus-spec. CD8+ T
0%
27%
27%
KLRG1

**Figure S3**

Representative flow cytometry gating of 6mpi heart samples using a T-cell tailored panel. Leuko – leukocytes, Neutro – neutrophils, Mo/Mac – monocytes/macrophages.

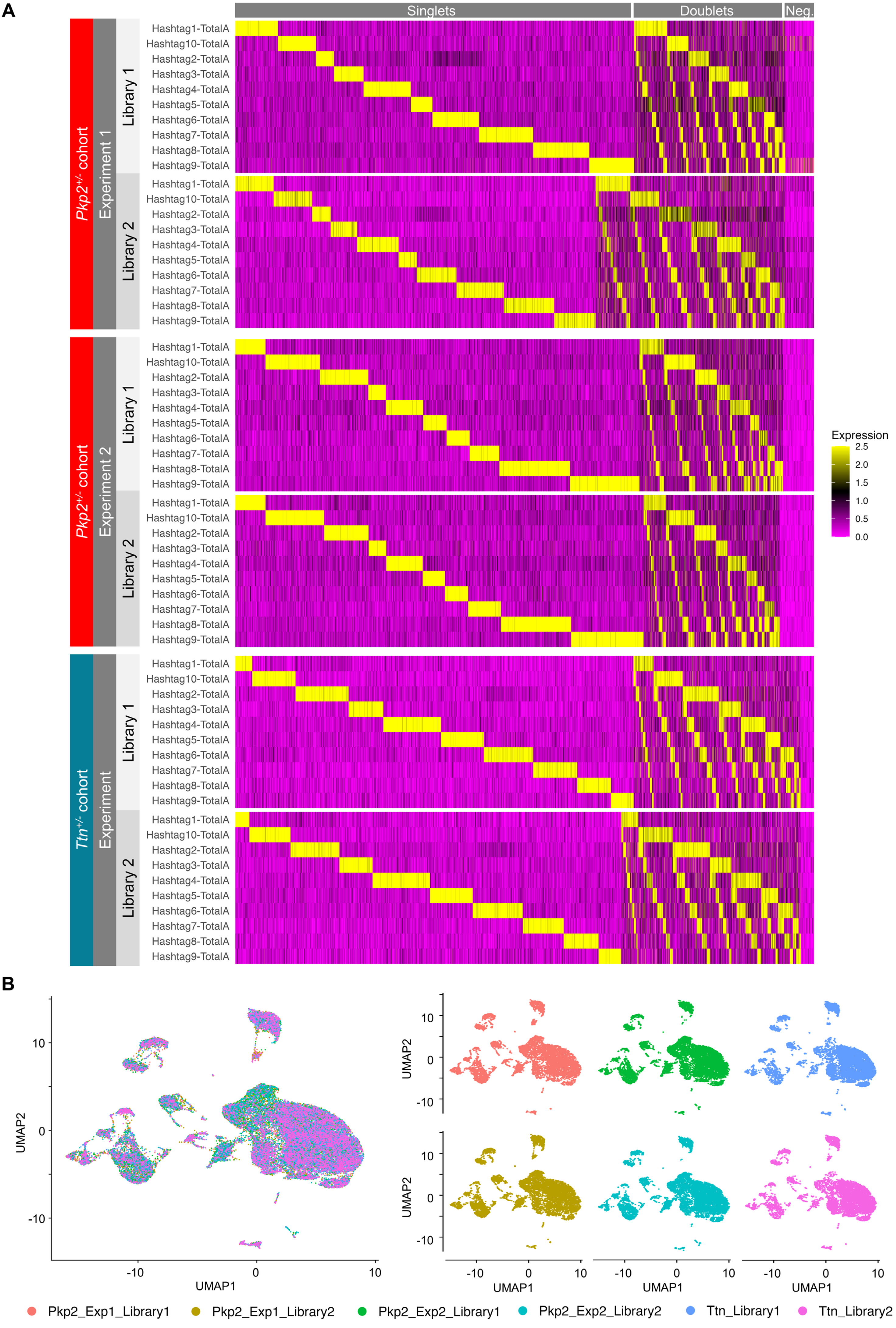
A
Singlets
Doublets
Neg.
Pkp2+/- cohort
Experiment 1
Library 1
Library 2
Pkp2+/- cohort
Experiment 2
Library 1
Library 2
Ttn+/- cohort
Experiment
Library 1
Library 2
Hashtag1-TotalA
Hashtag10-TotalA
Hashtag2-TotalA
Hashtag3-TotalA
Hashtag4-TotalA
Hashtag5-TotalA
Hashtag6-TotalA
Hashtag7-TotalA
Hashtag8-TotalA
Hashtag9-TotalA
Expression
2.5
2.0
1.5
1.0
0.5
0.0
B
UMAP1
UMAP2
10
0
-10
Pkp2_Exp1_Library1
Pkp2_Exp1_Library2
Pkp2_Exp2_Library1
Pkp2_Exp2_Library2
Ttn_Library1
Ttn_Library2

**Figure S4**

**A**, Heatmaps showing TotalSeq-A hashtag demultiplexing signatures for all single-cell RNA and CITE-sequencing experiments (*Pkp2*$^{+/-}$ and *Ttn*$^{+/-}$ cohorts) to distinguish singlets, doublets, and negative cells. A demultiplexing table correlating hashtags with experimental groups is available in the Supplemental Methods. **B**, UMAP grouped by library after data set integration with Harmony. Cells from different libraries are overlaid in the left figure. The right figure shows UMAPs separated by library. UMAP - uniform manifold approximation and projection.

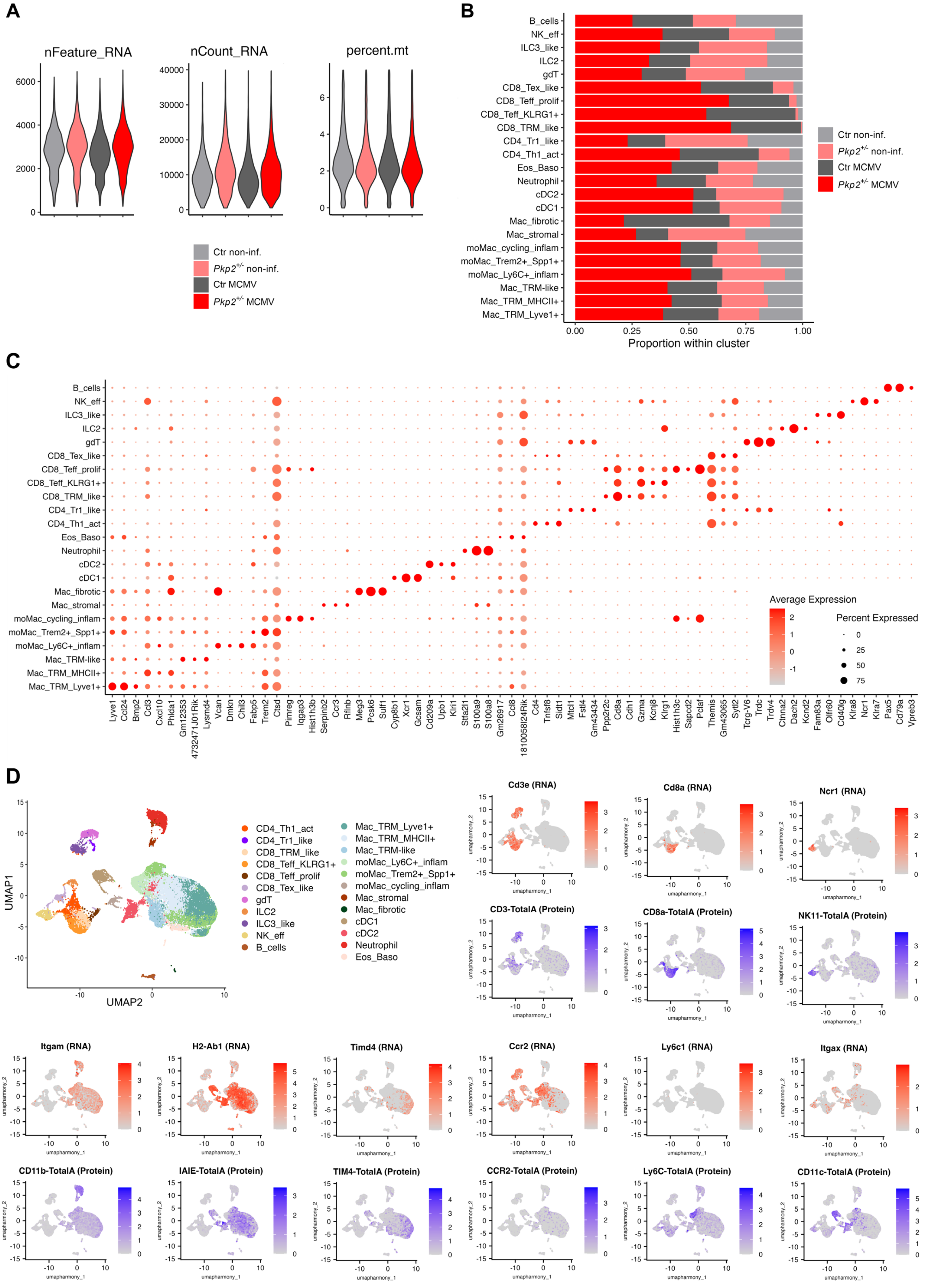
A
nFeature_RNA
nCount_RNA
percent.mt
Ctr non-inf.
Pkp2+/- non-inf.
Ctr MCMV
Pkp2+/- MCMV
B
B_cells
NK_eff
ILC3_like
ILC2
gdT
CD8_Tex_like
CD8_Teff_prolif
CD8_Teff_KLRG1+
CD8_TRM_like
CD4_Tr1_like
CD4_Th1_act
Eos_Baso
Neutrophil
cDC2
cDC1
Mac_fibrotic
Mac_stromal
moMac_cycling_inflam
moMac_Trem2+_Spp1+
moMac_Ly6C+_inflam
Mac_TRM-like
Mac_TRM_MHCII+
Mac_TRM_Lyve1+
Proportion within cluster
C
Average Expression
Percent Expressed
Lyve1 Ccl24 Bmp2 Ccl3 Cxcl10 Phlda1 Gm12353 4732471J01Rik Lysmd4 Vcan Dmkn Chil3 Fabp5 Trem2 Ctsd Pimreg Iqgap3 Hist1h3b Serpinb2 Ccr3 Rflnb Meg3 Pcsk6 Sulf1 Cyp8b1 Xcr1 Gcsam Cd209a Upb1 Klri1 Stfa2l1 S100a9 S100a8 Gm26917 Ccl8 1810058I24Rik Cd4 Tnfsf8 Sidt1 Mtcl1 Fstl4 Gm43434 Ppp2r2c Cd8a Cdh1 Gzma Kcnj8 Klrg1 Hist1h3c Sapcd2 Pclaf Themis Gm43065 Sytl2 Tcrg-V6 Trdc Trdv4 Ctnna2 Dach2 Kcnd2 Fam83a Olfr60 Cd40lg Klra8 Ncr1 Klra7 Pax5 Cd79a Vpreb3
D
UMAP1
UMAP2
CD4_Th1_act
CD4_Tr1_like
CD8_TRM_like
CD8_Teff_KLRG1+
CD8_Teff_prolif
CD8_Tex_like
gdT
ILC2
ILC3_like
NK_eff
B_cells
Mac_TRM_Lyve1+
Mac_TRM_MHCII+
Mac_TRM-like
moMac_Ly6C+_inflam
moMac_Trem2+_Spp1+
moMac_cycling_inflam
Mac_stromal
Mac_fibrotic
cDC1
cDC2
Neutrophil
Eos_Baso
Cd3e (RNA)
Cd8a (RNA)
Ncr1 (RNA)
CD3-TotalA (Protein)
CD8a-TotalA (Protein)
NK11-TotalA (Protein)
Itgam (RNA)
H2-Ab1 (RNA)
Timd4 (RNA)
Ccr2 (RNA)
Ly6c1 (RNA)
Itgax (RNA)
CD11b-TotalA (Protein)
IAIE-TotalA (Protein)
TIM4-TotalA (Protein)
CCR2-TotalA (Protein)
Ly6C-TotalA (Protein)
CD11c-TotalA (Protein)
umapharmony_1
umapharmony_2

**Figure S5**

**A**, Violin plots showing the distribution of nFeature_RNA, nCount_RNA, and percent.mt per cell after quality filtering. **B**, Bar plot depicting the relative contribution of each experimental group to the total number of cells within each identified cell type cluster. **C**, Dot plot displaying the top three differentially expressed genes per cluster based on RNA expression levels. **D**, FeaturePlots of selected surface proteins and their corresponding RNA transcripts, demonstrating expression across modalities. Annotated UMAP for cluster reference on the side. UMAP - uniform manifold approximation and projection.

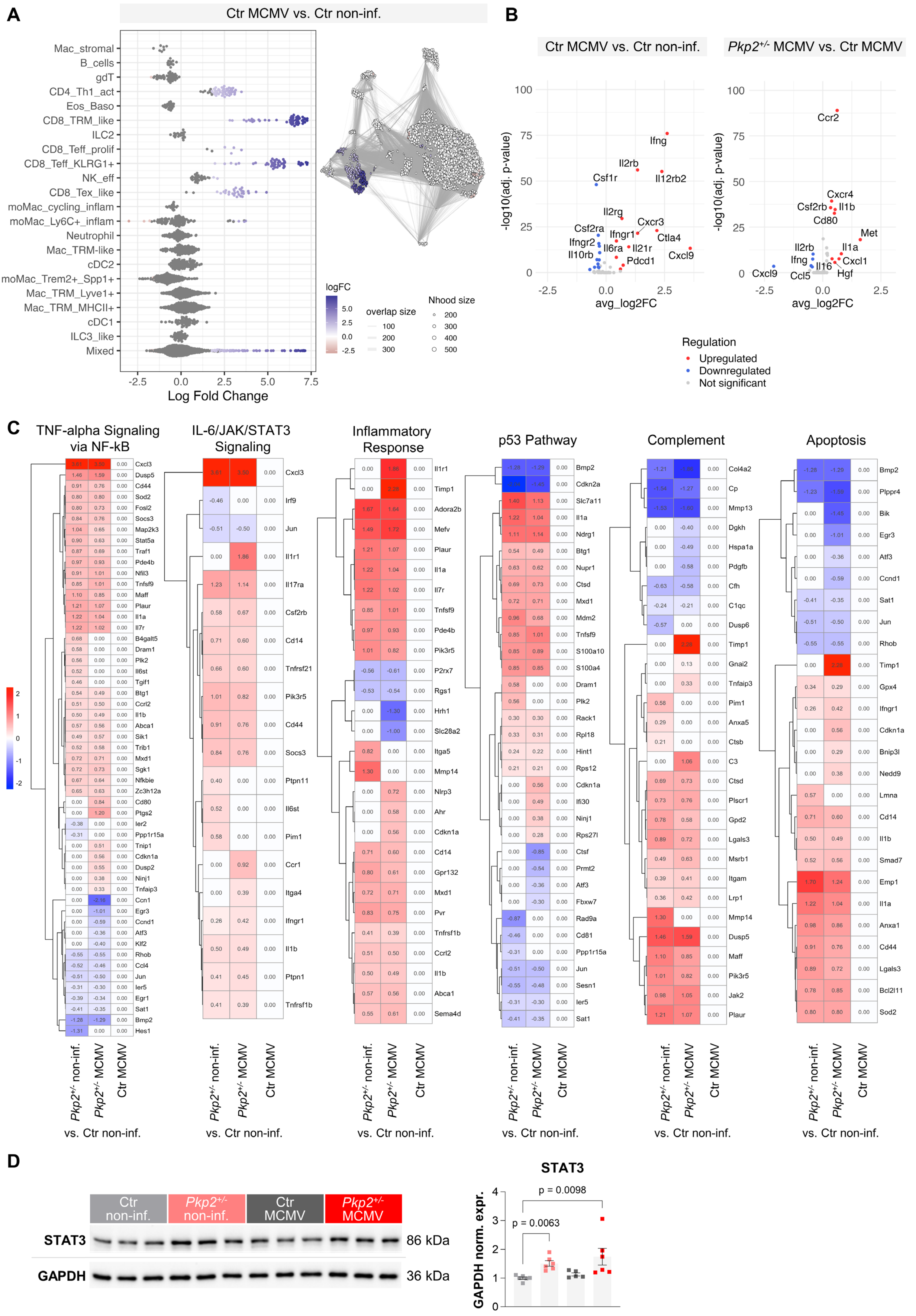
A
Ctr MCMV vs. Ctr non-inf.
Mac_stromal
B_cells
gdT
CD4_Th1_act
Eos_Baso
CD8_TRM_like
ILC2
CD8_Teff_prolif
CD8_Teff_KLRG1+
NK_eff
CD8_Tex_like
moMac_cycling_inflam
moMac_Ly6C+_inflam
Neutrophil
Mac_TRM-like
cDC2
moMac_Trem2+_Spp1+
Mac_TRM_Lyve1+
Mac_TRM_MHCII+
cDC1
ILC3_like
Mixed
-2.5
0.0
2.5
5.0
7.5
Log Fold Change
logFC
5.0
2.5
0.0
-2.5
overlap size
100
200
300
Nhood size
200
300
400
500
B
Ctr MCMV vs. Ctr non-inf.
Pkp2+/- MCMV vs. Ctr MCMV
-log10(adj. p-value)
avg_log2FC
Ifng
Il2rb
Il12rb2
Csf1r
Il2rg
Cxcr3
Csf2ra
Ifngr1
Ctla4
Ifngr2
Il6ra
Il21r
Il10rb
Cxcl9
Pdcd1
Ccr2
Cxcr4
Csf2rb
Il1b
Cd80
Met
Il2rb
Il1a
Ifng
Cxcl1
Il16
Cxcl9
Ccl5
Hgf
Regulation
Upregulated
Downregulated
Not significant
C
TNF-alpha Signaling via NF-kB
IL-6/JAK/STAT3 Signaling
Inflammatory Response
p53 Pathway
Complement
Apoptosis
Pkp2+/- non-inf.
Pkp2+/- MCMV
Ctr MCMV
vs. Ctr non-inf.
D
Ctr non-inf.
Pkp2+/- non-inf.
Ctr MCMV
Pkp2+/- MCMV
STAT3
86 kDa
GAPDH
36 kDa
STAT3
GAPDH norm. expr.
p = 0.0098
p = 0.0063

**Figure S6**

**A**, Differential abundance testing using MiloR in Ctr MCMV versus Ctr non-infected mice. **B**, Comparative cytokine expression analysis in cardiac leukocytes in Ctr MCMV versus Ctr non-infected and in *Pkp2*$^{+/-}$ MCMV versus Ctr MCMV, respectively. **C**, Differentially expressed genes in selected upregulated pathways in the subcluster Mac_TRM_MHCII+ of *Pkp2*$^{+/-}$ non-infected, *Pkp2*$^{+/-}$ MCMV and Ctr MCMV compared to Ctr non-infected.

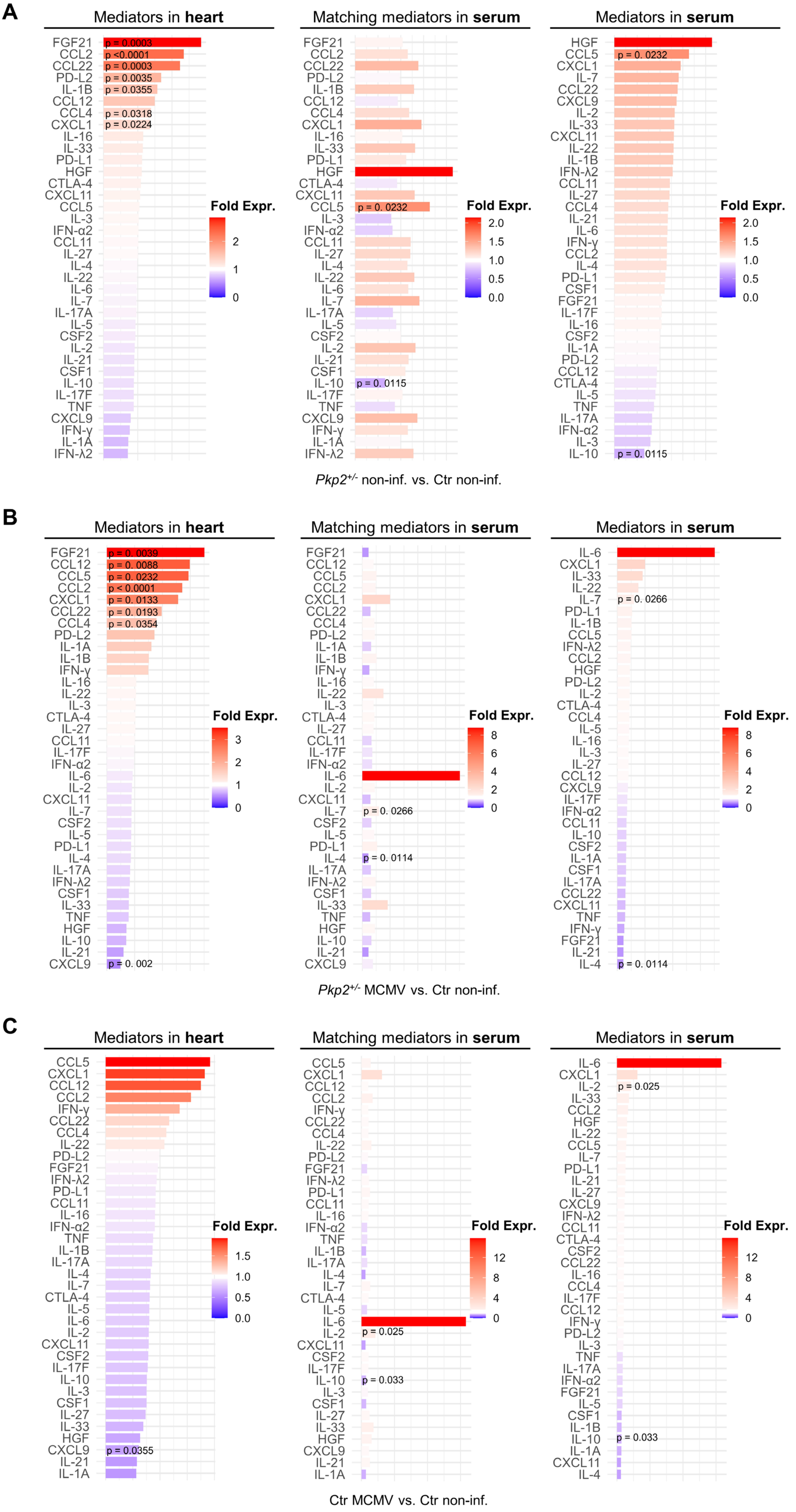
A
Mediators in heart
FGF21 CCL2 CCL22 PD-L2 IL-1B CCL12 CCL4 CXCL1 IL-16 IL-33 PD-L1 HGF CTLA-4 CXCL11 CCL5 IL-3 IFN-α2 CCL11 IL-27 IL-4 IL-22 IL-6 IL-7 IL-17A IL-5 CSF2 IL-2 IL-21 CSF1 IL-10 IL-17F TNF CXCL9 IFN-γ IL-1A IFN-λ2
p = 0.0003
p <0.0001
p = 0.0003
p = 0.0035
p = 0.0355
p = 0.0318
p = 0.0224
Fold Expr.
2
1
0
Matching mediators in serum
FGF21 CCL2 CCL22 PD-L2 IL-1B CCL12 CCL4 CXCL1 IL-16 IL-33 PD-L1 HGF CTLA-4 CXCL11 CCL5 IL-3 IFN-α2 CCL11 IL-27 IL-4 IL-22 IL-6 IL-7 IL-17A IL-5 CSF2 IL-2 IL-21 CSF1 IL-10 IL-17F TNF CXCL9 IFN-γ IL-1A IFN-λ2
p = 0. 0232
p = 0. 0115
Fold Expr.
2.0
1.5
1.0
0.5
0.0
Mediators in serum
HGF CCL5 CXCL1 IL-7 CCL22 CXCL9 IL-2 IL-33 CXCL11 IL-22 IL-1B IFN-λ2 CCL11 IL-27 CCL4 IL-21 IL-6 IFN-γ CCL2 IL-4 PD-L1 CSF1 FGF21 IL-17F IL-16 CSF2 IL-1A PD-L2 CCL12 CTLA-4 IL-5 TNF IL-17A IFN-α2 IL-3 IL-10
p = 0. 0232
p = 0. 0115
Fold Expr.
2.0
1.5
1.0
0.5
0.0
Pkp2+/- non-inf. vs. Ctr non-inf.
B
Mediators in heart
FGF21 CCL12 CCL5 CCL2 CXCL1 CCL22 CCL4 PD-L2 IL-1A IL-1B IFN-γ IL-16 IL-22 IL-3 CTLA-4 IL-27 CCL11 IL-17F IFN-α2 IL-6 IL-2 CXCL11 IL-7 CSF2 IL-5 PD-L1 IL-4 IL-17A IFN-λ2 CSF1 IL-33 TNF HGF IL-10 IL-21 CXCL9
p = 0. 0039
p = 0. 0088
p = 0. 0232
p < 0.0001
p = 0. 0133
p = 0. 0193
p = 0. 0354
p = 0. 002
Fold Expr.
3
2
1
0
Matching mediators in serum
FGF21 CCL12 CCL5 CCL2 CXCL1 CCL22 CCL4 PD-L2 IL-1A IL-1B IFN-γ IL-16 IL-22 IL-3 CTLA-4 IL-27 CCL11 IL-17F IFN-α2 IL-6 IL-2 CXCL11 IL-7 CSF2 IL-5 PD-L1 IL-4 IL-17A IFN-λ2 CSF1 IL-33 TNF HGF IL-10 IL-21 CXCL9
p = 0. 0266
p = 0. 0114
Fold Expr.
8
6
4
2
0
Mediators in serum
IL-6 CXCL1 IL-33 IL-22 IL-7 PD-L1 IL-1B CCL5 IFN-λ2 CCL2 HGF PD-L2 IL-2 CTLA-4 CCL4 IL-5 IL-16 IL-3 IL-27 CCL12 CXCL9 IL-17F IFN-α2 CCL11 IL-10 CSF2 IL-1A CSF1 IL-17A CCL22 CXCL11 TNF IFN-γ FGF21 IL-21 IL-4
p = 0. 0266
p = 0. 0114
Fold Expr.
8
6
4
2
0
Pkp2+/- MCMV vs. Ctr non-inf.
C
Mediators in heart
CCL5 CXCL1 CCL12 CCL2 IFN-γ CCL22 CCL4 IL-22 PD-L2 FGF21 IFN-λ2 PD-L1 CCL11 IL-16 IFN-α2 TNF IL-1B IL-17A IL-4 IL-7 CTLA-4 IL-5 IL-6 IL-2 CXCL11 CSF2 IL-17F IL-10 IL-3 CSF1 IL-27 IL-33 HGF CXCL9 IL-21 IL-1A
p = 0.0355
Fold Expr.
1.5
1.0
0.5
0.0
Matching mediators in serum
CCL5 CXCL1 CCL12 CCL2 IFN-γ CCL22 CCL4 IL-22 PD-L2 FGF21 IFN-λ2 PD-L1 CCL11 IL-16 IFN-α2 TNF IL-1B IL-17A IL-4 IL-7 CTLA-4 IL-5 IL-6 IL-2 CXCL11 CSF2 IL-17F IL-10 IL-3 CSF1 IL-27 IL-33 HGF CXCL9 IL-21 IL-1A
p = 0.025
p = 0.033
Fold Expr.
12
8
4
0
Mediators in serum
IL-6 CXCL1 IL-2 IL-33 CCL2 HGF IL-22 CCL5 IL-7 PD-L1 IL-21 IL-27 CXCL9 IFN-λ2 CCL11 CTLA-4 CSF2 CCL22 IL-16 CCL4 IL-17F CCL12 IFN-γ PD-L2 IL-3 TNF IL-17A IFN-α2 FGF21 IL-5 CSF1 IL-1B IL-10 IL-1A CXCL11 IL-4
p = 0.025
p = 0.033
Fold Expr.
12
8
4
0
Ctr MCMV vs. Ctr non-inf.

**Figure S7**

**A-C**, Comparative analyses of cytokine profiling data at 6mpi. The shown expression ratios compare *Pkp2*$^{+/-}$ non-infected (A), *Pkp2*$^{+/-}$ MCMV (B), and Ctr MCMV (C) to Ctr non-infected, respectively. The figures display the differentially expressed heart mediators sorted by fold expression (left) with corresponding serum levels (middle), and serum mediators sorted by fold expression (right); n = heart/serum: Ctr non-infected n = 10/10, *Pkp2*$^{+/-}$ non-infected n = 10/10, Ctr MCMV n = 10/7, *Pkp2*$^{+/-}$ MCMV n = 10/8.

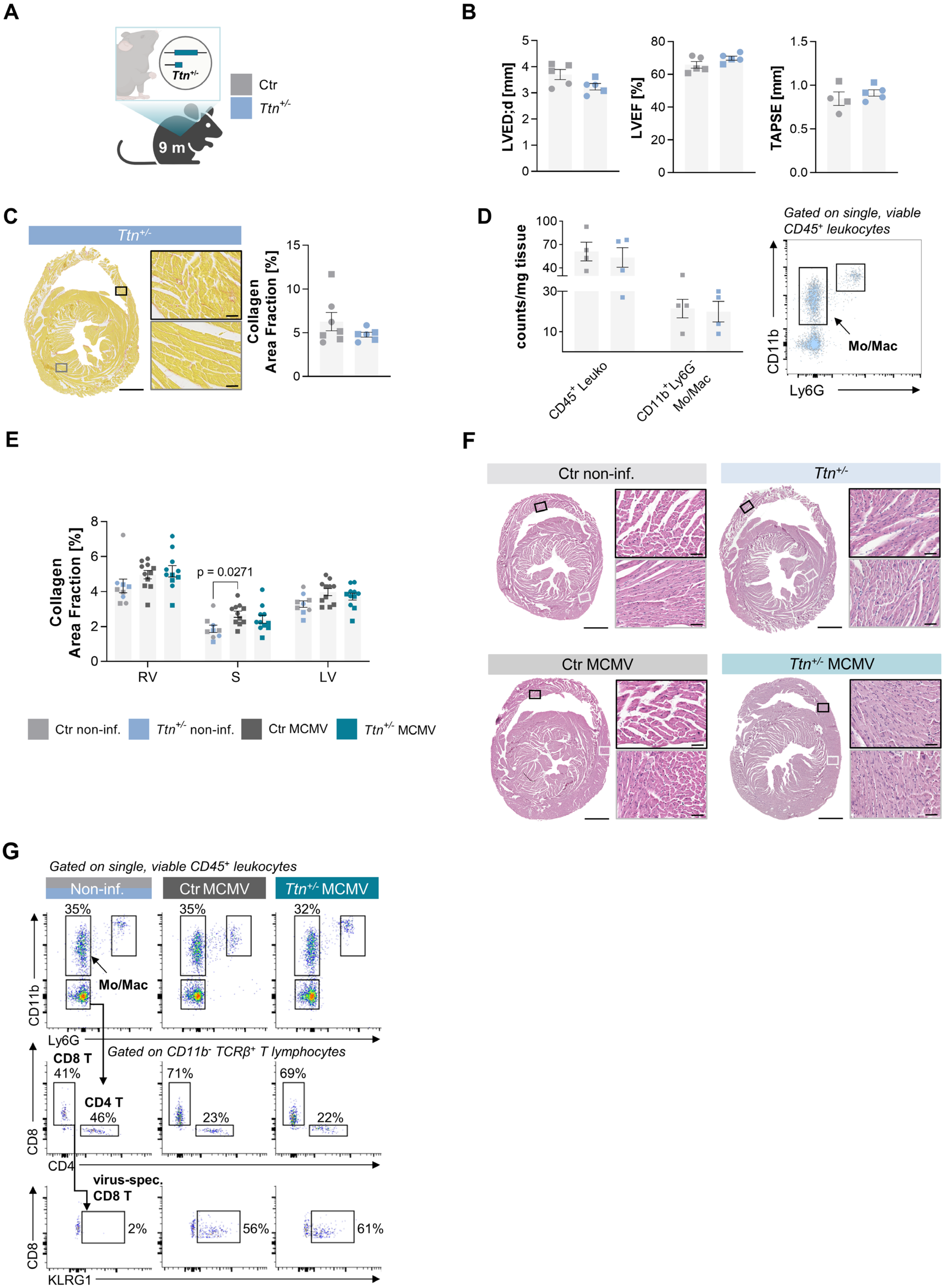
A
Ttn+/-
9 m
Ctr
Ttn+/-
B
LVED;d [mm]
LVEF [%]
TAPSE [mm]
C
Ttn+/-
Collagen Area Fraction [%]
D
counts/mg tissue
CD45+ Leuko
CD11b+Ly6G- Mo/Mac
Gated on single, viable CD45+ leukocytes
CD11b
Ly6G
Mo/Mac
E
Collagen Area Fraction [%]
p = 0.0271
RV
S
LV
Ctr non-inf.
Ttn+/- non-inf.
Ctr MCMV
Ttn+/- MCMV
F
Ctr non-inf.
Ttn+/-
Ctr MCMV
Ttn+/- MCMV
G
Gated on single, viable CD45+ leukocytes
Non-inf.
Ctr MCMV
Ttn+/- MCMV
35%
35%
32%
Mo/Mac
CD11b
Ly6G
Gated on CD11b- TCRβ+ T lymphocytes
CD8 T
41%
CD4 T
46%
71%
23%
69%
22%
CD8
CD4
virus-spec. CD8 T
2%
56%
61%
CD8
KLRG1

**Figure S8**

**A**, Scheme of heterozygous global *Ttn* mutant (*Ttn*$^{+/-}$) mouse model; signature colors: Ctr – grey, *Ttn*$^{+/-}$ – light blue. **B**, Quantification of echocardiographic parameters at 9 months; Ctr n = 4, *Ttn*$^{+/-}$ n = 5. **C**, Representative picrosirius red stained cardiac cross-section of a *Ttn*$^{+/-}$ heart at 9 months and quantification of collagen area fraction; scale bars: overview 1000µm, detail 50µm; Ctr n = 7, *Ttn*$^{+/-}$ n = 6. **D**, Quantification of cardiac CD45$^{+}$ leukocytes and monocytes/macrophages by flow cytometry in counts per milligram tissue; representative gating showing monocytes/macrophages of *Ttn*$^{+/-}$ superimposed on Ctr cells; Ctr n = 4, *Ttn*$^{+/-}$ n = 4. **E**, Quantification of collagen area fraction in picrosirius red stained cardiac sections from *Ttn*$^{+/-}$ mice challenged with MCMV at 6mpi, analyzed separately in the right ventricle, septum, and left ventricle. **F**, Representative hematoxylin-eosin staining of cardiac sections at 6mpi; scale bars: overview 1000µm, detail 50µm. **G**, Representative flow cytometry gating of 6mpi heart samples using a T-cell tailored panel. Data are shown as mean ± SEM. squares; female: circles. Unpaired t test with or without Welch's correction or Mann-Whitney test, as appropriate for 2 groups. Ordinary one-way ANOVA with or without Brown-Forsythe and Welch's correction and Tukey's or Dunnett's T3 multiple comparisons, respectively or Kruskal-Wallis test with Dunn's multiple comparisons, as appropriate for >2 groups. m – months, LVEDD – left ventricular end-diastolic diameter, LVEF – left ventricular ejection fraction, TAPSE – tricuspid annular plane systolic excursion, Leuko – leukocytes, Mo/Mac – monocytes/macrophages, RV – right ventricle, S – septum, LV – left ventricle.

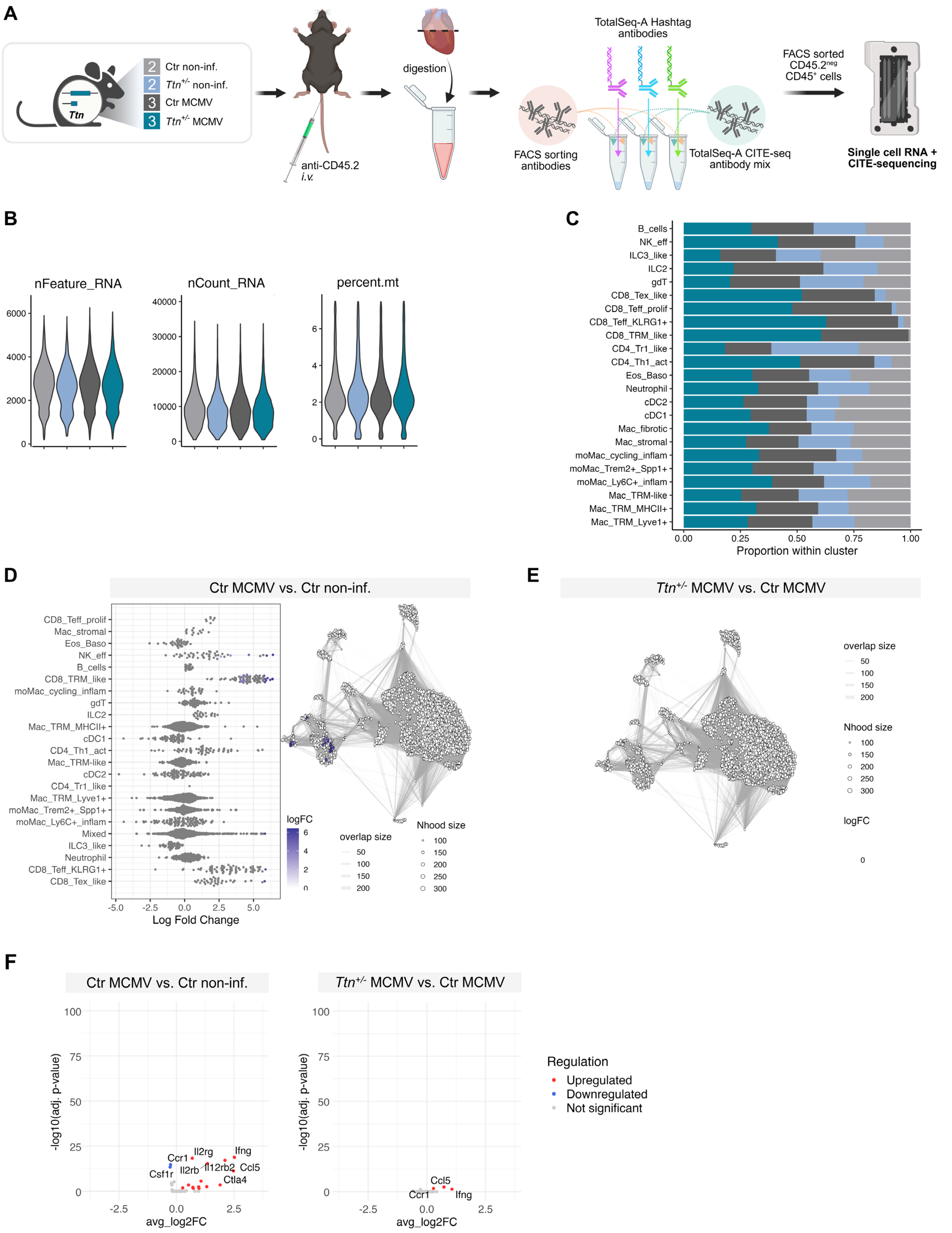
A
Ctr non-inf.
Ttn+/- non-inf.
Ctr MCMV
Ttn+/- MCMV
Ttn
anti-CD45.2
i.v.
digestion
TotalSeq-A Hashtag antibodies
FACS sorting antibodies
TotalSeq-A CITE-seq antibody mix
FACS sorted CD45.2neg CD45+ cells
Single cell RNA + CITE-sequencing
B
nFeature_RNA
nCount_RNA
percent.mt
C
Proportion within cluster
D
Ctr MCMV vs. Ctr non-inf.
Log Fold Change
logFC
overlap size
Nhood size
E
Ttn+/- MCMV vs. Ctr MCMV
F
Ctr MCMV vs. Ctr non-inf.
Ttn+/- MCMV vs. Ctr MCMV
-log10(adj. p-value)
avg_log2FC
Regulation
Upregulated
Downregulated
Not significant

**Figure S9**

**A**, Mice received an *i.v.* injection of anti-CD45.2-APC to exclude circulating leukocytes. After removing atria and valvular tissue, the remaining ventricles were processed into single-cell suspensions. Samples were multiplexed with TotalSeq-A hashtags, stained with the CITE-seq antibody mix and an anti-CD45 antibody, and sorted for $CD45^{+}$ $CD45.2\text{-}APC^{-}$ leukocytes. Two libraries comprising 10 pooled samples were sequenced: Ctr non-infected (n = 2), *Ttn*$^{+/-}$ non-infected (n = 2), Ctr MCMV (n = 3), and *Ttn*$^{+/-}$ MCMV (n = 3). **B**, Violin plots showing the distribution of nFeature_RNA, nCount_RNA, and percent.mt per cell after quality filtering. **C**, Bar plot depicting the relative contribution of each experimental group to the total number of cells within each identified cell type cluster. **D+E**, Differential abundance testing using MiloR in Ctr MCMV versus Ctr non-infected mice (D) and *Ttn*$^{+/-}$ MCMV versus Ctr MCMV mice (E). No significant changes in cell abundance were detected in the latter comparison; therefore, no beeswarm plot is shown. **F**, Comparative cytokine expression analysis in cardiac leukocytes in Ctr MCMV versus Ctr non-infected and in *Ttn*$^{+/-}$ MCMV versus Ctr MCMV, respectively. *i.v.* – intravenous, FACS – fluorescence-activated cell sorting, CITE-seq - cellular indexing of transcriptomes and epitopes by sequencing.

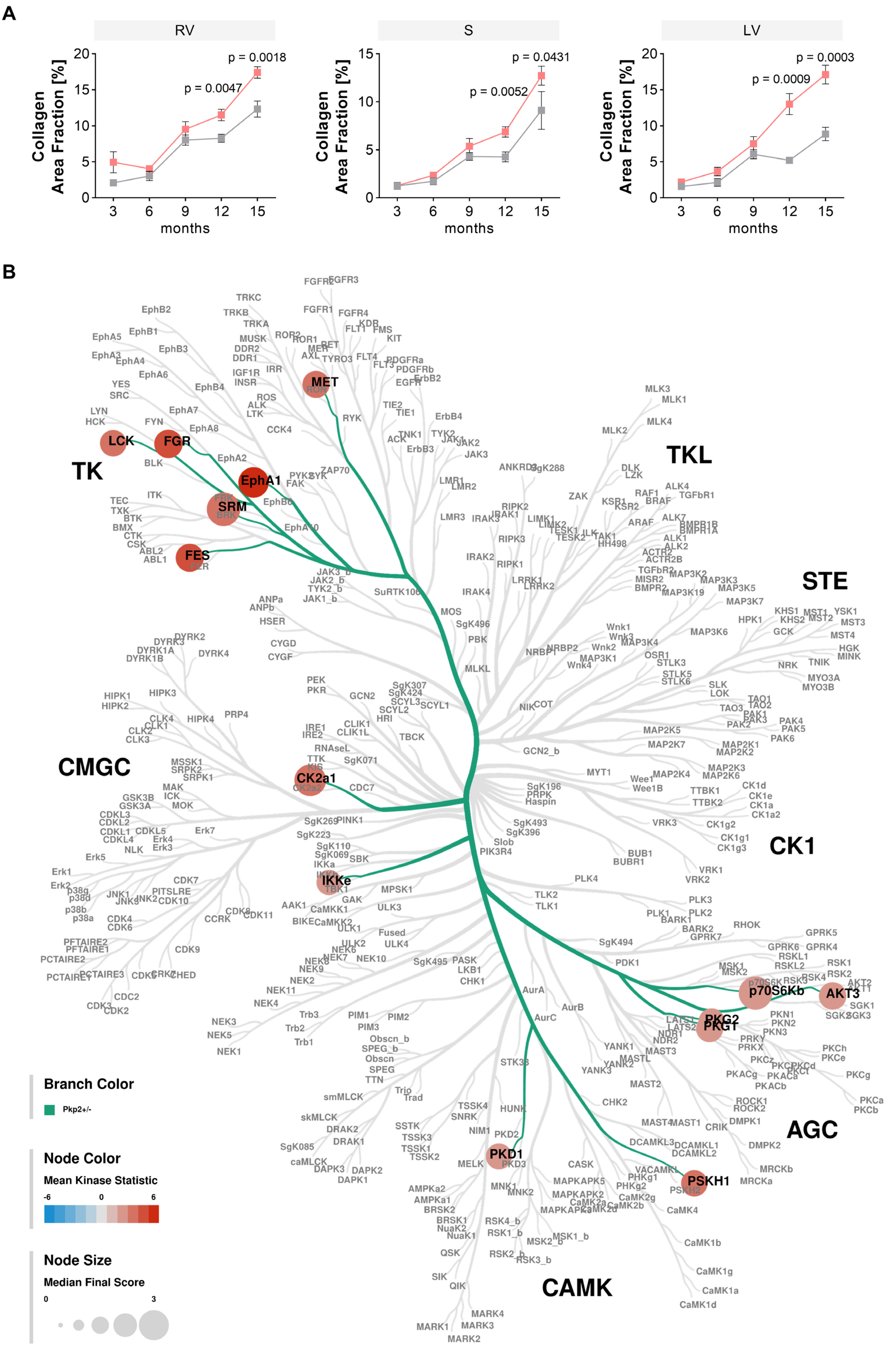
A
RV
S
LV
Collagen Area Fraction [%]
months
3 6 9 12 15
p = 0.0018
p = 0.0047
p = 0.0431
p = 0.0052
p = 0.0003
p = 0.0009
B
TK
TKL
STE
CK1
AGC
CAMK
CMGC
LCK
FGR
EphA1
SRM
FES
MET
CK2a1
IKKe
p70S6Kb
AKT3
PKG2
PKG1
PKD1
PSKH1
Branch Color
Pkp2+/-
Node Color
Mean Kinase Statistic
-6
0
6
Node Size
Median Final Score
0
3

**Figure S10**

**A**, Quantification of collagen area fraction in picrosirius red stained cardiac sections of 3 to 15-month-old *Pkp2*$^{+/-}$ and Ctr mice, analyzed separately in the right ventricle, septum, and left ventricle; n = 3mo/6mo/9mo/12mo/15mo: Ctr n = 4/7/12/6/7, *Pkp2*$^{+/-}$ n = 5/6/10/8/10. **B**, Significantly regulated kinases (specificity score ≥ 1 and significance score ≥ 0.5) from 12-month-old *Pkp2*$^{+/-}$ versus Ctr mice were mapped onto a human kinome tree using Coral for visualization. Node color represents the mean kinase statistic, indicating the direction and magnitude of regulation (red = upregulated, blue = downregulated), while node size corresponds to the median final score, serving as a surrogate for the overall confidence or strength of kinase regulation across replicates. Data are shown as mean ± SEM. Unpaired t test with or without Welch's correction or Mann-Whitney test, as appropriate for 2 groups. Ordinary one-way ANOVA with or without Brown-Forsythe and Welch's correction and Tukey's or Dunnett's T3 multiple comparisons, respectively or Kruskal-Wallis test with Dunn's multiple comparisons, as appropriate for >2 groups. P-values for comparisons between groups at a given time point are shown in the graph; p-values for longitudinal comparisons within groups are provided in Table S14-S17. RV – right ventricle, S – septum, LV – left ventricle.

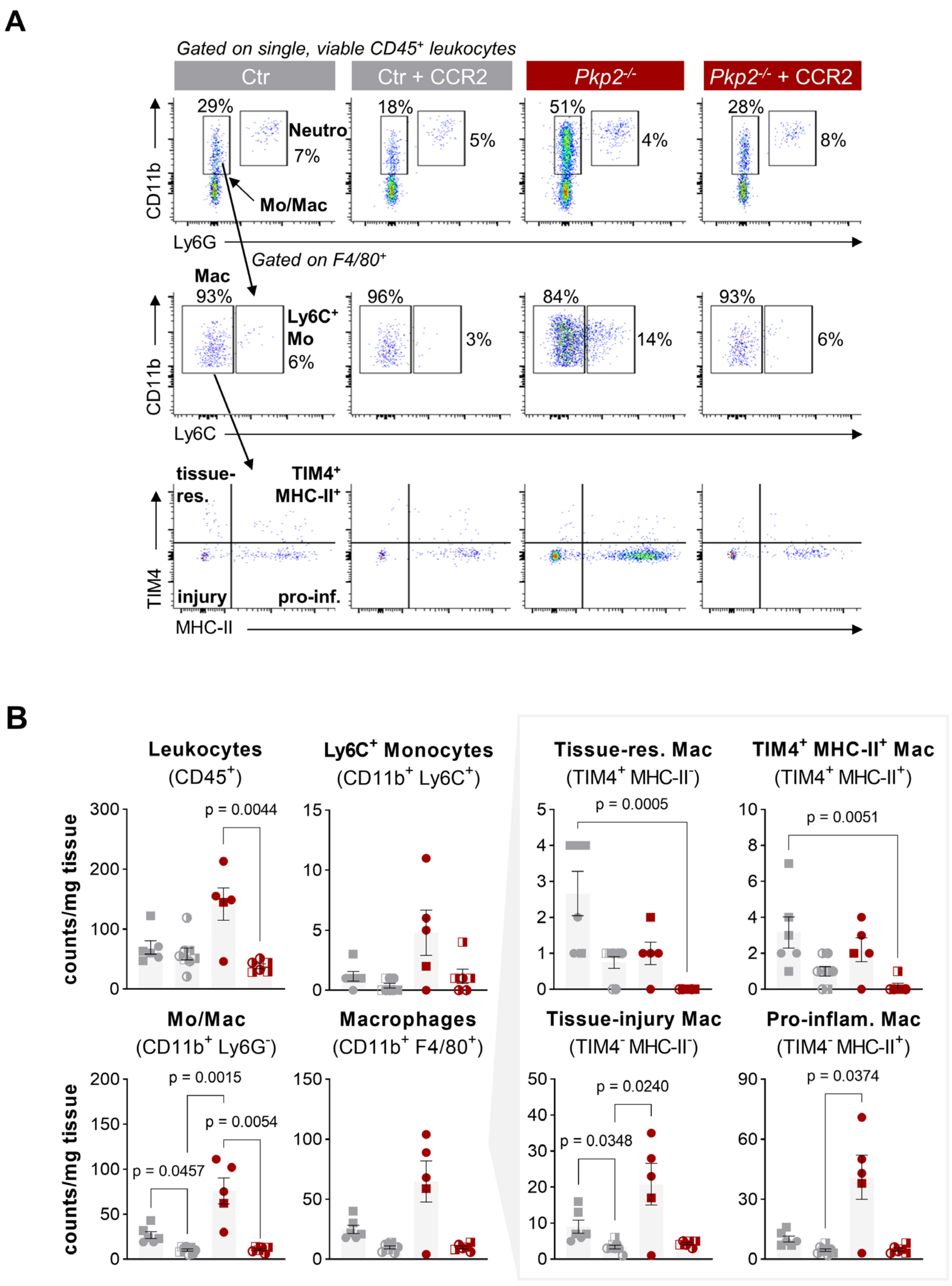

A
Gated on single, viable CD45+ leukocytes
Ctr
Ctr + CCR2
Pkp2-/-
Pkp2-/- + CCR2
CD11b
Ly6G
29%
Neutro
7%
Mo/Mac
18%
5%
51%
4%
28%
8%
Gated on F4/80+
Mac
93%
Ly6C+
Mo
6%
96%
3%
84%
14%
93%
6%
Ly6C
tissue-
res.
TIM4+
MHC-II+
injury
pro-inf.
TIM4
MHC-II
B
Leukocytes
(CD45+)
Ly6C+ Monocytes
(CD11b+ Ly6C+)
Tissue-res. Mac
(TIM4+ MHC-II-)
TIM4+ MHC-II+ Mac
(TIM4+ MHC-II+)
counts/mg tissue
p = 0.0044
p = 0.0005
p = 0.0051
Mo/Mac
(CD11b+ Ly6G-)
Macrophages
(CD11b+ F4/80+)
Tissue-injury Mac
(TIM4- MHC-II-)
Pro-inflam. Mac
(TIM4- MHC-II+)
p = 0.0015
p = 0.0054
p = 0.0457
p = 0.0240
p = 0.0348
p = 0.0374

**Figure S11**

**A+B**, More detailed representative flow cytometry gating and quantification of different cardiac CD45$^+$ leukocytes following anti-CCR2 treatment represented in counts per milligram tissue; untreated – filled symbol, anti-CCR2 treated – half-filled symbol; Ctr n = 6, Ctr+CCR2 n = 8, *Pkp2*$^{-/-}$ n = 5, *Pkp2*$^{-/-}$+CCR2 n = 6. Data are shown as mean ± SEM. Male: squares; female: circles. Ordinary one-way ANOVA with or without Brown-Forsythe and Welch's correction and Tukey's or Dunnett's T3 multiple comparisons, respectively or Kruskal-Wallis test with Dunn's multiple comparisons, as appropriate. Neutro – neutrophils, Mo/Mac – monocytes/macrophages, Mo – monocytes, Mac – macrophage

### C. Supplemental Tables

**Supplemental Table S1** Echocardiographic parameters (mean ± SEM) and p-values in 6-8-week-old *Pkp2*$^{-/-}$ and Ctr mice.

| **6-8w** | Genotype | Ctr | *Pkp2*$^{-/-}$ | Comparison | |
|---|---|---|---|---|---|
| | n | 11 | 8 | *Pkp2*$^{-/-}$ vs. Ctr | |
| | Sex [m/f] | 5/6 | 4/4 | | |
| Parameter | Unit | Value ± SEM | Value ± SEM | p-value | |
| LVAW;s | mm | 1.49 ± 0.09 | 1.18 ± 0.06 | **0.0168** | * |
| LVAW;d | mm | 0.99 ± 0.08 | 0.80 ± 0.05 | 0.0611 | n.s. |
| LVPW;s | mm | 1.39 ± 0.11 | 1.24 ± 0.14 | 0.3906 | n.s. |
| LVPW;d | mm | 0.88 ± 0.10 | 0.96 ± 0.12 | 0.6124 | n.s. |
| LVESD | mm | 1.47 ± 0.16 | 2.68 ± 0.20 | **0.0002** | *** |
| LVEDD | mm | 2.82 ± 0.26 | 3.78 ± 0.18 | **0.0068** | ** |
| HR | bpm | 510.05 ± 13.64 | 508.36 ± 13.29 | 0.9327 | n.s. |
| FS | % | 54.03 ± 3.67 | 30.49 ± 2.90 | **0.0002** | *** |
| SimpV; s | µl | 21.76 ± 2.62 | 45.65 ± 3.81 | **<0.0001** | **** |
| SimpV; d | µl | 58.34 ± 4.37 | 75.35 ± 5.16 | **0.0203** | * |
| SimpFAC | % | 52.45 ± 3.53 | 34.36 ± 2.25 | **0.001** | ** |
| SimpLVEF | % | 63.31 ± 2.57 | 39.67 ± 2.35 | **<0.0001** | **** |
| SimpCO | ml/min | 18.49 ± 1.50 | 14.92 ± 1.34 | 0.1082 | n.s. |
| SimpSV | µl | 36.58 ± 2.72 | 29.70 ± 2.44 | 0.0892 | n.s. |
| IVRT | ms | 22.85 ± 0.95 | 29.20 ± 2.28 | **0.0294** | * |
| MV A | mm/s | 417.55 ± 30.20 | 240.69 ± 39.38 | **0.0031** | ** |
| MV E | mm/s | 650.64 ± 25.10 | 590.10 ± 69.28 | 0.4329 | n.s. |
| MV E/A | a.u. | 1.62 ± 0.11 | 2.81 ± 0.52 | 0.0716 | n.s. |
| TAPSE | mm | 0.94 ± 0.09 | 0.54 ± 0.04 | **0.0019** | ** |
| RVOT;d | mm | 1.42 ± 0.08 | 1.66 ± 0.09 | 0.0694 | n.s. |

**Supplemental Table S2** Echocardiographic parameters (mean ± SEM) and p-values in 9-month-old *Pkp2*$^{+/-}$ and Ctr mice.

| **9m** | Genotype | Ctr | *Pkp2*$^{+/-}$ | Comparison | |
|---|---|---|---|---|---|
| | n | 7 | 7 | *Pkp2*$^{+/-}$ vs. Ctr | |
| | Sex [m/f] | 7/0 | 7/0 | | |
| Parameter | Unit | Value ± SEM | Value ± SEM | p-value | |
| LVAW;s | mm | 1.99 ± 0.12 | 1.83 ± 0.08 | 0.2499 | n.s. |
| LVAW;d | mm | 1.31 ± 0.08 | 1.18 ± 0.08 | 0.2134 | n.s. |
| LVPW;s | mm | 1.96 ± 0.12 | 1.75 ± 0.03 | 0.1280 | n.s. |
| LVPW;d | mm | 1.07 ± 0.06 | 1.03 ± 0.06 | 0.6419 | n.s. |
| LVESD | mm | 1.57 ± 0.17 | 1.99 ± 0.12 | 0.0601 | n.s. |
| LVEDD | mm | 3.45 ± 0.08 | 3.70 ± 0.10 | 0.0645 | n.s. |
| HR | bpm | 563.33 ± 11.39 | 577.55 ± 4.64 | 0.2811 | n.s. |
| FS | % | 60.45 ± 5.45 | 49.64 ± 2.73 | 0.1649 | n.s. |
| SimpV; s | µl | 20.48 ± 2.43 | 25.13 ± 1.78 | 0.1496 | n.s. |
| SimpV; d | µl | 75.45 ± 5.56 | 80.92 ± 2.73 | 0.2593 | n.s. |
| SimpFAC | % | 67.88 ± 2.52 | 62.85 ± 1.61 | 0.1184 | n.s. |
| SimpLVEF | % | 72.99 ± 2.03 | 68.98 ± 1.87 | 0.1716 | n.s. |
| SimpCO | ml/min | 30.54 ± 2.53 | 30.52 ± 1.79 | 0.9015 | n.s. |
| SimpSV | µl | 54.97 ± 4.01 | 55.79 ± 2.39 | 0.8630 | n.s. |
| IVRT | ms | 17.86 ± 1.29 | 19.17 ± 0.95 | 0.4299 | n.s. |
| MV A | mm/s | 356.11 ± 32.64 | 381.85 ± 29.47 | 0.5799 | n.s. |
| MV E | mm/s | 518.32 ± 45.01 | 502.51 ± 44.02 | 0.8095 | n.s. |
| MV E/A | a.u. | 1.48 ± 0.11 | 1.32 ± 0.10 | 0.3463 | n.s. |
| TAPSE | mm | 0.91 ± 0.11 | 1.00 ± 0.13 | 0.6234 | n.s. |
| RVOT;d | mm | 1.62 ± 0.05 | 1.63 ± 0.09 | 0.6200 | n.s. |

**Supplemental Table S3** Echocardiographic parameters, presented as mean ± SEM, with p-values for the most relevant comparisons between *Pkp2*$^{+/-}$ and Ctr mice. Mice were MCMV-infected/ received PBS at 3 months of age and were analyzed at 1mpi.

| **1mpi** | Group | Ctr non-inf. | *Pkp2*$^{+/-}$ non-inf. | Ctr MCMV | *Pkp2*$^{+/-}$ MCMV | Main comparisons | | | | | | | | | |
|---|---|---|---|---|---|---|---|---|---|---|---|---|---|---|---|
| | n | 6 | 7 | 6 | 8 | *Pkp2*$^{+/-}$ non-inf. vs. Ctr non-inf. | | Ctr MCMV vs. Ctr non-inf. | | *Pkp2*$^{+/-}$ MCMV vs. Ctr non-inf. | | *Pkp2*$^{+/-}$ MCMV vs. *Pkp2*$^{+/-}$ non-inf. | | *Pkp2*$^{+/-}$ MCMV vs. Ctr MCMV | |
| | Sex [m/f] | 6/0 | 7/0 | 6/0 | 8/0 | | | | | | | | | | |
| Parameter | Unit | Value ± SEM | Value ± SEM | Value ± SEM | Value ± SEM | p-value | | p-value | | p-value | | p-value | | p-value | |
| LVAW;s | mm | 1.62 ± 0.05 | 1.56 ± 0.05 | 1.73 ± 0.07 | 1.77 ± 0.06 | >0.9999 | n.s. | 0.9046 | n.s. | 0.3111 | n.s. | 0.0574 | n.s. | >0.9999 | n.s. |
| LVAW;d | mm | 0.96 ± 0.06 | 0.93 ± 0.06 | 1.05 ± 0.06 | 1.07 ± 0.05 | 0.9722 | n.s. | 0.7305 | n.s. | 0.573 | n.s. | 0.2896 | n.s. | 0.9974 | n.s. |
| LVPW;s | mm | 1.79 ± 0.09 | 1.78 ± 0.08 | 1.72 ± 0.06 | 1.78 ± 0.07 | >0.9999 | n.s. | 0.942 | n.s. | >0.9999 | n.s. | >0.9999 | n.s. | 0.9487 | n.s. |
| LVPW;d | mm | 1.09 ± 0.04 | 1.11 ± 0.05 | 1.03 ± 0.04 | 1.12 ± 0.08 | 0.9988 | n.s. | 0.8847 | n.s. | 0.9856 | n.s. | 0.9971 | n.s. | 0.6792 | n.s. |
| LVESD | mm | 1.91 ± 0.04 | 2.15 ± 0.13 | 2.03 ± 0.16 | 1.82 ± 0.12 | 0.5311 | n.s. | 0.9157 | n.s. | 0.9504 | n.s. | 0.2079 | n.s. | 0.6189 | n.s. |
| LVEDD | mm | 3.58 ± 0.13 | 3.75 ± 0.14 | 3.70 ± 0.10 | 3.49 ± 0.07 | 0.743 | n.s. | 0.9023 | n.s. | 0.9297 | n.s. | 0.3364 | n.s. | 0.551 | n.s. |
| HR | bpm | 573.47 ± 5.60 | 566.76 ± 4.57 | 563.57 ± 14.73 | 575.45 ± 5.52 | 0.9135 | n.s. | 0.9825 | n.s. | >0.9999 | n.s. | 0.7783 | n.s. | 0.9598 | n.s. |
| FS | % | 49.91 ± 1.41 | 47.77 ± 1.70 | 51.27 ± 2.91 | 50.70 ± 3.02 | 0.9314 | n.s. | 0.9825 | n.s. | 0.9956 | n.s. | 0.8139 | n.s. | 0.9984 | n.s. |
| SimpV; s | µl | 17.56 ± 1.24 | 20.14 ± 1.05 | 20.84 ± 1.61 | 21.43 ± 2.09 | 0.5452 | n.s. | 0.534 | n.s. | 0.5412 | n.s. | 0.9925 | n.s. | >0.9999 | n.s. |
| SimpV; d | µl | 64.81 ± 7.07 | 66.81 ± 3.72 | 68.51 ± 3.85 | 60.62 ± 2.68 | 0.9885 | n.s. | 0.9409 | n.s. | 0.9004 | n.s. | 0.7161 | n.s. | 0.5754 | n.s. |
| SimpFAC | % | 64.98 ± 3.82 | 63.30 ± 0.90 | 65.50 ± 2.18 | 60.46 ± 2.99 | 0.9975 | n.s. | >0.9999 | n.s. | 0.9126 | n.s. | 0.9183 | n.s. | 0.6867 | n.s. |
| SimpLVEF | % | 72.22 ± 3.95 | 69.74 ± 0.93 | 69.53 ± 1.97 | 64.69 ± 3.06 | 0.7741 | n.s. | 0.8855 | n.s. | 0.2685 | n.s. | 0.5578 | n.s. | 0.7058 | n.s. |
| SimpCO | ml/min | 27.72 ± 3.75 | 25.89 ± 1.63 | 26.32 ± 1.95 | 22.67 ± 1.20 | 0.9412 | n.s. | 0.9747 | n.s. | 0.3916 | n.s. | 0.7083 | n.s. | 0.6543 | n.s. |
| SimpSV | µl | 47.25 ± 6.37 | 46.67 ± 2.96 | 47.67 ± 3.09 | 39.19 ± 2.57 | >0.9999 | n.s. | >0.9999 | n.s. | >0.9999 | n.s. | 0.6295 | n.s. | 0.3639 | n.s. |
| IVRT | ms | 17.59 ± 1.35 | 19.60 ± 0.73 | 19.72 ± 0.95 | 18.33 ± 0.51 | 0.4108 | n.s. | 0.3933 | n.s. | 0.9366 | n.s. | 0.7296 | n.s. | 0.6990 | n.s. |
| TAPSE | mm | 1.03 ± 0.11 | 1.05 ± 0.07 | 0.91 ± 0.05 | 0.95 ± 0.04 | >0.9999 | n.s. | 0.4479 | n.s. | 0.9688 | n.s. | >0.9999 | n.s. | >0.9999 | n.s. |

**Supplemental Table S4** Echocardiographic parameters, presented as mean ± SEM, with p-values for the most relevant comparisons between *Pkp2*$^{+/-}$ and Ctr mice. Mice were MCMV-infected/ received PBS at 3 months of age and were analyzed at 3mpi.

| **3mpi** | Group | Ctr non-inf. | *Pkp2*$^{+/-}$ non-inf. | Ctr MCMV | *Pkp2*$^{+/-}$ MCMV | Main comparisons | | | | | | | | | |
|---|---|---|---|---|---|---|---|---|---|---|---|---|---|---|---|
| | n | 9 | 10 | 14 | 13 | *Pkp2*$^{+/-}$ non-inf. vs. Ctr non-inf. | | Ctr MCMV vs. Ctr non-inf. | | *Pkp2*$^{+/-}$ MCMV vs. Ctr non-inf. | | *Pkp2*$^{+/-}$ MCMV vs. *Pkp2*$^{+/-}$ non-inf. | | *Pkp2*$^{+/-}$ MCMV vs. Ctr MCMV | |
| | Sex [m/f] | 9/0 | 10/0 | 14/0 | 13/0 | | | | | | | | | | |
| Parameter | Unit | Value ± SEM | Value ± SEM | Value ± SEM | Value ± SEM | p-value | | p-value | | p-value | | p-value | | p-value | |
| LVAW;s | mm | 1.95 ± 0.10 | 1.82 ± 0.05 | 1.68 ± 0.03 | 1.71 ± 0.05 | >0.9999 | n.s. | 0.0843 | n.s. | 0.1562 | n.s. | 0.4699 | n.s. | >0.9999 | n.s. |
| LVAW;d | mm | 1.23 ± 0.08 | 1.12 ± 0.05 | 1.08 ± 0.03 | 1.09 ± 0.05 | 0.5323 | n.s. | 0.2194 | n.s. | 0.2677 | n.s. | 0.9707 | n.s. | >0.9999 | n.s. |
| LVPW;s | mm | 1.99 ± 0.08 | 1.81 ± 0.07 | 1.60 ± 0.05 | 1.69 ± 0.05 | 0.2097 | n.s. | **0.0002** | *** | **0.0082** | ** | 0.5315 | n.s. | 0.6035 | n.s. |
| LVPW;d | mm | 1.16 ± 0.07 | 1.09 ± 0.07 | 0.98 ± 0.04 | 0.99 ± 0.04 | 0.8433 | n.s. | 0.0886 | n.s. | 0.137 | n.s. | 0.5098 | n.s. | 0.9987 | n.s. |
| LVESD | mm | 1.67 ± 0.14 | 1.94 ± 0.11 | 1.96 ± 0.12 | 2.03 ± 0.10 | 0.4629 | n.s. | 0.3222 | n.s. | 0.1886 | n.s. | 0.9524 | n.s. | 0.9757 | n.s. |
| LVEDD | mm | 3.57 ± 0.09 | 3.65 ± 0.08 | 3.63 ± 0.10 | 3.80 ± 0.09 | >0.9999 | n.s. | >0.9999 | n.s. | 0.5617 | n.s. | >0.9999 | n.s. | 0.6358 | n.s. |
| HR | bpm | 571.41 ± 9.22 | 574.20 ± 5.55 | 560.14 ± 9.59 | 567.24 ± 4.93 | >0.9999 | n.s. | >0.9999 | n.s. | >0.9999 | n.s. | >0.9999 | n.s. | >0.9999 | n.s. |
| FS | % | 58.83 ± 4.35 | 52.36 ± 1.82 | 48.75 ± 1.91 | 49.96 ± 1.29 | >0.9999 | n.s. | 0.1554 | n.s. | 0.3125 | n.s. | >0.9999 | n.s. | >0.9999 | n.s. |
| SimpV; s | µl | 19.97 ± 1.32 | 23.02 ± 1.43 | 21.32 ± 1.68 | 25.15 ± 1.53 | 0.5985 | n.s. | 0.9328 | n.s. | 0.1269 | n.s. | 0.7773 | n.s. | 0.256 | n.s. |
| SimpV; d | µl | 71.55 ± 3.33 | 74.16 ± 4.09 | 71.68 ± 3.36 | 70.04 ± 2.64 | 0.9587 | n.s. | >0.9999 | n.s. | 0.99 | n.s. | 0.825 | n.s. | 0.9818 | n.s. |
| SimpFAC | % | 66.72 ± 2.18 | 62.82 ± 1.22 | 64.83 ± 1.54 | 59.10 ± 1.64 | 0.4394 | n.s. | 0.8595 | n.s. | **0.0162** | * | 0.4049 | n.s. | 0.0533 | n.s. |
| SimpEF | % | 71.90 ± 1.71 | 68.92 ± 1.02 | 70.70 ± 1.29 | 64.11 ± 1.58 | 0.5486 | n.s. | 0.9379 | n.s. | **0.0035** | ** | 0.1035 | n.s. | **0.0058** | ** |
| SimpCO | ml/min | 29.08 ± 2.04 | 27.97 ± 1.81 | 27.82 ± 1.28 | 24.03 ± 0.98 | >0.9999 | n.s. | >0.9999 | n.s. | 0.3253 | n.s. | 0.6106 | n.s. | 0.4278 | n.s. |
| SimpSV | µl | 51.58 ± 3.04 | 51.14 ± 30.4 | 50.36 ± 2.02 | 44.89 ± 2.09 | 0.9813 | n.s. | 0.9813 | n.s. | 0.3597 | n.s. | 0.3597 | n.s. | 0.3597 | n.s. |
| IVRT | ms | 17.69 ± 0.81 | 18.36 ± 0.53 | 17.38 ± 1.11 | 18.43 ± 0.84 | >0.9999 | n.s. | >0.9999 | n.s. | >0.9999 | n.s. | >0.9999 | n.s. | 0.5072 | n.s. |
| TAPSE | mm | 0.91 ± 0.08 | 1.04 ± 0.09 | 0.89 ± 0.07 | 0.99 ± 0.09 | 0.7355 | n.s. | 0.9958 | n.s. | 0.9314 | n.s. | 0.9625 | n.s. | 0.8009 | n.s. |

**Supplemental Table S5** Echocardiographic parameters, presented as mean ± SEM, with p-values for the most relevant comparisons between *Pkp2*$^{+/-}$ and Ctr mice. Mice were MCMV-infected/ received PBS at 3 months of age and were analyzed at 6mpi.

| **6mpi** | Group | Ctr non-inf. | *Pkp2*$^{+/-}$ non-inf. | Ctr MCMV | *Pkp2*$^{+/-}$ MCMV | Main comparisons | | | | | | | | | |
|---|---|---|---|---|---|---|---|---|---|---|---|---|---|---|---|
| | n | 7 | 7 | 5 | 7 | *Pkp2*$^{+/-}$ non-inf. vs. Ctr non-inf. | | Ctr MCMV vs. Ctr non-inf. | | *Pkp2*$^{+/-}$ MCMV vs. Ctr non-inf. | | *Pkp2*$^{+/-}$ MCMV vs. *Pkp2*$^{+/-}$ non-inf. | | *Pkp2*$^{+/-}$ MCMV vs. Ctr MCMV | |
| | Sex [m/f] | 7/0 | 7/0 | 5/0 | 7/0 | | | | | | | | | | |
| Parameter | Unit | Value ± SEM | Value ± SEM | Value ± SEM | Value ± SEM | p-value | | p-value | | p-value | | p-value | | p-value | |
| LVAW;s | mm | 1.99 ± 0.12 | 1.83 ± 0.08 | 1.84 ± 0.11 | 1.77 ± 0.06 | 0.5478 | n.s. | 0.6976 | n.s. | 0.3163 | n.s. | 0.9739 | n.s. | 0.9538 | n.s. |
| LVAW;d | mm | 1.31 ± 0.08 | 1.18 ± 0.08 | 1.13 ± 0.10 | 1.23 ± 0.06 | 0.5467 | n.s. | 0.3824 | n.s. | 0.8541 | n.s. | 0.9473 | n.s. | 0.8039 | n.s. |
| LVPW;s | mm | 1.96 ± 0.12 | 1.75 ± 0.03 | 1.86 ± 0.05 | 1.66 ± 0.06 | 0.4811 | n.s. | 0.9469 | n.s. | 0.2363 | n.s. | 0.7653 | n.s. | 0.1783 | n.s. |
| LVPW;d | mm | 1.07 ± 0.06 | 1.03 ± 0.06 | 1.24 ± 0.06 | 1.15 ± 0.05 | 0.9548 | n.s. | 0.2494 | n.s. | 0.7476 | n.s. | 0.4428 | n.s. | 0.7509 | n.s. |
| LVESD | mm | 1.57 ± 0.17 | 1.99 ± 0.12 | 2.02 ± 0.26 | 2.33 ± 0.14 | 0.2677 | n.s. | 0.282 | n.s. | **0.0137** | * | 0.4576 | n.s. | 0.6124 | n.s. |
| LVEDD | mm | 3.45 ± 0.08 | 3.70 ± 0.10 | 3.49 ± 0.30 | 3.70 ± 0.13 | 0.5838 | n.s. | 0.9977 | n.s. | 0.5799 | n.s. | >0.9999 | n.s. | 0.7556 | n.s. |
| HR | bpm | 563.33 ± 11.39 | 577.55 ± 4.64 | 577.28 ± 10.14 | 559.48 ± 11.58 | 0.7204 | n.s. | 0.7821 | n.s. | 0.9917 | n.s. | 0.549 | n.s. | 0.6309 | n.s. |
| FS | % | 60.45 ± 5.45 | 49.64 ± 2.73 | 48.45 ± 2.32 | 39.67 ± 2.19 | >0.9999 | n.s. | 0.793 | n.s. | **0.0022** | ** | 0.1388 | n.s. | 0.4828 | n.s. |
| SimpV; s | µl | 20.48 ± 2.43 | 25.13 ± 1.78 | 22.06 ± 4.49 | 31.56 ± 1.90 | 0.5537 | n.s. | 0.976 | n.s. | **0.0212** | * | 0.2812 | n.s. | 0.0903 | n.s. |
| SimpV; d | µl | 75.45 ± 5.56 | 80.92 ± 2.73 | 66.86 ± 10.00 | 71.59 ± 3.24 | 0.8739 | n.s. | 0.7039 | n.s. | 0.9504 | n.s. | 0.581 | n.s. | 0.9321 | n.s. |
| SimpFAC | % | 67.88 ± 2.52 | 62.85 ± 1.61 | 58.91 ± 1.06 | 45.35 ± 2.25 | 0.306 | n.s. | **0.0378** | * | **<0.0001** | **** | **<0.0001** | **** | **0.0012** | ** |
| SimpEF | % | 72.99 ± 2.03 | 68.98 ± 1.87 | 68.11 ± 1.97 | 55.77 ± 2.22 | 0.4941 | n.s. | 0.4046 | n.s. | **<0.0001** | **** | **0.0006** | *** | **0.003** | ** |
| SimpCO | ml/min | 30.54 ± 2.53 | 30.52 ± 1.79 | 25.43 ± 3.26 | 22.71 ± 1.93 | >0.9999 | n.s. | 0.4695 | n.s. | 0.0924 | n.s. | 0.0935 | n.s. | 0.8606 | n.s. |
| SimpSV | µl | 54.97 ± 4.01 | 55.79 ± 2.39 | 44.80 ± 5.59 | 40.03 ± 2.85 | >0.9999 | n.s. | >0.9999 | n.s. | 0.0786 | n.s. | **0.0279** | * | >0.9999 | n.s. |
| IVRT | ms | 17.86 ± 1.29 | 19.17 ± 0.95 | 19.06 ± 2.09 | 20.99 ± 0.81 | >0.9999 | n.s. | >0.9999 | n.s. | 0.2245 | n.s. | >0.9999 | n.s. | 0.7521 | n.s. |
| TAPSE | mm | 0.91 ± 0.11 | 1.00 ± 0.13 | 0.83 ± 0.07 | 0.84 ± 0.07 | 0.9275 | n.s. | 0.9447 | n.s. | 0.9538 | n.s. | 0.6759 | n.s. | 0.9998 | n.s. |

**Supplemental Table S6** Group sizes and within-group p-values for selected echocardiographic parameters over time post-infection in the *Pkp2*$^{+/-}$ cohort.

| | Group | Ctr non-inf. | | *Pkp2*$^{+/-}$ non-inf. | | Ctr MCMV | | *Pkp2*$^{+/-}$ MCMV | |
|---|---|---|---|---|---|---|---|---|---|
| 1mpi | n | 6 | | 7 | | 6 | | 8 | |
| | Sex [m/f] | 6/0 | | 7/0 | | 6/0 | | 8/0 | |
| 3mpi | n | 9 | | 10 | | 14 | | 13 | |
| | Sex [m/f] | 9/0 | | 10/0 | | 14/0 | | 13/0 | |
| 6mpi | n | 7 | | 7 | | 5 | | 7 | |
| | Sex [m/f] | 7/0 | | 7/0 | | 5/0 | | 7/0 | |
| LVEDD | 1mpi vs. 3mpi | 0.9947 | n.s. | 0.8004 | n.s. | >0.9999 | n.s. | 0.0645 | n.s. |
| | 1mpi vs. 6mpi | 0.6345 | n.s. | 0.9602 | n.s. | >0.9999 | n.s. | 0.3270 | n.s. |
| | 3mpi vs. 6mpi | 0.6385 | n.s. | 0.9369 | n.s. | >0.9999 | n.s. | 0.7719 | n.s. |
| HR | 1mpi vs. 3mpi | >0.9999 | n.s. | 0.5171 | n.s. | >0.9999 | n.s. | 0.6714 | n.s. |
| | 1mpi vs. 6mpi | >0.9999 | n.s. | 0.5018 | n.s. | >0.9999 | n.s. | 0.3215 | n.s. |
| | 3mpi vs. 6mpi | >0.9999 | n.s. | >0.9999 | n.s. | 0.7959 | n.s. | 0.7192 | n.s. |
| FS | 1mpi vs. 3mpi | 0.3334 | n.s. | 0.2774 | n.s. | 0.7325 | n.s. | 0.9638 | n.s. |
| | 1mpi vs. 6mpi | 0.1996 | n.s. | 0.8263 | n.s. | 0.7764 | n.s. | **0.0061** | ** |
| | 3mpi vs. 6mpi | >0.9999 | n.s. | 0.6244 | n.s. | 0.9962 | n.s. | **0.0054** | ** |
| SimpFAC | 1mpi vs. 3mpi | 0.8976 | n.s. | 0.9602 | n.s. | 0.9623 | n.s. | 0.8950 | n.s. |
| | 1mpi vs. 6mpi | 0.7659 | n.s. | 0.9708 | n.s. | 0.1152 | n.s. | **0.0006** | *** |
| | 3mpi vs. 6mpi | 0.9488 | n.s. | 0.9998 | n.s. | 0.0969 | n.s. | **0.0006** | *** |
| SimpLVEF | 1mpi vs. 3mpi | >0.9999 | n.s. | 0.8932 | n.s. | 0.8703 | n.s. | 0.9796 | n.s. |
| | 1mpi vs. 6mpi | >0.9999 | n.s. | 0.9208 | n.s. | 0.8748 | n.s. | **0.0418** | * |
| | 3mpi vs. 6mpi | >0.9999 | n.s. | 0.9994 | n.s. | 0.5565 | n.s. | **0.0349** | * |
| SimpCO | 1mpi vs. 3mpi | >0.9999 | n.s. | 0.6898 | n.s. | 0.8329 | n.s. | 0.7223 | n.s. |
| | 1mpi vs. 6mpi | 0.9763 | n.s. | 0.2275 | n.s. | 0.9595 | n.s. | 0.9998 | n.s. |
| | 3mpi vs. 6mpi | >0.9999 | n.s. | 0.5737 | n.s. | 0.6702 | n.s. | 0.7543 | n.s. |
| SimpSV | 1mpi vs. 3mpi | 0.7453 | n.s. | 0.527 | n.s. | 0.802 | n.s. | 0.1398 | n.s. |
| | 1mpi vs. 6mpi | 0.3009 | n.s. | 0.1224 | n.s. | 0.8498 | n.s. | >0.9999 | n.s. |
| | 3mpi vs. 6mpi | >0.9999 | n.s. | 0.5009 | n.s. | 0.4487 | n.s. | 0.2732 | n.s. |
| IVRT | 1mpi vs. 3mpi | >0.9999 | n.s. | 0.4361 | n.s. | 0.0962 | n.s. | 0.9959 | n.s. |
| | 1mpi vs. 6mpi | >0.9999 | n.s. | 0.9135 | n.s. | >0.9999 | n.s. | 0.1166 | n.s. |
| | 3mpi vs. 6mpi | >0.9999 | n.s. | 0.6993 | n.s. | 0.9609 | n.s. | 0.0839 | n.s. |
| TAPSE | 1mpi vs. 3mpi | 0.7066 | n.s. | >0.9999 | n.s. | 0.9762 | n.s. | 0.9791 | n.s. |
| | 1mpi vs. 6mpi | 0.7339 | n.s. | >0.9999 | n.s. | 0.8120 | n.s. | 0.4424 | n.s. |
| | 3mpi vs. 6mpi | >0.9999 | n.s. | >0.9999 | n.s. | 0.8649 | n.s. | 0.4793 | n.s. |

**Supplemental Table S7** P-values for group-wise and within-group comparisons of $CD8^+$ T-cell density at defined time points post MCMV infection, with sample sizes per group.

*Group-wise comparisons at a certain time point*

| **$CD8^+$ T-cell density [cells/mm$^2$]** | Acute | | 1mpi | | 6mpi | |
|---|---|---|---|---|---|---|
| Ctr non-inf. vs. *Pkp2$^{+/-}$* non-inf. | >0.9999 | n.s. | >0.9999 | n.s. | >0.9999 | n.s. |
| Ctr non-inf. vs. Ctr MCMV | **0.0047** | ** | 0.5724 | n.s. | 0.0667 | n.s. |
| Ctr non-inf. vs. *Pkp2$^{+/-}$* MCMV | **0.0006** | *** | **0.0375** | * | **0.002** | ** |
| *Pkp2$^{+/-}$* non-inf. vs. Ctr MCMV | **0.0416** | * | 0.9108 | n.s. | 0.4342 | n.s. |
| *Pkp2$^{+/-}$* non-inf. vs. *Pkp2$^{+/-}$* MCMV | **0.0085** | ** | **0.048** | * | **0.0299** | * |
| Ctr MCMV vs. *Pkp2$^{+/-}$* MCMV | >0.9999 | n.s. | >0.9999 | n.s. | >0.9999 | n.s. |

*Time-wise comparisons within a certain group*

| **$CD8^+$ T-cell density [cells/mm$^2$]** | Ctr non-inf. | | *Pkp2$^{+/-}$* non-inf. | | Ctr MCMV | | *Pkp2$^{+/-}$* MCMV | |
|---|---|---|---|---|---|---|---|---|
| Acute vs. 1mpi | **0.0382** | * | 0.31 | n.s. | **0.027** | * | **0.0078** | ** |
| Acute vs. 6mpi | 0.398 | n.s. | >0.9999 | n.s. | **0.0017** | ** | **0.0008** | *** |
| 1mpi vs. 6mpi | 0.6517 | n.s. | **0.0447** | * | >0.9999 | n.s. | 0.9466 | n.s. |

*Sample size used at each time point*

| Group | | Ctr non-inf. | *Pkp2$^{+/-}$* non-inf. | Ctr MCMV | *Pkp2$^{+/-}$* MCMV |
|---|---|---|---|---|---|
| Acute | n | 11 | 4 | 9 | 7 |
| | Sex [m/f] | 6/5 | 0/4 | 9/0 | 7/0 |
| 1mpi | n | 4 | 6 | 5 | 11 |
| | Sex [m/f] | 4/0 | 6/0 | 5/0 | 11/0 |
| 6mpi | n | 8 | 9 | 5 | 7 |
| | Sex [m/f] | 8/0 | 9/0 | 5/0 | 7/0 |

**Supplemental Table S8** Cytokine concentrations (mean ± SEM) in heart tissue at 6mpi.

| Group | Ctr non-inf. | *Pkp2*$^{+/-}$ non-inf. | Ctr MCMV | *Pkp2*$^{+/-}$ MCMV | Main comparisons | | | | | | | | | |
|---|---|---|---|---|---|---|---|---|---|---|---|---|---|---|
| n | 10 | 10 | 10 | 10 | *Pkp2*$^{+/-}$ non-inf. vs. Ctr non-inf. | | Ctr MCMV vs. Ctr non-inf. | | *Pkp2*$^{+/-}$ MCMV vs. Ctr non-inf. | | *Pkp2*$^{+/-}$ MCMV vs. *Pkp2*$^{+/-}$ non-inf. | | *Pkp2*$^{+/-}$ MCMV vs. Ctr MCMV | |
| Sex [m/f] | 10/0 | 10/0 | 10/0 | 10/0 | | | | | | | | | | |
| Mediator | Values ± SEM [pg/ml] | | | | p-values | | | | | | | | | |
| IL-1α | 1.17 ± 0.38 | 0.84 ± 0.15 | 0.66 ± 0.08 | 1.87 ± 1.01 | >0.9999 | n.s. | >0.9999 | n.s. | >0.9999 | n.s. | >0.9999 | n.s. | >0.9999 | n.s. |
| IL-1β | 0.13 ± 0.01 | 0.20 ± 0.02 | 0.11 ± 0.02 | 0.19 ± 0.04 | 0.1130 | n.s. | 0.9720 | n.s. | 0.4852 | n.s. | >0.9999 | n.s. | 0.3108 | n.s. |
| IL-6 | 12.97 ± 2.50 | 12.55 ± 2.10 | 10.40 ± 1.93 | 12.02 ± 2.41 | >0.9999 | n.s. | >0.9999 | n.s. | >0.9999 | n.s. | >0.9999 | n.s. | >0.9999 | n.s. |
| IL-17A | 0.04 ± 0.01 | 0.04 ± 0.00 | 0.03 ± 0.00 | 0.03 ± 0.01 | >0.9999 | n.s. | >0.9999 | n.s. | >0.9999 | n.s. | >0.9999 | n.s. | >0.9999 | n.s. |
| IL-17F | 0.14 ± 0.04 | 0.13 ± 0.03 | 0.11 ± 0.03 | 0.14 ± 0.04 | 0.9800 | n.s. | 0.8879 | n.s. | 0.9999 | n.s. | 0.9926 | n.s. | 0.9379 | n.s. |
| IL-22 | 0.16 ± 0.00 | 0.15 ± 0.01 | 0.17 ± 0.02 | 0.17 ± 0.02 | 0.9998 | n.s. | 0.9815 | n.s. | 0.9990 | n.s. | 0.9962 | n.s. | >0.9999 | n.s. |
| TNFα | 4.23 ± 0.50 | 3.73 ± 0.38 | 3.69 ± 0.36 | 3.32 ± 0.31 | >0.9999 | n.s. | >0.9999 | n.s. | >0.9999 | n.s. | >0.9999 | n.s. | >0.9999 | n.s. |
| IFN-γ | 0.03 ± 0.00 | 0.02 ± 0.00 | 0.04 ± 0.01 | 0.05 ± 0.01 | >0.9999 | n.s. | >0.9999 | n.s. | >0.9999 | n.s. | 0.6743 | n.s. | >0.9999 | n.s. |
| IFN-λ2 | 0.03 ± 0.01 | 0.02 ± 0.01 | 0.03 ± 0.01 | 0.02 ± 0.00 | >0.9999 | n.s. | >0.9999 | n.s. | >0.9999 | n.s. | >0.9999 | n.s. | >0.9999 | n.s. |
| IFN-α2 | 0.03 ± 0.00 | 0.03 ± 0.00 | 0.03 ± 0.00 | 0.03 ± 0.00 | >0.9999 | n.s. | >0.9999 | n.s. | >0.9999 | n.s. | >0.9999 | n.s. | >0.9999 | n.s. |
| IL-4 | 0.17 ± 0.01 | 0.16 ± 0.01 | 0.14 ± 0.03 | 0.14 ± 0.02 | >0.9999 | n.s. | 0.8448 | n.s. | >0.9999 | n.s. | >0.9999 | n.s. | >0.9999 | n.s. |
| IL-10 | 0.05 ± 0.01 | 0.05 ± 0.02 | 0.04 ± 0.01 | 0.04 ± 0.01 | >0.9999 | n.s. | >0.9999 | n.s. | >0.9999 | n.s. | >0.9999 | n.s. | >0.9999 | n.s. |
| IL-21 | 3.61 ± 0.76 | 3.24 ± 0.68 | 2.08 ± 0.64 | 2.17 ± 0.73 | 0.9818 | n.s. | 0.4748 | n.s. | 0.4979 | n.s. | 0.6774 | n.s. | 0.9998 | n.s. |
| IL-27 | 2.16 ± 0.48 | 2.17 ± 0.41 | 1.59 ± 0.33 | 2.22 ± 0.55 | >0.9999 | n.s. | >0.9999 | n.s. | >0.9999 | n.s. | >0.9999 | n.s. | >0.9999 | n.s. |
| CXCL1 | 0.46 ± 0.05 | 0.62 ± 0.04 | 0.83 ± 0.18 | 1.17 ± 0.40 | 0.3747 | n.s. | 0.3428 | n.s. | 0.1040 | n.s. | >0.9999 | n.s. | >0.9999 | n.s. |
| CXCL9 | 39.07 ± 4.22 | 30.99 ± 3.39 | 23.90 ± 4.64 | 19.09 ± 3.56 | >0.9999 | n.s. | 0.1239 | n.s. | **0.0151** | * | 0.2676 | n.s. | >0.9999 | n.s. |
| CXCL11 | 0.58 ± 0.13 | 0.62 ± 0.13 | 0.46 ± 0.09 | 0.51 ± 0.14 | >0.9999 | n.s. | >0.9999 | n.s. | >0.9999 | n.s. | >0.9999 | n.s. | >0.9999 | n.s. |
| IL-16 | 1421.00 ± 91.08 | 1646.00 ± 106.30 | 1288.00 ± 110.20 | 1506.00 ± 49.04 | 0.3314 | n.s. | 0.7468 | n.s. | 0.9223 | n.s. | 0.7267 | n.s. | 0.3876 | n.s. |
| CCL11 | 7.20 ± 0.98 | 7.31 ± 0.68 | 6.59 ± 0.34 | 7.24 ± 0.95 | >0.9999 | n.s. | 0.9896 | n.s. | >0.9999 | n.s. | >0.9999 | n.s. | 0.9830 | n.s. |
| CCL2 | 34.48 ± 1.98 | 80.80 ± 7.38 | 54.07 ± 9.09 | 93.58 ± 10.41 | **0.0018** | ** | 0.7278 | n.s. | **0.0004** | *** | >0.9999 | n.s. | 0.0726 | n.s. |
| CCL5 | 9.57 ± 0.83 | 10.03 ± 0.92 | 18.38 ± 3.33 | 28.11 ± 6.46 | >0.9999 | n.s. | 0.4152 | n.s. | 0.0733 | n.s. | 0.1008 | n.s. | >0.9999 | n.s. |
| CCL22 | 2.71 ± 0.27 | 6.05 ± 0.78 | 3.17 ± 0.65 | 5.34 ± 0.92 | **0.0102** | * | 0.9834 | n.s. | 0.0957 | n.s. | 0.9905 | n.s. | 0.3287 | n.s. |
| CCL4 | 0.85 ± 0.08 | 1.18 ± 0.12 | 0.95 ± 0.14 | 1.52 ± 0.26 | 0.1709 | n.s. | 0.9879 | n.s. | 0.1672 | n.s. | 0.7897 | n.s. | 0.3422 | n.s. |
| CCL12 | 11.49 ± 1.66 | 17.21 ± 3.35 | 20.13 ± 5.84 | 34.28 ± 6.84 | 0.5770 | n.s. | 0.6495 | n.s. | **0.0469** | * | 0.2108 | n.s. | 0.5416 | n.s. |
| CSF1 | 50.89 ± 7.21 | 45.59 ± 4.60 | 38.30 ± 6.17 | 40.25 ± 6.50 | 0.9297 | n.s. | 0.4852 | n.s. | 0.6217 | n.s. | 0.9285 | n.s. | 0.9961 | n.s. |
| CSF2 | 0.21 ± 0.03 | 0.19 ± 0.02 | 0.16 ± 0.02 | 0.18 ± 0.03 | >0.9999 | n.s. | >0.9999 | n.s. | >0.9999 | n.s. | >0.9999 | n.s. | >0.9999 | n.s. |
| HGF | 27.71 ± 4.83 | 30.98 ± 4.21 | 17.58 ± 4.15 | 19.50 ± 3.18 | 0.9433 | n.s. | 0.3228 | n.s. | 0.5286 | n.s. | 0.2423 | n.s. | 0.9884 | n.s. |
| FGF21 | 0.63 ± 0.07 | 1.79 ± 0.36 | 0.60 ± 0.17 | 2.21 ± 0.56 | 0.0333 | * | >0.9999 | n.s. | 0.0527 | n.s. | >0.9999 | n.s. | **0.0141** | * |
| IL-3 | 0.04 ± 0.01 | 0.04 ± 0.01 | 0.03 ± 0.01 | 0.04 ± 0.01 | >0.9999 | n.s. | >0.9999 | n.s. | >0.9999 | n.s. | >0.9999 | n.s. | >0.9999 | n.s. |
| PD-L2 | 3.04 ± 0.36 | 5.11 ± 0.50 | 3.00 ± 0.39 | 5.20 ± 1.04 | 0.0589 | n.s. | >0.9999 | n.s. | 0.2557 | n.s. | >0.9999 | n.s. | 0.1118 | n.s. |
| PD-L1 | 343.80 ± 21.85 | 387.20 ± 67.83 | 316.40 ± 35.21 | 299.30 ± 39.52 | >0.9999 | n.s. | >0.9999 | n.s. | >0.9999 | n.s. | >0.9999 | n.s. | >0.9999 | n.s. |
| CTLA4 | 0.13 ± 0.03 | 0.14 ± 0.02 | 0.10 ± 0.02 | 0.13 ± 0.03 | >0.9999 | n.s. | >0.9999 | n.s. | >0.9999 | n.s. | >0.9999 | n.s. | >0.9999 | n.s. |
| IL-7 | 4.35 ± 0.78 | 4.19 ± 0.65 | 3.55 ± 0.58 | 3.86 ± 0.80 | >0.9999 | n.s. | >0.9999 | n.s. | >0.9999 | n.s. | >0.9999 | n.s. | >0.9999 | n.s. |
| IL-2 | 0.49 ± 0.06 | 0.45 ± 0.05 | 0.39 ± 0.05 | 0.44 ± 0.07 | >0.9999 | n.s. | >0.9999 | n.s. | >0.9999 | n.s. | >0.9999 | n.s. | >0.9999 | n.s. |
| IL-5 | 0.06 ± 0.01 | 0.05 ± 0.01 | 0.05 ± 0.01 | 0.05 ± 0.01 | >0.9999 | n.s. | >0.9999 | n.s. | >0.9999 | n.s. | >0.9999 | n.s. | >0.9999 | n.s. |
| IL-33 | 21.10 ± 2.75 | 24.20 ± 2.67 | 14.61 ± 1.34 | 16.67 ± 2.53 | 0.7966 | n.s. | 0.2373 | n.s. | 0.5626 | n.s. | 0.1355 | n.s. | 0.9279 | n.s. |

**Supplemental Table S9** Cytokine concentrations (mean ± SEM) in serum at 6mpi.

| Group | Ctr non-inf. | *Pkp2*$^{+/-}$ non-inf. | Ctr MCMV | *Pkp2*$^{+/-}$ MCMV | Main comparisons | | | | | | | | | |
|---|---|---|---|---|---|---|---|---|---|---|---|---|---|---|
| n | 10 | 10 | 7 | 8 | *Pkp2*$^{+/-}$ non-inf. vs. Ctr non-inf. | | Ctr MCMV vs. Ctr non-inf. | | *Pkp2*$^{+/-}$ MCMV vs. Ctr non-inf. | | *Pkp2*$^{+/-}$ MCMV vs. *Pkp2*$^{+/-}$ non-inf. | | *Pkp2*$^{+/-}$ MCMV vs. Ctr MCMV | |
| Sex [m/f] | 10/0 | 10/0 | 7/0 | 8/0 | | | | | | | | | | |
| Mediator | Values ± SEM [pg/ml] | | | | p-values | | | | | | | | | |
| IL-1α | 52.20 ± 27.21 | 51.42 ± 17.43 | 35.02 ± 11.50 | 41.29 ± 11.78 | >0.9999 | n.s. | >0.9999 | n.s. | >0.9999 | n.s. | >0.9999 | n.s. | >0.9999 | n.s. |
| IL-1β | 0.88 ± 0.29 | 1.13 ± 0.31 | 0.62 ± 0.16 | 1.12 ± 0.52 | >0.9999 | n.s. | >0.9999 | n.s. | >0.9999 | n.s. | >0.9999 | n.s. | >0.9999 | n.s. |
| IL-6 | 4.61 ± 0.87 | 5.36 ± 1.59 | 72.88 ± 59.73 | 40.49 ± 30.28 | >0.9999 | n.s. | >0.9999 | n.s. | >0.9999 | n.s. | >0.9999 | n.s. | >0.9999 | n.s. |
| IL-17A | 2.12 ± 0.66 | 1.75 ± 0.20 | 1.85 ± 0.20 | 1.64 ± 0.28 | >0.9999 | n.s. | >0.9999 | n.s. | >0.9999 | n.s. | >0.9999 | n.s. | >0.9999 | n.s. |
| IL-17F | 6.02 ± 0.97 | 6.25 ± 1.44 | 6.28 ± 1.03 | 5.43 ± 1.13 | >0.9999 | n.s. | >0.9999 | n.s. | >0.9999 | n.s. | >0.9999 | n.s. | >0.9999 | n.s. |
| IL-22 | 8.10 ± 1.51 | 10.50 ± 2.09 | 12.20 ± 1.89 | 15.41 ± 6.16 | >0.9999 | n.s. | 0.8945 | n.s. | >0.9999 | n.s. | >0.9999 | n.s. | >0.9999 | n.s. |
| TNFα | 6.44 ± 0.47 | 5.59 ± 1.17 | 5.64 ± 0.94 | 4.55 ± 0.83 | 0.9309 | n.s. | 0.9309 | n.s. | 0.6016 | n.s. | 0.9309 | n.s. | 0.9309 | n.s. |
| IFN-γ | 0.57 ± 0.08 | 0.66 ± 0.10 | 0.59 ± 0.06 | 0.36 ± 0.08 | >0.9999 | n.s. | >0.9999 | n.s. | 0.8311 | n.s. | 0.3038 | n.s. | 0.619 | n.s. |
| IFN-λ2 | 0.26 ± 0.02 | 0.33 ± 0.07 | 0.29 ± 0.03 | 0.31 ± 0.06 | >0.9999 | n.s. | >0.9999 | n.s. | >0.9999 | n.s. | >0.9999 | n.s. | >0.9999 | n.s. |
| IFN-α2 | 0.08 ± 0.01 | 0.07 ± 0.01 | 0.07 ± 0.01 | 0.07 ± 0.01 | >0.9999 | n.s. | >0.9999 | n.s. | >0.9999 | n.s. | >0.9999 | n.s. | >0.9999 | n.s. |
| IL-4 | 0.17 ± 0.03 | 0.20 ± 0.04 | 0.10 ± 0.02 | 0.09 ± 0.01 | >0.9999 | n.s. | 0.9785 | n.s. | 0.114 | n.s. | 0.0911 | n.s. | >0.9999 | n.s. |
| IL-10 | 4.80 ± 0.61 | 3.15 ± 0.45 | 3.32 ± 0.36 | 3.92 ± 0.79 | 0.0641 | n.s. | 0.4622 | n.s. | >0.9999 | n.s. | >0.9999 | n.s. | >0.9999 | n.s. |
| IL-21 | 0.93 ± 0.26 | 1.08 ± 0.22 | 1.18 ± 0.42 | 0.51 ± 0.15 | 0.9682 | n.s. | 0.9112 | n.s. | 0.6858 | n.s. | 0.4308 | n.s. | 0.3785 | n.s. |
| IL-27 | 2.74 ± 0.29 | 3.31 ± 1.23 | 3.35 ± 0.61 | 2.77 ± 0.37 | >0.9999 | n.s. | >0.9999 | n.s. | >0.9999 | n.s. | >0.9999 | n.s. | >0.9999 | n.s. |
| CXCL1 | 18.73 ± 1.57 | 27.19 ± 4.97 | 58.08 ± 28.55 | 46.88 ± 18.35 | 0.9752 | n.s. | 0.6944 | n.s. | >0.9999 | n.s. | >0.9999 | n.s. | >0.9999 | n.s. |
| CXCL9 | 163.70 ± 17.55 | 223.00 ± 71.74 | 187.10 ± 30.16 | 155.60 ± 15.35 | >0.9999 | n.s. | >0.9999 | n.s. | >0.9999 | n.s. | >0.9999 | n.s. | >0.9999 | n.s. |
| CXCL11 | 0.05 ± 0.01 | 0.07 ± 0.01 | 0.03 ± 0.01 | 0.04 ± 0.01 | 0.6335 | n.s. | 0.5009 | n.s. | 0.7464 | n.s. | 0.1644 | n.s. | 0.9742 | n.s. |
| IL-16 | 2596.00 ± 326.40 | 2681.00 ± 238.20 | 2733.00 ± 474.10 | 2698.00 ± 474.60 | >0.9999 | n.s. | >0.9999 | n.s. | >0.9999 | n.s. | >0.9999 | n.s. | >0.9999 | n.s. |
| CCL11 | 269.40 ± 23.98 | 327.50 ± 34.69 | 291.20 ± 25.39 | 222.30 ± 23.57 | 0.4225 | n.s. | 0.9519 | n.s. | 0.6404 | n.s. | 0.0584 | n.s. | 0.3999 | n.s. |
| CCL2 | 399.20 ± 42.69 | 459.20 ± 66.91 | 671.40 ± 217.50 | 475.70 ± 103.40 | >0.9999 | n.s. | >0.9999 | n.s. | >0.9999 | n.s. | >0.9999 | n.s. | >0.9999 | n.s. |
| CCL5 | 48.27 ± 6.35 | 78.89 ± 11.12 | 67.65 ± 14.04 | 61.56 ± 17.02 | 0.229 | n.s. | >0.9999 | n.s. | >0.9999 | n.s. | 0.6789 | n.s. | >0.9999 | n.s. |
| CCL22 | 342.20 ± 37.83 | 473.60 ± 51.68 | 365.30 ± 43.41 | 250.00 ± 30.40 | 0.1167 | n.s. | 0.9826 | n.s. | 0.4324 | n.s. | **0.0043** | ** | 0.3145 | n.s. |
| CCL4 | 18.68 ± 1.51 | 22.07 ± 3.68 | 19.51 ± 1.92 | 20.35 ± 2.04 | >0.9999 | n.s. | >0.9999 | n.s. | >0.9999 | n.s. | >0.9999 | n.s. | >0.9999 | n.s. |
| CCL12 | 154.70 ± 7.51 | 144.30 ± 17.41 | 160.40 ± 15.62 | 156.10 ± 19.08 | 0.5574 | n.s. | >0.9999 | n.s. | >0.9999 | n.s. | >0.9999 | n.s. | >0.9999 | n.s. |
| CSF1 | 3806.00 ± 504.60 | 4188.00 ± 440.10 | 2677.00 ± 168.50 | 2986.00 ± 223.20 | 0.992 | n.s. | 0.2657 | n.s. | 0.6056 | n.s. | 0.1519 | n.s. | 0.8384 | n.s. |
| CSF2 | 0.12 ± 0.02 | 0.12 ± 0.03 | 0.13 ± 0.03 | 0.10 ± 0.02 | >0.9999 | n.s. | >0.9999 | n.s. | >0.9999 | n.s. | >0.9999 | n.s. | >0.9999 | n.s. |
| HGF | 1576.00 ± 512.80 | 3372.00 ± 1087.00 | 2419.00 ± 824.80 | 1783.00 ± 616.50 | 0.7278 | n.s. | >0.9999 | n.s. | >0.9999 | n.s. | 0.8989 | n.s. | >0.9999 | n.s. |
| FGF21 | 282.00 ± 50.81 | 293.10 ± 34.20 | 237.10 ± 22.17 | 157.00 ± 27.82 | >0.9999 | n.s. | >0.9999 | n.s. | 0.1684 | n.s. | 0.0515 | n.s. | 0.6857 | n.s. |
| IL-3 | 0.04 ± 0.01 | 0.03 ± 0.01 | 0.04 ± 0.01 | 0.04 ± 0.02 | >0.9999 | n.s. | >0.9999 | n.s. | >0.9999 | n.s. | >0.9999 | n.s. | >0.9999 | n.s. |
| PD-L2 | 2257.00 ± 250.70 | 2215.00 ± 248.10 | 2238.00 ± 164.60 | 2549.00 ± 289.80 | 0.999 | n.s. | 0.999 | n.s. | 0.9307 | n.s. | 0.9251 | n.s. | 0.9307 | n.s. |
| PD-L1 | 65.65 ± 8.88 | 73.40 ± 13.40 | 84.96 ± 19.23 | 87.41 ± 17.95 | >0.9999 | n.s. | >0.9999 | n.s. | >0.9999 | n.s. | >0.9999 | n.s. | >0.9999 | n.s. |
| CTLA4 | 0.41 ± 0.06 | 0.38 ± 0.04 | 0.44 ± 0.06 | 0.45 ± 0.07 | >0.9999 | n.s. | >0.9999 | n.s. | >0.9999 | n.s. | >0.9999 | n.s. | >0.9999 | n.s. |
| IL-7 | 0.48 ± 0.06 | 0.68 ± 0.11 | 0.63 ± 0.09 | 0.72 ± 0.10 | 0.8623 | n.s. | 0.7896 | n.s. | 0.1598 | n.s. | >0.9999 | n.s. | >0.9999 | n.s. |
| IL-2 | 1.22 ± 0.08 | 1.61 ± 0.22 | 2.57 ± 0.75 | 1.37 ± 0.11 | >0.9999 | n.s. | 0.1194 | n.s. | >0.9999 | n.s. | >0.9999 | n.s. | 0.8691 | n.s. |
| IL-5 | 0.38 ± 0.15 | 0.34 ± 0.06 | 0.31 ± 0.04 | 0.40 ± 0.11 | >0.9999 | n.s. | >0.9999 | n.s. | >0.9999 | n.s. | >0.9999 | n.s. | >0.9999 | n.s. |
| IL-33 | 6.46 ± 3.29 | 8.48 ± 5.60 | 11.74 ± 5.72 | 14.74 ± 9.30 | >0.9999 | n.s. | >0.9999 | n.s. | >0.9999 | n.s. | >0.9999 | n.s. | >0.9999 | n.s. |

**Supplemental Table S10** Echocardiographic parameters, presented as mean ± SEM, with p-values for the most relevant comparisons in the *Ttn*$^{+/-}$ cohort. Mice were MCMV-infected/ received PBS at 3 months of age and were analyzed at 1mpi.

| **1mpi** | Group | Non-inf. | Ctr MCMV | *Ttn*$^{+/-}$ MCMV | Main comparisons | | | | | |
|---|---|---|---|---|---|---|---|---|---|---|
| | n | 10 | 9 | 9 | Non-inf. vs. Ctr MCMV | | Non-inf. vs. *Ttn*$^{+/-}$ MCMV | | Ctr MCMV vs. *Ttn*$^{+/-}$ MCMV | |
| | Sex [m/f] | 5/5 | 5/4 | 5/4 | | | | | | |
| Parameter | Unit | Value ± SEM | Value ± SEM | Value ± SEM | p-value | | p-value | | p-value | |
| LVAW;s | mm | 1.86 ± 0.09 | 1.91 ± 0.09 | 1.75 ± 0.06 | 0.9073 | n.s. | 0.6445 | n.s. | 0.4145 | n.s. |
| LVAW;d | mm | 1.16 ± 0.08 | 1.17 ± 0.07 | 1.03 ± 0.06 | >0.9999 | n.s. | 0.5260 | n.s. | 0.3866 | n.s. |
| LVPW;s | mm | 1.98 ± 0.07 | 2.00 ± 0.09 | 1.87 ± 0.07 | 0.9842 | n.s. | 0.5744 | n.s. | 0.4901 | n.s. |
| LVPW;d | mm | 1.26 ± 0.05 | 1.94 ± 0.64 | 1.25 ± 0.05 | 0.7843 | n.s. | >0.9999 | n.s. | 0.7552 | n.s. |
| LVESD | mm | 1.51 ± 0.12 | 1.77 ± 0.14 | 1.90 ± 0.10 | 0.3061 | n.s. | 0.0778 | n.s. | 0.7344 | n.s. |
| LVEDD | mm | 3.49 ± 0.10 | 3.76 ± 0.13 | 3.84 ± 0.09 | 0.2010 | n.s. | 0.0746 | n.s. | 0.8639 | n.s. |
| HR | bpm | 563.34 ± 9.59 | 552.54 ± 10.19 | 557.88 ± 8.08 | 0.6934 | n.s. | 0.9098 | n.s. | 0.9173 | n.s. |
| FS | % | 57.05 ± 2.56 | 53.52 ± 2.76 | 50.82 ± 1.64 | 0.5534 | n.s. | 0.1726 | n.s. | 0.7151 | n.s. |
| SimpV;s | µl | 19.63 ± 1.76 | 21.73 ± 2.62 | 18.46 ± 1.95 | >0.9999 | n.s. | >0.9999 | n.s. | 0.7914 | n.s. |
| SimpV;d | µl | 68.31 ± 5.05 | 73.72 ± 8.65 | 63.24 ± 5.69 | 0.8275 | n.s. | 0.8465 | n.s. | 0.8465 | n.s. |
| SimpFAC | % | 67.55 ± 1.45 | 65.30 ± 1.55 | 66.87 ± 1.84 | 0.5870 | n.s. | 0.9506 | n.s. | 0.7809 | n.s. |
| SimpEF | % | 71.45 ± 1.20 | 70.48 ± 1.47 | 70.96 ± 1.25 | >0.9999 | n.s. | >0.9999 | n.s. | >0.9999 | n.s. |
| SimpCO | ml/min | 25.42 ± 1.74 | 28.77 ± 3.80 | 24.11 ± 2.24 | >0.9999 | n.s. | >0.9999 | n.s. | 0.9898 | n.s. |
| SimpSV | µl | 48.68 ± 3.57 | 51.99 ± 6.48 | 44.78 ± 3.99 | >0.9999 | n.s. | >0.9999 | n.s. | >0.9999 | n.s. |
| IVRT | ms | 19.97 ± 0.45 | 20.25 ± 0.78 | 22.69 ± 1.52 | >0.9999 | n.s. | 0.6184 | n.s. | 0.7168 | n.s. |
| TAPSE | mm | 0.91 ± 0.06 | 0.88 ± 0.06 | 0.87 ± 0.05 | 0.9196 | n.s. | 0.8986 | n.s. | 0.9987 | n.s. |

**Supplemental Table S11** Echocardiographic parameters, presented as mean ± SEM, with p-values for the most relevant comparisons in the *Ttn*$^{+/-}$ cohort. Mice were MCMV-infected/ received PBS at 3 months of age and were analyzed at 3mpi.

| **3mpi** | Group | Non-inf. | Ctr MCMV | *Ttn*$^{+/-}$ MCMV | Main comparisons | | | | | |
|---|---|---|---|---|---|---|---|---|---|---|
| | n | 10 | 9 | 9 | Non-inf. vs. Ctr MCMV | | Non-inf. vs. *Ttn*$^{+/-}$ MCMV | | Ctr MCMV vs. *Ttn*$^{+/-}$ MCMV | |
| | Sex [m/f] | 5/5 | 5/4 | 5/4 | | | | | | |
| Parameter | Unit | Value ± SEM | Value ± SEM | Value ± SEM | p-value | | p-value | | p-value | |
| LVAW;s | mm | 1.78 ± 0.06 | 1.95 ± 0.09 | 1.72 ± 0.06 | 0.2166 | n.s. | 0.8130 | n.s. | 0.0775 | n.s. |
| LVAW;d | mm | 1.10 ± 0.06 | 1.28 ± 0.07 | 1.06 ± 0.05 | 0.1118 | n.s. | 0.8532 | n.s. | 0.0427 | * |
| LVPW;s | mm | 2.01 ± 0.09 | 1.91 ± 0.05 | 1.91 ± 0.04 | 0.5774 | n.s. | 0.5771 | n.s. | >0.9999 | n.s. |
| LVPW;d | mm | 1.37 ± 0.05 | 1.34 ± 0.04 | 1.34 ± 0.05 | 0.8845 | n.s. | 0.8806 | n.s. | >0.9999 | n.s. |
| LVESD | mm | 1.65 ± 0.19 | 1.88 ± 0.19 | 1.89 ± 0.15 | >0.9999 | n.s. | 0.8185 | n.s. | >0.9999 | n.s. |
| LVEDD | mm | 3.54 ± 0.13 | 3.68 ± 0.17 | 3.69 ± 0.11 | 0.7360 | n.s. | 0.7082 | n.s. | 0.9989 | n.s. |
| HR | bpm | 554.61 ± 4.35 | 571.29 ± 6.27 | 560.77 ± 11.29 | 0.1247 | n.s. | 0.9389 | n.s. | 0.8010 | n.s. |
| FS | % | 54.12 ± 3.90 | 50.14 ± 2.93 | 49.15 ± 2.96 | 0.6784 | n.s. | 0.5492 | n.s. | 0.9771 | n.s. |
| SimpV;s | µl | 16.76 ± 2.46 | 20.58 ± 2.53 | 18.55 ± 2.47 | 0.4621 | n.s. | >0.9999 | n.s. | >0.9999 | n.s. |
| SimpV;d | µl | 62.50 ± 5.62 | 72.20 ± 7.76 | 67.28 ± 6.17 | >0.9999 | n.s. | >0.9999 | n.s. | >0.9999 | n.s. |
| SimpFAC | % | 66.94 ± 2.43 | 65.57 ± 2.16 | 67.72 ± 1.11 | 0.8776 | n.s. | 0.9577 | n.s. | 0.7255 | n.s. |
| SimpEF | % | 74.06 ± 1.50 | 71.72 ± 1.40 | 73.11 ± 1.40 | 0.6049 | n.s. | >0.9999 | n.s. | >0.9999 | n.s. |
| SimpCO | ml/min | 26.70 ± 1.57 | 28.26 ± 2.37 | 26.58 ± 2.18 | >0.9999 | n.s. | >0.9999 | n.s. | >0.9999 | n.s. |
| SimpSV | µl | 45.74 ± 3.27 | 51.62 ± 5.51 | 48.73 ± 3.91 | >0.9999 | n.s. | >0.9999 | n.s. | >0.9999 | n.s. |
| IVRT | ms | 20.36 ± 0.52 | 19.31 ± 0.39 | 18.43 ± 0.40 | 0.2496 | n.s. | 0.0131 | * | 0.3915 | n.s. |
| TAPSE | mm | 1.02 ± 0.05 | 0.93 ± 0.06 | 1.08 ± 0.05 | 0.4800 | n.s. | 0.7115 | n.s. | 0.1283 | n.s. |

**Supplemental Table S12** Echocardiographic parameters, presented as mean ± SEM, with p-values for the most relevant comparisons in the *Ttn*$^{+/-}$ cohort. Mice were MCMV-infected/ received PBS at 3 months of age and were analyzed at 6mpi.

| 6mpi | Group | Non-inf. | Ctr MCMV | *Ttn*$^{+/-}$ MCMV | Main comparisons | | | | | |
|---|---|---|---|---|---|---|---|---|---|---|
| | n | 10 | 9 | 8 | Non-inf. vs. Ctr MCMV | | Non-inf. vs. *Ttn*$^{+/-}$ MCMV | | Ctr MCMV vs. *Ttn*$^{+/-}$ MCMV | |
| | Sex [m/f] | 5/5 | 5/4 | 5/3 | | | | | | |
| Parameter | Unit | Value ± SEM | Value ± SEM | Value ± SEM | p-value | | p-value | | p-value | |
| LVAW;s | mm | 1.65 ± 0.07 | 1.81 ± 0.08 | 1.83 ± 0.08 | 0.3199 | n.s. | 0.2508 | n.s. | 0.9781 | n.s. |
| LVAW;d | mm | 1.09 ± 0.04 | 1.20 ± 0.06 | 1.17 ± 0.08 | 0.4528 | n.s. | 0.6504 | n.s. | 0.9567 | n.s. |
| LVPW;s | mm | 1.83 ± 0.07 | 1.98 ± 0.10 | 1.85 ± 0.05 | 0.3284 | n.s. | 0.9640 | n.s. | 0.5073 | n.s. |
| LVPW;d | mm | 1.35 ± 0.05 | 1.45 ± 0.08 | 1.36 ± 0.03 | 0.7151 | n.s. | >0.9999 | n.s. | >0.9999 | n.s. |
| LVESD | mm | 1.87 ± 0.16 | 1.95 ± 0.23 | 1.86 ± 0.17 | 0.9505 | n.s. | 0.9997 | n.s. | 0.9480 | n.s. |
| LVEDD | mm | 3.47 ± 0.13 | 3.63 ± 0.18 | 3.65 ± 0.12 | 0.7226 | n.s. | 0.6607 | n.s. | 0.9917 | n.s. |
| HR | bpm | 570.26 ± 11.98 | 568.79 ± 7.18 | 572.77 ± 8.51 | >0.9999 | n.s. | >0.9999 | n.s. | >0.9999 | n.s. |
| FS | % | 46.98 ± 2.90 | 47.69 ± 3.77 | 49.45 ± 3.64 | >0.9999 | n.s. | >0.9999 | n.s. | >0.9999 | n.s. |
| SimpV;s | µl | 18.14 ± 2.64 | 20.10 ± 3.67 | 17.24 ± 1.60 | >0.9999 | n.s. | >0.9999 | n.s. | >0.9999 | n.s. |
| SimpV;d | µl | 54.39 ± 5.74 | 59.91 ± 7.34 | 58.28 ± 4.19 | 0.7875 | n.s. | 0.9015 | n.s. | 0.9826 | n.s. |
| SimpFAC | % | 56.99 ± 1.51 | 60.45 ± 3.30 | 63.95 ± 3.51 | 0.6273 | n.s. | 0.2106 | n.s. | 0.6714 | n.s. |
| SimpEF | % | 67.85 ± 1.32 | 68.11 ± 2.04 | 70.57 ± 1.33 | 0.9922 | n.s. | 0.4948 | n.s. | 0.5753 | n.s. |
| SimpCO | ml/min | 20.82 ± 1.62 | 21.81 ± 1.81 | 22.79 ± 1.48 | 0.8999 | n.s. | 0.7017 | n.s. | 0.9196 | n.s. |
| SimpSV | µl | 36.25 ± 3.11 | 39.81 ± 3.80 | 41.04 ± 2.85 | 0.7166 | n.s. | 0.5955 | n.s. | 0.9674 | n.s. |
| IVRT | ms | 19.92 ± 0.40 | 18.61 ± 0.63 | 18.02 ± 0.68 | 0.2221 | n.s. | 0.0624 | n.s. | 0.7560 | n.s. |
| TAPSE | mm | 0.88 ± 0.04 | 0.89 ± 0.08 | 0.92 ± 0.04 | 0.9999 | n.s. | 0.8596 | n.s. | 0.9762 | n.s. |

**Supplemental Table S13** Group sizes and within-group p-values for selected echocardiographic parameters over time post-infection in the *Ttn*$^{+/-}$ cohort.

| | Group | Non-inf. | | Ctr MCMV | | *Ttn*$^{+/-}$ MCMV | |
|---|---|---|---|---|---|---|---|
| 1mpi | n | 10 | | 9 | | 9 | |
| | Sex [m/f] | 5/5 | | 5/4 | | 5/4 | |
| 3mpi | n | 10 | | 9 | | 9 | |
| | Sex [m/f] | 5/5 | | 5/4 | | 5/4 | |
| 6mpi | n | 10 | | 9 | | 8 | |
| | Sex [m/f] | 5/5 | | 5/4 | | 5/3 | |
| LVEDD | 1mpi vs. 3mpi | 0.9611 | n.s. | 0.9331 | n.s. | 0.5709 | n.s. |
| | 1mpi vs. 6mpi | 0.9922 | n.s. | 0.8241 | n.s. | 0.4471 | n.s. |
| | 3mpi vs. 6mpi | 0.9207 | n.s. | 0.9688 | n.s. | 0.9687 | n.s. |
| HR | 1mpi vs. 3mpi | >0.9999 | n.s. | 0.2466 | n.s. | 0.9739 | n.s. |
| | 1mpi vs. 6mpi | 0.9656 | n.s. | 0.3437 | n.s. | 0.5252 | n.s. |
| | 3mpi vs. 6mpi | 0.1703 | n.s. | 0.9739 | n.s. | 0.6551 | n.s. |
| FS | 1mpi vs. 3mpi | 0.7925 | n.s. | 0.7353 | n.s. | 0.9040 | n.s. |
| | 1mpi vs. 6mpi | 0.0821 | n.s. | 0.4118 | n.s. | 0.9386 | n.s. |
| | 3mpi vs. 6mpi | 0.2674 | n.s. | 0.8512 | n.s. | 0.9968 | n.s. |
| SimpFAC | 1mpi vs. 3mpi | 0.9691 | n.s. | 0.9967 | n.s. | 0.9689 | n.s. |
| | 1mpi vs. 6mpi | **0.0008** | *** | 0.3557 | n.s. | 0.8426 | n.s. |
| | 3mpi vs. 6mpi | **0.0019** | ** | 0.3184 | n.s. | 0.6785 | n.s. |
| SimpLVEF | 1mpi vs. 3mpi | 0.3750 | n.s. | 0.8578 | n.s. | 0.4768 | n.s. |
| | 1mpi vs. 6mpi | 0.1489 | n.s. | 0.5800 | n.s. | 0.9779 | n.s. |
| | 3mpi vs. 6mpi | **0.0087** | ** | 0.2925 | n.s. | 0.4070 | n.s. |
| SimpCO | 1mpi vs. 3mpi | 0.8524 | n.s. | >0.9999 | n.s. | 0.6634 | n.s. |
| | 1mpi vs. 6mpi | 0.1317 | n.s. | 0.2550 | n.s. | 0.9006 | n.s. |
| | 3mpi vs. 6mpi | **0.0496** | * | 0.1304 | n.s. | 0.4363 | n.s. |
| SimpSV | 1mpi vs. 3mpi | 0.8114 | n.s. | >0.9999 | n.s. | 0.7237 | n.s. |
| | 1mpi vs. 6mpi | **0.0324** | * | 0.1607 | n.s. | 0.7744 | n.s. |
| | 3mpi vs. 6mpi | 0.1337 | n.s. | 0.1399 | n.s. | 0.3544 | n.s. |
| IVRT | 1mpi vs. 3mpi | 0.8239 | n.s. | 0.5549 | n.s. | 0.1028 | n.s. |
| | 1mpi vs. 6mpi | 0.9959 | n.s. | 0.1857 | n.s. | **0.0329** | * |
| | 3mpi vs. 6mpi | 0.7761 | n.s. | 0.7357 | n.s. | >0.9999 | n.s. |
| TAPSE | 1mpi vs. 3mpi | 0.3211 | n.s. | 0.8535 | n.s. | **0.0106** | * |
| | 1mpi vs. 6mpi | 0.9213 | n.s. | 0.9950 | n.s. | 0.7965 | n.s. |
| | 3mpi vs. 6mpi | 0.1899 | n.s. | 0.8980 | n.s. | 0.0860 | n.s. |

**Supplemental Table S14** Echocardiographic parameters (mean ± SEM) and p-values in 12-month-old *Pkp2*$^{+/-}$ and Ctr mice.

| **12mo** | Group | Ctr | *Pkp2*$^{+/-}$ | Comparison | |
|---|---|---|---|---|---|
| | n | 9 | 9 | *Pkp2*$^{+/-}$ vs. Ctr | |
| | Sex [m/f] | 4/5 | 8/1 | | |
| Parameter | Unit | Value ± SEM | Value ± SEM | p-value | |
| Area;s | mm$^2$ | 15.72 ± 1.08 | 22.74 ± 2.49 | 0.0597 | n.s. |
| Area;d | mm$^2$ | 23.97 ± 1.47 | 29.28 ± 2.23 | 0.2224 | n.s. |
| HR | bpm | 477.12 ± 20.82 | 503.95 ± 17.98 | 0.1984 | n.s. |
| FS;long. | % | 11.84 ± 2.19 | 8.24 ± 1.21 | 0.3296 | n.s. |
| V;s | µl | 32.40 ± 3.60 | 62.97 ± 11.73 | 0.0573 | n.s. |
| V;d | µl | 66.34 ± 6.94 | 93.76 ± 11.77 | 0.2224 | n.s. |
| LVEF | % | 51.10 ± 2.57 | 36.66 ± 3.90 | **0.0252** | * |
| CO | ml/min | 16.23 ± 2.02 | 15.70 ± 1.60 | 0.9622 | n.s. |
| SV | µl | 33.94 ± 3.93 | 30.79 ± 2.38 | 0.6312 | n.s. |
| MV E | mm/s | 646.67 ± 35.35 | 655.69 ± 38.02 | 0.8646 | n.s. |
| MV A | mm/s | 400.10 ± 45.19 | 405.71 ± 43.04 | 0.9296 | n.s. |
| MV E/A | a.u. | 1.75 ± 0.20 | 1.67 ± 0.10 | 0.7773 | n.s. |
| LVESD | mm | 3.05 ± 0.19 | 4.04 ± 0.29 | **0.0300** | * |
| LVEDD | mm | 4.20 ± 0.17 | 4.87 ± 0.21 | 0.0783 | n.s. |
| LVPW;s | mm | 1.04 ± 0.07 | 0.91 ± 0.06 | 0.2916 | n.s. |
| LVPW;d | mm | 0.77 ± 0.04 | 0.73 ± 0.04 | 0.4477 | n.s. |
| IVS;s | mm | 1.15 ± 0.09 | 0.89 ± 0.06 | 0.0843 | n.s. |
| IVS;d | mm | 0.87 ± 0.07 | 0.81 ± 0.05 | 0.5671 | n.s. |
| LV mass | mg | 133.35 ± 11.72 | 158.59 ± 14.94 | 0.5859 | n.s. |
| RVOT | mm | 1.53 ± 0.10 | 1.56 ± 0.07 | 0.9941 | n.s. |

**Supplemental Table S15** Echocardiographic parameters (mean ± SEM) and p-values in 15-month-old *Pkp2*$^{+/-}$ and Ctr mice.

| **3mpi (15mo)** | Group | Ctr | *Pkp2*$^{+/-}$ | Comparison | |
|---|---|---|---|---|---|
| | n | 9 | 9 | *Pkp2*$^{+/-}$ vs. Ctr | |
| | Sex [m/f] | 4/5 | 8/1 | | |
| Parameter | Unit | Value ± SEM | Value ± SEM | p-value | |
| Area;s | mm$^2$ | 16.47 ± 0.92 | 24.93 ± 2.55 | **0.0010** | ** |
| Area;d | mm$^2$ | 25.92 ± 1.72 | 31.37 ± 2.09 | 0.0927 | n.s. |
| HR | bpm | 463.37 ± 14.99 | 477.75 ± 22.71 | 0.6146 | n.s. |
| FS;long. | % | 13.79 ± 2.01 | 9.35 ± 1.37 | 0.0828 | n.s. |
| V;s | µl | 34.95 ± 3.53 | 72.32 ± 13.49 | **0.0010** | ** |
| V;d | µl | 74.97 ± 8.91 | 102.10 ± 12.15 | 0.0745 | n.s. |
| LVEF | % | 52.86 ± 1.68 | 31.86 ± 3.74 | **<0.0001** | **** |
| CO | ml/min | 18.39 ± 2.44 | 13.95 ± 1.48 | 0.1311 | n.s. |
| SV | µl | 40.02 ± 5.65 | 29.78 ± 3.14 | 0.1229 | n.s. |
| MV E | mm/s | 559.84 ± 33.45 | 626.32 ± 18.22 | 0.0938 | n.s. |
| MV A | mm/s | 363.41 ± 29.56 | 440.12 ± 28.20 | 0.0834 | n.s. |
| MV E/A | a.u. | 1.57 ± 0.09 | 1.45 ± 0.07 | 0.3169 | n.s. |
| LVESD | mm | 3.23 ± 0.18 | 4.22 ± 0.28 | **0.0050** | ** |
| LVEDD | mm | 4.33 ± 0.18 | 5.01 ± 0.23 | **0.0206** | * |
| LVPW;s | mm | 1.08 ± 0.05 | 0.92 ± 0.06 | 0.0747 | n.s. |
| LVPW;d | mm | 0.80 ± 0.05 | 0.77 ± 0.04 | 0.5654 | n.s. |
| IVS;s | mm | 0.89 ± 0.05 | 0.81 ± 0.05 | 0.2265 | n.s. |
| IVS;d | mm | 0.71 ± 0.05 | 0.71 ± 0.04 | 0.9968 | n.s. |
| LV mass | mg | 124.88 ± 8.29 | 154.72 ± 9.09 | **0.0297** | * |
| RVOT | mm | 1.46 ± 0.06 | 1.54 ± 0.09 | 0.4774 | n.s. |

**Supplemental Table S16** Cytokine concentrations (mean ± SEM) in heart tissue from 9- and 15-month-old *Pkp2*$^{+/-}$ mice.

| | 9mo | 15mo | 9mo | 15mo | Main comparisons | | | | | | | |
|---|---|---|---|---|---|---|---|---|---|---|---|---|
| Group | Ctr | Ctr | *Pkp2*$^{+/-}$ | *Pkp2*$^{+/-}$ | Ctr | | *Pkp2*$^{+/-}$ | | 9mo | | 15mo | |
| n | 10 | 6 | 10 | 10 | 9mo vs. 15mo | | 9mo vs. 15mo | | *Pkp2*$^{+/-}$ vs. Ctr | | *Pkp2*$^{+/-}$ vs. Ctr | |
| Sex [m/f] | 10/0 | 6/0 | 10/0 | 10/0 | | | | | | | | |
| Mediator | Values ± SEM [pg/ml] | | | | p-values | | | | | | | |
| IL-1α | 1.17 ± 0.38 | 0.53 ± 0.19 | 0.84 ± 0.15 | 0.87 ± 0.21 | 0.3132 | n.s. | 0.6842 | n.s. | 0.8534 | n.s. | 0.3132 | n.s. |
| IL-1β | 0.13 ± 0.01 | 0.09 ± 0.02 | 0.20 ± 0.02 | 0.25 ± 0.05 | 0.1866 | n.s. | 0.8534 | n.s. | **0.0187** | * | **0.0110** | * |
| IL-6 | 12.97 ± 2.50 | 13.14 ± 2.97 | 12.55 ± 2.10 | 11.26 ± 1.74 | 0.9578 | n.s. | 0.6842 | n.s. | 0.9705 | n.s. | 0.5663 | n.s. |
| IL-17A | 0.04 ± 0.01 | 0.04 ± 0.01 | 0.04 ± 0.00 | 0.03 ± 0.00 | 0.9578 | n.s. | 0.3616 | n.s. | 0.8534 | n.s. | 0.4393 | n.s. |
| IL-17F | 0.14 ± 0.04 | 0.11 ± 0.03 | 0.13 ± 0.03 | 0.07 ± 0.02 | 0.6281 | n.s. | 0.0672 | n.s. | 0.7164 | n.s. | 0.1747 | n.s. |
| IL-22 | 0.16 ± 0.00 | 0.13 ± 0.02 | 0.15 ± 0.01 | 0.14 ± 0.02 | 0.1596 | n.s. | 0.4981 | n.s. | 0.6305 | n.s. | 0.8865 | n.s. |
| TNFα | 4.23 ± 0.50 | 3.85 ± 0.50 | 3.73 ± 0.38 | 3.46 ± 0.39 | 0.8252 | n.s. | 0.6330 | n.s. | 0.4807 | n.s. | 0.5716 | n.s. |
| IFN-γ | 0.03 ± 0.00 | 0.03 ± 0.01 | 0.02 ± 0.00 | 0.02 ± 0.01 | 0.6087 | n.s. | 0.9246 | n.s. | 0.2158 | n.s. | 0.7179 | n.s. |
| IFN-λ2 | 0.03 ± 0.01 | 0.02 ± 0.00 | 0.02 ± 0.01 | 0.03 ± 0.00 | 0.7922 | n.s. | 0.1893 | n.s. | 0.7308 | n.s. | **0.0264** | * |
| IFN-α2 | 0.03 ± 0.00 | 0.03 ± 0.00 | 0.03 ± 0.00 | 0.03 ± 0.00 | 0.3689 | n.s. | 0.1227 | n.s. | 0.8812 | n.s. | 0.9530 | n.s. |
| IL-4 | 0.17 ± 0.01 | 0.11 ± 0.02 | 0.16 ± 0.01 | 0.10 ± 0.01 | **0.0203** | * | **0.0029** | ** | 0.9023 | n.s. | 0.8201 | n.s. |
| IL-10 | 0.05 ± 0.01 | 0.04 ± 0.01 | 0.05 ± 0.02 | 0.03 ± 0.01 | 0.1695 | n.s. | 0.3924 | n.s. | 0.7386 | n.s. | 0.4496 | n.s. |
| IL-21 | 3.61 ± 0.76 | 2.17 ± 0.68 | 3.24 ± 0.68 | 2.04 ± 0.50 | 0.1898 | n.s. | 0.1684 | n.s. | 0.7215 | n.s. | 0.9578 | n.s. |
| IL-27 | 2.16 ± 0.48 | 2.93 ± 0.79 | 2.17 ± 0.41 | 1.82 ± 0.38 | 0.6828 | n.s. | 0.5054 | n.s. | 0.9591 | n.s. | 0.1804 | n.s. |
| CXCL1 | 0.46 ± 0.05 | 0.44 ± 0.04 | 0.62 ± 0.04 | 0.75 ± 0.13 | 0.7857 | n.s. | 0.9118 | n.s. | **0.0224** | * | **0.0420** | * |
| CXCL9 | 39.07 ± 4.22 | 35.85 ± 4.63 | 30.99 ± 3.39 | 27.42 ± 3.27 | 0.6305 | n.s. | 0.4583 | n.s. | 0.1532 | n.s. | 0.1498 | n.s. |
| CXCL11 | 0.58 ± 0.13 | 0.61 ± 0.18 | 0.62 ± 0.13 | 0.45 ± 0.08 | 0.9578 | n.s. | 0.3008 | n.s. | 0.9705 | n.s. | 0.3921 | n.s. |
| IL-16 | 1420.84 ± 91.08 | 1169.39 ± 132.07 | 1646.48 ± 106.31 | 1623.18 ± 100.09 | 0.1282 | n.s. | 0.8750 | n.s. | 0.1244 | n.s. | 0.0934 | n.s. |
| CCL11 | 7.20 ± 0.98 | 4.02 ± 0.66 | 7.31 ± 0.68 | 4.79 ± 0.54 | **0.0366** | * | **0.0095** | ** | 0.9273 | n.s. | 0.3874 | n.s. |
| CCL2 | 34.48 ± 1.98 | 34.66 ± 6.18 | 80.80 ± 7.38 | 83.97 ± 11.99 | 0.7128 | n.s. | 0.7394 | n.s. | **<0.0001** | **** | **0.0017** | ** |
| CCL5 | 9.57 ± 0.83 | 9.94 ± 1.19 | 10.03 ± 0.92 | 12.61 ± 0.95 | 0.8024 | n.s. | 0.0666 | n.s. | 0.7131 | n.s. | 0.1173 | n.s. |
| CCL22 | 2.71 ± 0.27 | 3.06 ± 0.70 | 6.05 ± 0.78 | 6.55 ± 0.91 | 0.9578 | n.s. | 0.6796 | n.s. | **0.0003** | *** | **0.0179** | * |
| CCL4 | 0.85 ± 0.08 | 0.90 ± 0.14 | 1.18 ± 0.12 | 1.48 ± 0.24 | 0.6354 | n.s. | 0.2914 | n.s. | **0.0318** | * | 0.1084 | n.s. |
| CCL12 | 11.49 ± 1.66 | 10.99 ± 1.93 | 17.21 ± 3.35 | 26.02 ± 5.37 | 0.8489 | n.s. | 0.2176 | n.s. | 0.1500 | n.s. | **0.0160** | * |
| CSF1 | 50.89 ± 7.21 | 36.85 ± 8.45 | 45.59 ± 4.60 | 41.89 ± 3.68 | 0.2382 | n.s. | 0.5379 | n.s. | 0.5429 | n.s. | 0.5389 | n.s. |
| CSF2 | 0.21 ± 0.03 | 0.19 ± 0.04 | 0.19 ± 0.02 | 0.15 ± 0.02 | 0.6908 | n.s. | 0.2162 | n.s. | 0.6842 | n.s. | 0.3899 | n.s. |
| HGF | 27.71 ± 4.83 | 29.15 ± 2.97 | 30.98 ± 4.21 | 37.59 ± 4.21 | 0.8332 | n.s. | 0.2814 | n.s. | 0.6163 | n.s. | 0.1776 | n.s. |
| FGF21 | 0.63 ± 0.07 | 0.80 ± 0.23 | 1.79 ± 0.36 | 3.73 ± 0.76 | >0.9999 | n.s. | **0.0388** | * | **0.0003** | *** | **0.0039** | ** |
| IL-3 | 0.04 ± 0.01 | 0.05 ± 0.01 | 0.04 ± 0.01 | 0.03 ± 0.01 | 0.7679 | n.s. | 0.4668 | n.s. | 0.9118 | n.s. | 0.2024 | n.s. |
| PD-L2 | 3.04 ± 0.36 | 4.36 ± 0.98 | 5.11 ± 0.50 | 5.35 ± 0.75 | 0.3132 | n.s. | 0.7995 | n.s. | **0.0035** | ** | 0.4361 | n.s. |
| PD-L1 | 343.81 ± 21.85 | 290.92 ± 23.96 | 387.24 ± 67.83 | 286.94 ± 24.02 | 0.1406 | n.s. | 0.1903 | n.s. | 0.7394 | n.s. | 0.9142 | n.s. |
| CTLA4 | 0.13 ± 0.03 | 0.14 ± 0.03 | 0.14 ± 0.02 | 0.10 ± 0.02 | 0.8518 | n.s. | 0.3527 | n.s. | 0.8286 | n.s. | 0.3187 | n.s. |
| IL-7 | 4.35 ± 0.78 | 4.38 ± 1.13 | 4.19 ± 0.65 | 3.07 ± 0.55 | 0.9530 | n.s. | 0.2428 | n.s. | 0.8534 | n.s. | 0.2594 | n.s. |
| IL-2 | 0.49 ± 0.06 | 0.43 ± 0.09 | 0.45 ± 0.05 | 0.40 ± 0.04 | 0.5775 | n.s. | 0.4637 | n.s. | 0.5787 | n.s. | 0.6948 | n.s. |
| IL-5 | 0.06 ± 0.01 | 0.05 ± 0.01 | 0.05 ± 0.01 | 0.04 ± 0.01 | 0.6730 | n.s. | 0.1910 | n.s. | 0.7394 | n.s. | 0.3721 | n.s. |
| IL-33 | 21.10 ± 2.75 | 26.11 ± 4.45 | 24.20 ± 2.67 | 41.03 ± 6.34 | 0.3268 | n.s. | **0.0433** | * | 0.4297 | n.s. | 0.1182 | n.s. |

**Supplemental Table S17** Cytokine concentrations (mean ± SEM) in serum from 9- and 15-month-old *Pkp2*$^{+/-}$ mice.

| | **9mo** | **15mo** | **9mo** | **15mo** | Main comparisons | | | | | | | |
|---|---|---|---|---|---|---|---|---|---|---|---|---|
| Group | Ctr | Ctr | *Pkp2*$^{+/-}$ | *Pkp2*$^{+/-}$ | Ctr | | *Pkp2*$^{+/-}$ | | 9mo | | 15mo | |
| n | 10 | 6 | 10 | 10 | 9mo vs. 15mo | | 9mo vs. 15mo | | *Pkp2*$^{+/-}$ vs. Ctr | | *Pkp2*$^{+/-}$ vs. Ctr | |
| Sex [m/f] | 10/0 | 6/0 | 10/0 | 10/0 | | | | | | | | |
| Mediator | Values ± SEM [pg/ml] | | | | p-values | | | | | | | |
| IL-1α | 52.20 ± 27.21 | 35.48 ± 22.89 | 51.42 ± 17.43 | 30.50 ± 7.57 | 0.4278 | n.s. | 0.6842 | n.s. | 0.4813 | n.s. | 0.1471 | n.s. |
| IL-1β | 0.88 ± 0.29 | 0.59 ± 0.17 | 1.13 ± 0.31 | 0.61 ± 0.09 | 0.9578 | n.s. | 0.1377 | n.s. | 0.6305 | n.s. | 0.7128 | n.s. |
| IL-6 | 4.61 ± 0.87 | 12.20 ± 3.01 | 5.36 ± 1.59 | 8.21 ± 2.14 | 0.0525 | n.s. | 0.1903 | n.s. | 0.6836 | n.s. | 0.2868 | n.s. |
| IL-17A | 2.12 ± 0.66 | 4.33 ± 0.80 | 1.75 ± 0.20 | 4.64 ± 1.89 | **0.0160** | * | **0.0068** | ** | 0.7959 | n.s. | 0.1806 | n.s. |
| IL-17F | 6.02 ± 0.97 | 5.23 ± 0.84 | 6.25 ± 1.44 | 7.20 ± 1.54 | 0.5848 | n.s. | 0.4359 | n.s. | 0.8534 | n.s. | 0.4278 | n.s. |
| IL-22 | 8.10 ± 1.51 | 16.02 ± 3.09 | 10.50 ± 2.09 | 14.56 ± 3.95 | **0.0213** | * | 0.6842 | n.s. | 0.4813 | n.s. | 0.5622 | n.s. |
| TNFα | 6.44 ± 0.47 | 8.01 ± 0.86 | 5.59 ± 1.17 | 6.41 ± 0.84 | 0.1000 | n.s. | 0.5742 | n.s. | 0.5110 | n.s. | 0.2281 | n.s. |
| IFN-γ | 0.57 ± 0.08 | 0.86 ± 0.08 | 0.66 ± 0.10 | 0.62 ± 0.05 | **0.0317** | * | 0.7959 | n.s. | 0.5787 | n.s. | **0.0256** | * |
| IFN-λ2 | 0.26 ± 0.02 | 0.46 ± 0.14 | 0.33 ± 0.07 | 0.30 ± 0.06 | 0.0727 | n.s. | 0.7394 | n.s. | 0.4359 | n.s. | 0.2198 | n.s. |
| IFN-α2 | 0.08 ± 0.01 | 0.12 ± 0.02 | 0.07 ± 0.01 | 0.07 ± 0.01 | 0.1401 | n.s. | 0.8108 | n.s. | 0.3454 | n.s. | **0.0325** | * |
| IL-4 | 0.17 ± 0.03 | 0.49 ± 0.21 | 0.20 ± 0.04 | 0.15 ± 0.02 | 0.1806 | n.s. | >0.9999 | n.s. | 0.9118 | n.s. | 0.1806 | n.s. |
| IL-10 | 4.79 ± 0.61 | 7.55 ± 1.62 | 3.15 ± 0.45 | 4.51 ± 0.60 | 0.0934 | n.s. | **0.0288** | * | **0.0115** | * | 0.0727 | n.s. |
| IL-21 | 0.93 ± 0.26 | 0.22 ± 0.09 | 1.08 ± 0.22 | 0.32 ± 0.13 | 0.1000 | n.s. | **0.0113** | * | 0.6533 | n.s. | 0.7758 | n.s. |
| IL-27 | 2.74 ± 0.29 | 3.37 ± 0.71 | 3.31 ± 1.23 | 2.69 ± 0.37 | 0.3543 | n.s. | 0.3930 | n.s. | 0.1431 | n.s. | 0.3668 | n.s. |
| CXCL1 | 18.73 ± 1.57 | 19.82 ± 2.66 | 27.19 ± 4.97 | 18.79 ± 1.91 | 0.7091 | n.s. | 0.0892 | n.s. | 0.1230 | n.s. | 0.7527 | n.s. |
| CXCL9 | 39.07 ± 4.22 | 163.72 ± 17.55 | 30.99 ± 3.39 | 223.03 ± 71.74 | **<0.0001** | **** | **<0.0001** | **** | 0.1532 | n.s. | >0.9999 | n.s. |
| CXCL11 | 0.05 ± 0.01 | 0.09 ± 0.01 | 0.07 ± 0.01 | 0.11 ± 0.02 | **0.0191** | * | 0.1447 | n.s. | 0.3313 | n.s. | 0.5343 | n.s. |
| IL-16 | 2596.37 ± 326.41 | 2192.35 ± 163.49 | 2681.46 ± 238.18 | 2604.18 ± 167.86 | 0.4278 | n.s. | 0.7939 | n.s. | 0.6842 | n.s. | 0.1247 | n.s. |
| CCL11 | 269.43 ± 23.98 | 211.39 ± 28.53 | 327.50 ± 34.69 | 225.54 ± 19.09 | 0.1498 | n.s. | **0.0191** | * | 0.1854 | n.s. | 0.6749 | n.s. |
| CCL2 | 399.18 ± 42.69 | 526.43 ± 97.26 | 459.22 ± 66.91 | 389.91 ± 33.89 | 0.1471 | n.s. | 0.8534 | n.s. | >0.9999 | n.s. | 0.2315 | n.s. |
| CCL5 | 48.27 ± 6.35 | 37.09 ± 12.95 | 78.89 ± 11.12 | 48.20 ± 5.40 | 0.0727 | n.s. | **0.0355** | * | **0.0232** | * | 0.0727 | n.s. |
| CCL22 | 342.21 ± 37.83 | 277.00 ± 57.23 | 473.62 ± 51.68 | 292.08 ± 31.29 | 0.3383 | n.s. | **0.0076** | ** | 0.0550 | n.s. | 0.8038 | n.s. |
| CCL4 | 18.68 ± 1.51 | 19.92 ± 1.23 | 22.07 ± 3.68 | 19.73 ± 1.54 | 0.5773 | n.s. | 0.4813 | n.s. | 0.9705 | n.s. | 0.9331 | n.s. |
| CCL12 | 154.67 ± 7.51 | 213.18 ± 39.20 | 144.33 ± 17.41 | 162.03 ± 11.44 | 0.2198 | n.s. | 0.0753 | n.s. | 0.0524 | n.s. | 0.2198 | n.s. |
| CSF1 | 3806.01 ± 504.58 | 3380.16 ± 327.41 | 4187.93 ± 440.13 | 3087.68 ± 206.13 | 0.5557 | n.s. | **0.0417** | * | 0.5755 | n.s. | 0.4378 | n.s. |
| CSF2 | 0.12 ± 0.02 | 0.28 ± 0.05 | 0.12 ± 0.03 | 0.17 ± 0.02 | **0.0277** | * | 0.1230 | n.s. | 0.7959 | n.s. | 0.0762 | n.s. |
| HGF | 1576.43 ± 512.81 | 968.41 ± 255.46 | 3371.74 ± 1087.45 | 1426.97 ± 539.68 | 0.4278 | n.s. | **0.0433** | * | 0.1051 | n.s. | 0.7128 | n.s. |
| FGF21 | 282.03 ± 50.81 | 149.24 ± 37.99 | 293.13 ± 34.20 | 269.24 ± 26.17 | 0.0882 | n.s. | 0.5859 | n.s. | 0.8582 | n.s. | **0.0178** | * |
| IL-3 | 0.04 ± 0.01 | 0.22 ± 0.07 | 0.03 ± 0.01 | 0.07 ± 0.03 | **0.0420** | * | 0.3884 | n.s. | 0.4559 | n.s. | 0.0829 | n.s. |
| PD-L2 | 2257.12 ± 250.69 | 5853.99 ± 1009.31 | 2215.36 ± 248.13 | 3662.35 ± 597.04 | **0.0150** | * | **0.0052** | ** | 0.9071 | n.s. | **0.0312** | * |
| PD-L1 | 65.65 ± 8.88 | 94.99 ± 24.70 | 73.40 ± 13.40 | 74.76 ± 5.03 | 0.0727 | n.s. | 0.3150 | n.s. | >0.9999 | n.s. | 0.9578 | n.s. |
| CTLA4 | 0.41 ± 0.06 | 0.67 ± 0.13 | 0.38 ± 0.04 | 0.48 ± 0.06 | 0.0934 | n.s. | 0.1694 | n.s. | >0.9999 | n.s. | 0.1641 | n.s. |
| IL-7 | 0.48 ± 0.06 | 1.25 ± 0.28 | 0.68 ± 0.11 | 0.72 ± 0.08 | **0.0030** | ** | 0.3527 | n.s. | 0.1903 | n.s. | 0.0934 | n.s. |
| IL-2 | 1.22 ± 0.08 | 1.46 ± 0.08 | 1.61 ± 0.22 | 1.65 ± 0.14 | 0.0549 | n.s. | 0.8709 | n.s. | 0.2475 | n.s. | 0.3418 | n.s. |
| IL-5 | 0.38 ± 0.15 | 0.79 ± 0.30 | 0.34 ± 0.06 | 0.18 ± 0.01 | 0.0727 | n.s. | **0.0347** | * | 0.5288 | n.s. | **0.0030** | ** |
| IL-33 | 6.46 ± 3.29 | 1.60 ± 0.41 | 8.48 ± 5.60 | 3.18 ± 1.43 | 0.5622 | n.s. | 0.9705 | n.s. | 0.7394 | n.s. | 0.7128 | n.s. |
